# High-throughput thermodynamic and kinetic measurements of transcription factor/DNA mutations reveal how conformational heterogeneity can shape motif selectivity

**DOI:** 10.1101/2023.11.13.566946

**Authors:** R. Hastings, A. Aditham, N. DelRosso, P. Suzuki, P.M. Fordyce

## Abstract

Transcription factors (TFs) bind DNA sequences with a range of affinities, yet the mechanisms determining energetic differences between high- and low-affinity sequences (‘selectivity’) remain poorly understood. Here, we investigated two basic helix-loop-helix TFs, MAX *(H. sapiens)* and Pho4 *(S. cerevisiae)*, that bind the same high-affinity sequence with highly similar nucleotide-contacting residues and bound structures but are differentially selective for non-cognate sequences. By measuring >1700 *K*_d_s and >500 rate constants for Pho4 and MAX mutant libraries binding multiple DNA sequences and comparing these measurements with thermodynamic and kinetic models, we identify the biophysical mechanisms by which changes to TF sequence alter both bound and unbound conformational ensembles to shape specificity landscapes. These results highlight the importance of conformational heterogeneity in determining sequence specificity and selectivity and can guide future efforts to engineer nucleic acid-binding proteins with enhanced selectivity.

---

Transcription factors (TFs) specifically bind regulatory DNA sequences in the genome to control gene expression. Many prior efforts have uncovered the “sequence specificity” (the motif bound with the highest affinity)^1,2^ of thousands of TFs across multiple organisms, with preferences often represented as mononucleotide or dinucleotide models^3–8^. This information, combined with structural models, has fueled progress in designing^9,10^ and predicting^11–15^ sequence-specific nucleic acid binders. However, while motif representations effectively represent “sequence specificity”, they often fail to capture other important features of TF-DNA binding, including absolute binding affinities and preferences for sequences that fall outside of the highest affinity set^16^.

These weaknesses impact our ability to understand natural TF-DNA interactions, where selective pressures on protein and DNA sequence dictate and tune absolute interaction affinities^17^. TF paralogs with identical sequence specificity but different absolute affinities for that sequence play non-interchangeable roles in development^18^. In addition, many closely related TFs that bind the same high-affinity motifs bind alternate low-affinity sites with different sequences and/or absolute affinities^16,19–22^. Underscoring the importance of these interactions, low-affinity DNA sites are conserved and important for gene regulation^21,23,24^; many enhancers are evolved for suboptimal affinities such that increasing affinity is pathogenic or disrupts specific and properly timed gene expression^9–14^.

Focusing on highest affinity sequences alone also hinders the ability to engineer new TF-DNA interactions. Many bioengineering^9^ and gene therapy^30^ objectives require highly specific binding, yet mitigating off-target events remains challenging^31^. While previous attempts to engineer specificity have focused on engineering the lowest energy bound state^32,33^, this structure-driven approach is complicated by the structural plasticity of TFs, which are often enriched in intrinsically disordered sequences^34^, only fold when bound to DNA^35–37^, and adopt different conformations to facilitate binding to different DNA sequences^38–40^. Therefore, realizing the goal of designing binders with both user-specified “sequence specificity” and high “sequence selectivity” – the magnitude of the energetic difference between preferred- and non-preferred sequences – requires new data that can systematically vary both protein and DNA sequence across a range of affinities and map the full DNA binding affinity landscape.

Towards this goal, we systematically investigated how TF sequence shapes DNA binding landscapes by profiling binding for two bHLH TFs: *H. sapiens* Myc-associated factor X (MAX) and *S. cerevisiae* Pho4. While both TFs recognize the same CACGTG E-box motif with nearly identical bound structures and are largely unstructured in the absence of DNA^35,36^, Pho4 is highly selective for this motif while MAX is more promiscuous^41^. Using a recently developed technique for high-throughput microfluidic characterization of TF variant binding (STAMMP, for <u>S</u>imultaneous <u>T</u>ranscription <u>F</u>actor <u>A</u>ffinity <u>M</u>apping via <u>M</u>icrofluidic <u>P</u>rotein arrays)^42^ which can systematically characterize DNA binding for hundreds of TF mutants in parallel, we interrogated how 240 single amino acid substitutions impact MAX DNA binding affinities (*K*_d_s) and kinetics (*k*_off_s) for 7 motif-variant DNA sequences.

Comparisons of the 1,700+ novel *K*_d_s measured here for MAX mutants with data previously collected for Pho4^42^, in concert with thermodynamic and kinetic modeling, revealed that these two TFs differ in “sequence selectivity” due to increased conformational heterogeneity in MAX, which can partition between selective and promiscuous binding conformations. Within these 1700+ novel *K*_d_s, we identified 22 MAX mutations that increased binding “selectivity”, none of which make base-specific DNA contacts in previous structures; additional measurements of 500+ binding rate constants suggest that these mutations increase “selectivity” through perturbations to both bound and unbound states within this heterogeneous conformational ensemble. These results establish the utility of systematic and high-throughput thermodynamic and kinetic measurements of specificity, selectivity, and affinity in the context of combinatorial protein/ligand mutations to detect heterogeneous binding modes, consideration of which will be necessary to engineer highly specific and selective binders.

## Results

### MAX and Pho4 are ideal model systems for investigating how TF sequences encode DNA specificity

The DNA-binding domains (DBDs) of MAX and Pho4 are disordered in solution, with their DNA-contacting regions folding only upon recognition of a DNA binding site to assume highly similar structural conformations (RMSD = 1.519 Å, **Fig. 1A**)^43,44^. Both TFs possess similar domain architectures comprised of a DNA-contacting basic region followed by two helices separated by a flexible loop (**Figs. 1A**,**B**), make identical base contacts via identical nucleotide-contacting residues (**Fig. 1C**), and preferentially bind the same cognate CACGTG E-box site (**Fig. 1D**) as dimers. Despite these similarities, prior measurements of WT Pho4 and MAX affinities for a library of 256 DNA sequences comprised of mutations within an E-box half site revealed differences in their binding energy landscapes (**Fig. 1E**)^41^. Single-nucleotide mutations to the cognate sequence led to 6-fold higher reductions in binding affinity for Pho4 than for MAX (174-fold versus 28-fold), with Pho4 binding CACGTG more tightly and mutant motifs more weakly^41^ (see **Methods**). This difference in “sequence selectivity” can be conceptualized using free energy diagrams in which Pho4 binds its cognate E-box with a deeper energetic well (**Fig. 1E, F**). These differences in binding landscapes despite nearly identical DNA-binding interfaces motivated us to probe how variation in non-contacting residues (**Fig. S1)** can shape binding affinity landscapes.

**Figure 1:**
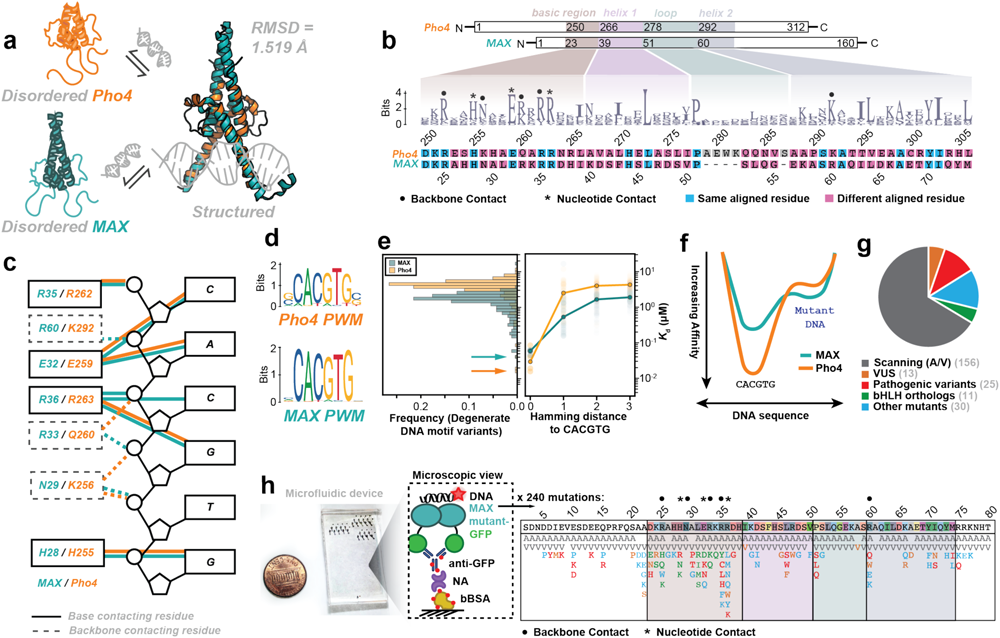
Amino acid sequence, structure, and DNA specificity of MAX and Pho4. **(A)** Schematic of folding-and-binding pathway and structural alignment for Pho4 (orange, PDB: 1A0A) and MAX (teal, PDB: 1HLO). **(B)** Domain architectures and sequence alignment for MAX and Pho4 DNA binding domains alongside conservation across bHLH TFs. **(C)** Crystallographic contacts between the CACGTG cognate E-box and TFs MAX (teal) and Pho4 (orange). **(D)** PWMs for MAX (JASPAR MA0058.3) and Pho4 (JASPAR MA0357.1). **(E)** Distribution of binding affinities for all degenerate E-box motif variants^41^ with most tightly bound sequences annotated (*left*); median affinity as a function of Hamming distance away from the CACGTG cognate motif (*right*). **(F)** Cartoon illustrating differential selectivity. **(G)** Classification of MAX mutations in this study. **(H)** Microfluidic device and zoomed-in view of surface-immobilized TFs (*left*) along with location and identity of MAX mutations studied here (*right*).

### STAMMP enables DNA-binding measurements for hundreds of MAX mutations

Using our microfluidic platform (STAMMP, **Figs. S2-S3**), we previously quantified impacts of 210 single amino acid substitutions within Pho4 on binding to 9 oligonucleotides (>1,800 *K*_d_s in total)^42^. Here, we apply STAMMP to investigate how 240 single amino acid substitutions in and around the MAX DBD **(Fig. S4)** impact DNA binding affinity, specificity, and kinetics. This mutant library included 156 alanine and valine scanning mutants to probe protein sequence determinants of specific DNA motif recognition, 10 mutations substituting orthologous amino acids present in other bHLHs^45^ to probe how evolutionary differences alter landscapes, 30 mutants hypothesized to alter helicity and charge to probe biophysical mechanisms contributing to recognition^46,47^, and 38 mutations from human allelic variants that were catalogued as pathogenic mutations or variants of unknown significance (VUS)^48,49^ (**Fig. 1G-H, Table S1**).

### Many substitutions throughout the MAX DNA-binding domain statistically significantly alter DNA binding

To identify MAX residues involved in DNA recognition, we first measured impacts of each mutation on binding to the cognate 5’-CACGTG-3’ motif (**Fig. 2**). Of 240 mutants, 237 expressed consistently across technical replicates (**Table S2, Fig. S5)**. Measured *K*_d_s fit from processed data (see **Methods, Fig. S6-7, Table S3**) were highly consistent across experimental replicates (RMSE < 0.3 kcal/mol over a 3.5 kcal/mol dynamic range), including for MAX mutants that increased binding affinity **(Figs. 2A, S8)**, and independent of immobilized TF-eGFP concentrations (as expected for TF concentrations well below measured *K*_d_s) **(Fig. S9)**. Subsequent analyses aggregated measurements across all replicates **(Fig. S10)** for each variant to report median *K*_d_ or ΔΔG value (±SEM).

**Figure 2:**
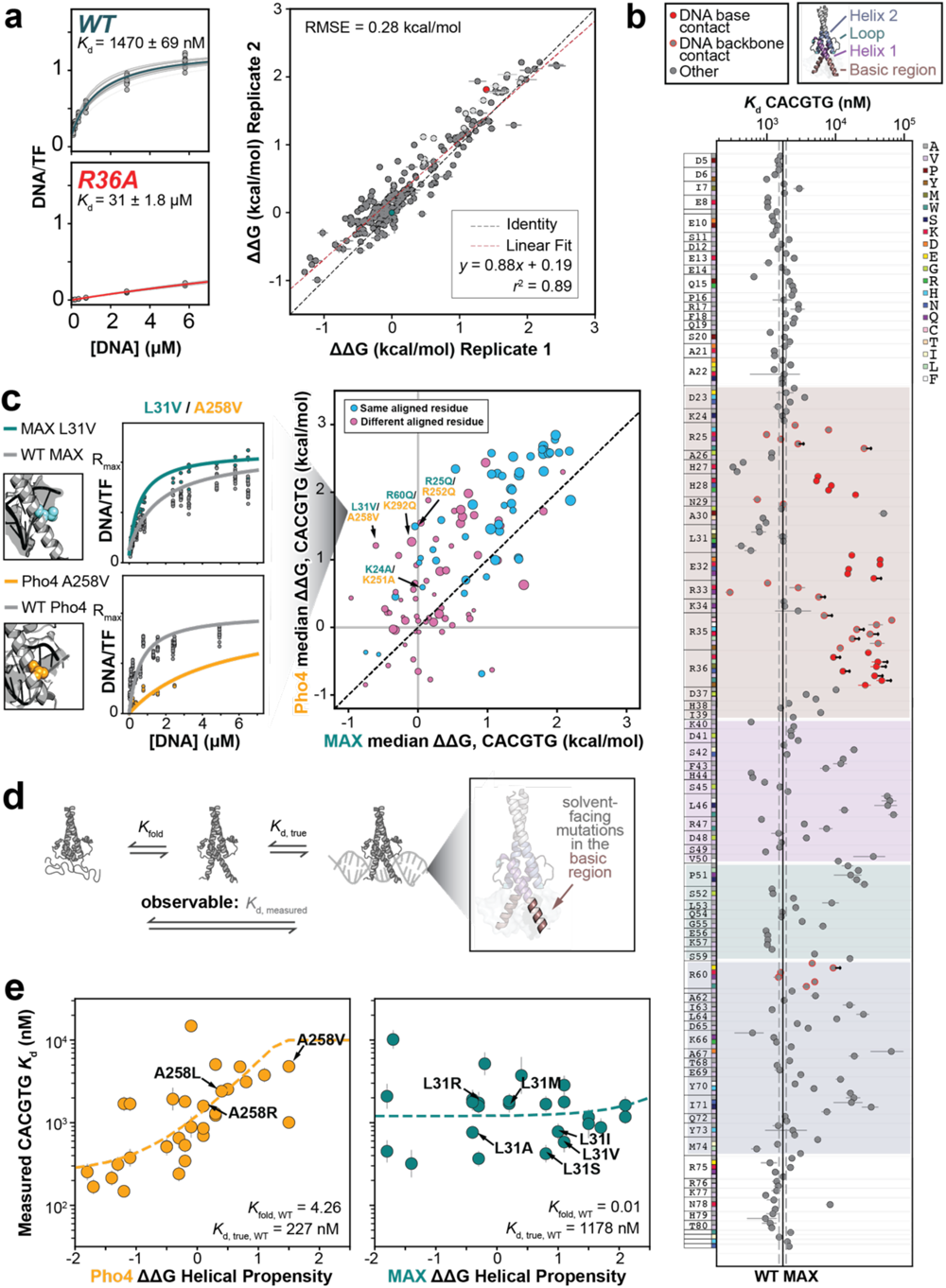
MAX and Pho4 differ in folding-and-binding to CACGTG. **(A)** Sample binding isotherms for WT (teal) and R36A (red) MAX variants binding to cognate DNA (*left*) and reproducibility of ΔΔG measurements across two technical replicates (*right*). Light grey markers indicate mutants un-resolvable from background binding for which reported *K*_d_s represent a lower limit. **(B)** Affinities for MAX mutants binding CACGTG (median ± SEM). Red markers denote mutations to DNA-contacting residues, grey markers with red outlines denote mutations to phosphate backbone-contacting residues, and arrows denote *K*_d_ limits. **(C)** Binding isotherms for WT MAX, MAX L31V, WT Pho4, and Pho4 A258V (*left*) and comparison of ΔΔG measurements for aligned substitutions to MAX and Pho4 (*right*); marker size indicates residue conservation across the bHLH family. **(D)** Thermodynamic model for a three-state system such that observable *K*_d_ depends on the folding equilibrium (*K*_fold_) and true binding affinity (*K*_d_). **(E)** Measured change in cognate affinity (ΔΔG, median ± SEM) versus changes in helical propensity^53^ for mutations to non-DNA contacting basic region residues in MAX (teal) and Pho4 (orange); dashed line indicates fitted thermodynamic model with indicated fitted values of *K*_fold_ and *K*_d_.

Overall, 112 mutants significantly altered the affinity for the consensus motif relative to WT (Bonferroni-corrected *p* < 0.05). Many mutations strongly reduced DNA binding, including substitutions at conserved nucleotide-contacting residues H28, E32, R35, and R36^44^ **(Figs. 2B**). Many significant pathogenic (18/31) and some (I71N and R47W) VUS reduced binding as well (**Fig S11**). Allelic variants imparted some of the largest changes in DNA binding affinity in the library, with significant mutations at the dimerization interface either decreasing (*e*.*g*. L46, A67, and I71) or enhancing binding (*e*.*g*. M74L^50^) (**Fig. S11**). Similarly, mutations to crystal structure-predicted phosphoryl oxygen backbone-contacting residues increased or decreased affinity depending on the substituted residue **(Fig. 2B)**. Nevertheless, 12 of the 29 significant mutations that enhanced binding occurred at non-contacting, solvent-facing residues in the basic region, confirming that TF regions well outside of known interfaces can have substantial impacts on binding^42^.

### Comparable substitutions in Pho4 decrease binding affinity to a greater degree

Just as the magnitude of changes in affinity upon DNA mutation was smaller for MAX than for Pho4, comparable mutations at corresponding residue positions also had smaller impacts in MAX (Wilcoxon signed-rank test *p=10*^*-9*^) (**Figs. 2C, 1E**). In particular, substitutions altering charged contacts with the DNA phosphate backbone yielded substantially larger reductions in binding affinity for Pho4 (Pho4 R252Q/MAX R25Q, Pho4 K292A/MAX R60A) (**Figs. 2C, S12**). This trend also held for mutations to residues that do not contact bound DNA in published structures (Pho4 K251/MAX K24), where some analogous substitutions even increased binding affinity for MAX (L31V) while reducing it for Pho4 (A258V) (**Figs. 2C, S12**). These differences in ‘mutational sensitivity’ support prior observations^51^ that the strength of otherwise similar molecular contacts is contextual and that only certain protein homologs can support ‘rheostat’ positions^52^.

### MAX and Pho4 differ in folding-and-binding transition, suggesting MAX adopts multiple conformations

For TFs that are unstructured in the absence of DNA, altering helical propensity^53^ can commensurately change binding affinity by modulating folding entropic penalties^32,33,42,54^. Thus, if mutations to comparable solvent-facing residues have different impacts on measured binding affinity for CACGTG in Pho4 and MAX, it could indicate differing folding-and-binding transitions.

To better understand how changes in a coupled folding and binding pathway impact measured *K*_d_s, we modeled equilibrium bHLH TF/DNA binding as a system in which: (1) unbound TFs can be either unfolded or helical, (2) only the helical form can bind DNA, and (3) TF mutations alter the folding equilibrium (**Figs. 2D, S13; Methods**). The observed DNA concentration at which half of the TF population is DNA-bound (*K*_d,apparent_) can then be expressed as a function of a *K*_d_ for the interaction between folded TF and free DNA, a folding equilibrium constant *K*_fold_ (describing the partitioning between folded and unfolded conformations of the unbound TF), and the total concentration of free DNA and protein. Simulated binding isotherms for many mutants that alter folding revealed that the magnitude by which mutations shift *K*_d,apparent_ depends on the WT value of *K*_fold_ (**Figs. S13, S14**), and provides a thermodynamic model framework to fit *K*_d, WT_ and *K*_fold, WT_ from measured affinities of non-contacting mutations.

For Pho4, mutations that alter helical propensity concomitantly shift apparent affinities (*K*_d,apparent_) (*r*^*2*^ = 0.55), consistent with a 3-state model in which WT Pho4’s basic region is significantly disordered (∼81%) in the unbound state (**Methods, Fig 2E, S15**) and in agreement with NMR data^35^ . By contrast, mutations that alter the predicted helicity of the MAX basic region have little impact on *K*_d,apparent_ (*r*^*2*^ = 0.20). For this observation to be consistent with a 3-state model, MAX would have to be primarily helical in solution (∼1% disordered), at odds with documented structural disorder in this region^36^ and the observation that many solvent-facing mutations strongly modulate binding affinity (**Methods, Fig 2E, S14-15**). Instead, these results suggest a more complex model in which MAX can adopt multiple conformations with different intrinsic affinities for DNA. In this model, rather than altering the propensity to fold into a *single* conformation, non-contacting mutations impact measured affinity by altering partitioning between *multiple* conformations. This hypothesis is consistent with previously identified bound state conformational heterogeneity within the MAX basic region^55^.

### Mutations that alter dimerization modulate MAX binding affinity independent of DNA sequence

To understand how MAX mutations alter low affinity binding and sequence selectivity within the MAX “DNA-binding landscape”, we also measured binding to 5 low-affinity sequences containing mutations within core nucleotide positions in the E-box consensus (**A**ACGTG, C**G**CGTG, CA**T**GTG, CACG**C**G, and CACGT**T**) **(Figs. 3A-B, Table S4)**. Across all replicates, expression **(Figs. S16-20)** and *K*_d_ measurements remained highly reproducible **(Figs. S21–S25**) and affinities were independent of expression **(Fig. S26)**. Over a median of 8-12 *K*_d_ measurements per variant **(Fig. S27)**, single nucleotide substitutions within the consensus E-box motif reduced WT MAX binding by 2 to 5-fold, with the **A**ACGTG mutation being most deleterious **(Figs. 3B, S28)**. MAX allelic variants yield diverse biochemical effects on the DNA-binding landscape, summarized in **Table S5**.

**Figure 3:**
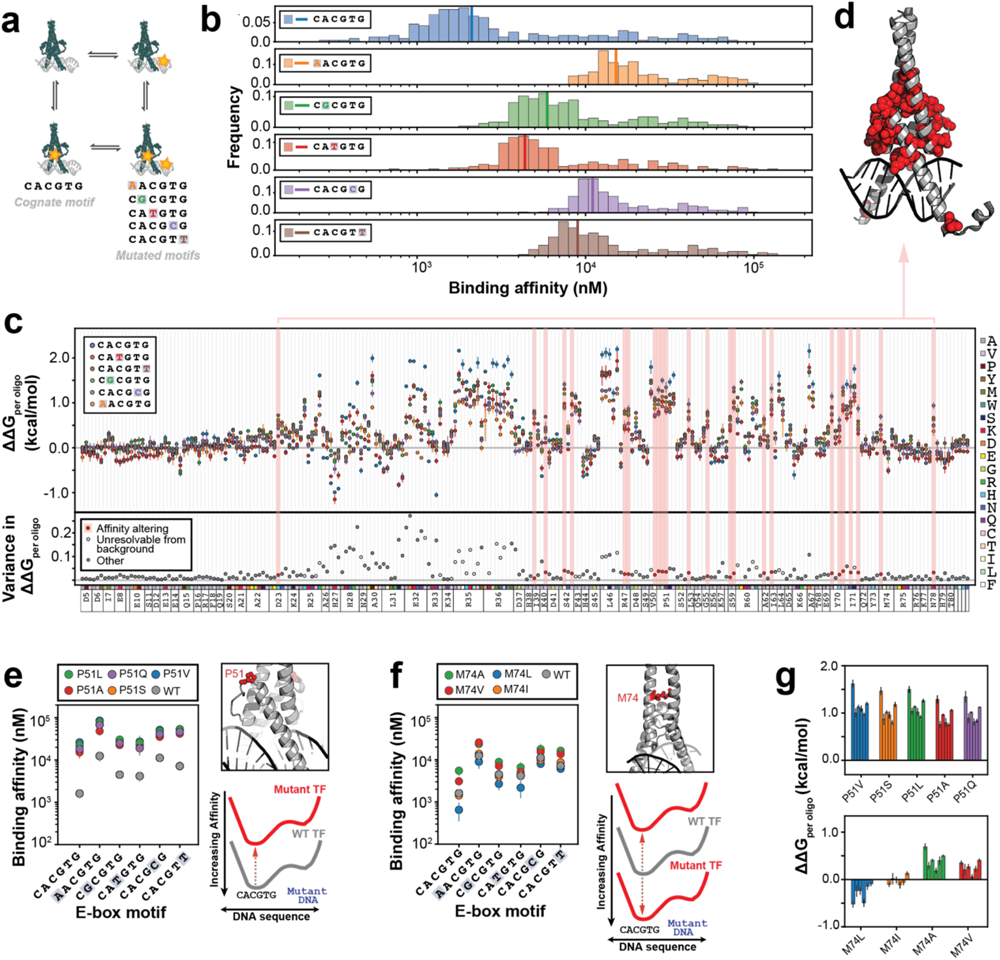
Dimer and loop mutations modulate MAX binding affinity. **(A)** Cartoon illustrating double mutant cycles across the TF/DNA interface. **(B)** Histograms of *K*_d_s for all MAX mutants against each measured E-box; vertical lines denote WT. **(C)** Measured ΔΔG_per oligo_ relative to WT across all E-boxes (median ± SEM) (*top*) and variance in ΔΔG_per oligo_ (*bottom*) for each mutation. Light grey markers indicate mutations where at least one motif was unresolvable from background; red markers indicate “affinity-altering” mutations. **(D)** “Affinity-altering” mutations projected on MAX structure (1HLO). **(E-F)** *K*_d_s (median ± SEM) for all E-box sequences (*left*), location of residue of interest (red) within MAX structure (1HLO, *top right*), and cartoon illustrating impact of mutations on DNA-binding landscape (*bottom right*) for substitutions to P51 (**E**) and M74 (**F)**. **(G)** ΔΔG_per oligo_ for MAX P51 (*top*) and M74 (*bottom*) mutations for all E-box sequences.

To identify TF mutations that modulate DNA-binding affinity (*i*.*e*. alter binding to all DNA sequences equally by shifting the binding energy landscape by a consistent free energy difference), we computed the variance across all measured ΔΔGs per mutant and selected mutants in the lowest quartile (**Methods**). Examination of the 24 identified “affinity-altering” mutations (**Figs 3C, S29**) revealed that they were generally located within the loop region or dimerization interface **(Fig. 3D)** and all weakened binding relative to WT MAX.

At some positions, all tested mutations had similar impacts. Substitutions to P51, a key residue for proper positioning of helix 1^43^, uniformly decreased binding affinity by ≥ 0.75 kcal/mol across DNA sequences (**Figs. 3E, 3G**). In contrast, mutations at other positions both increased and decreased affinity. MAX M74L, a mutation on the interior of the leucine zipper that forms stabilizing homotypic interactions, enhanced binding across all DNA sequences (consistent with prior studies of leucine zipper dimerization^56^), but other substitutions uniformly weakened binding (*e*.*g*. M74V, M74A). (**Figs. 3F, 3G**).

### Double-mutant cycles reveal drivers of DNA-binding specificity at the MAX-DNA interface

To systematically identify specificity-altering substitutions (which differentially affect binding to some DNA sequences, warping the binding energy landscape), we performed biochemical double mutant cycles^57^ comparing energetic impacts on binding from TF mutations and mutations to cognate DNA (**Figs. 4A, S30-S33)**. When visualized as a scatter plot, additive mutations to non-interacting residues and nucleotides lie along an additive fit line, while mutations to interacting and/or epistatic residue and nucleotide pairs yield non-additive energetic impacts that fall off-diagonal (**Fig. 4A**).

**Figure 4:**
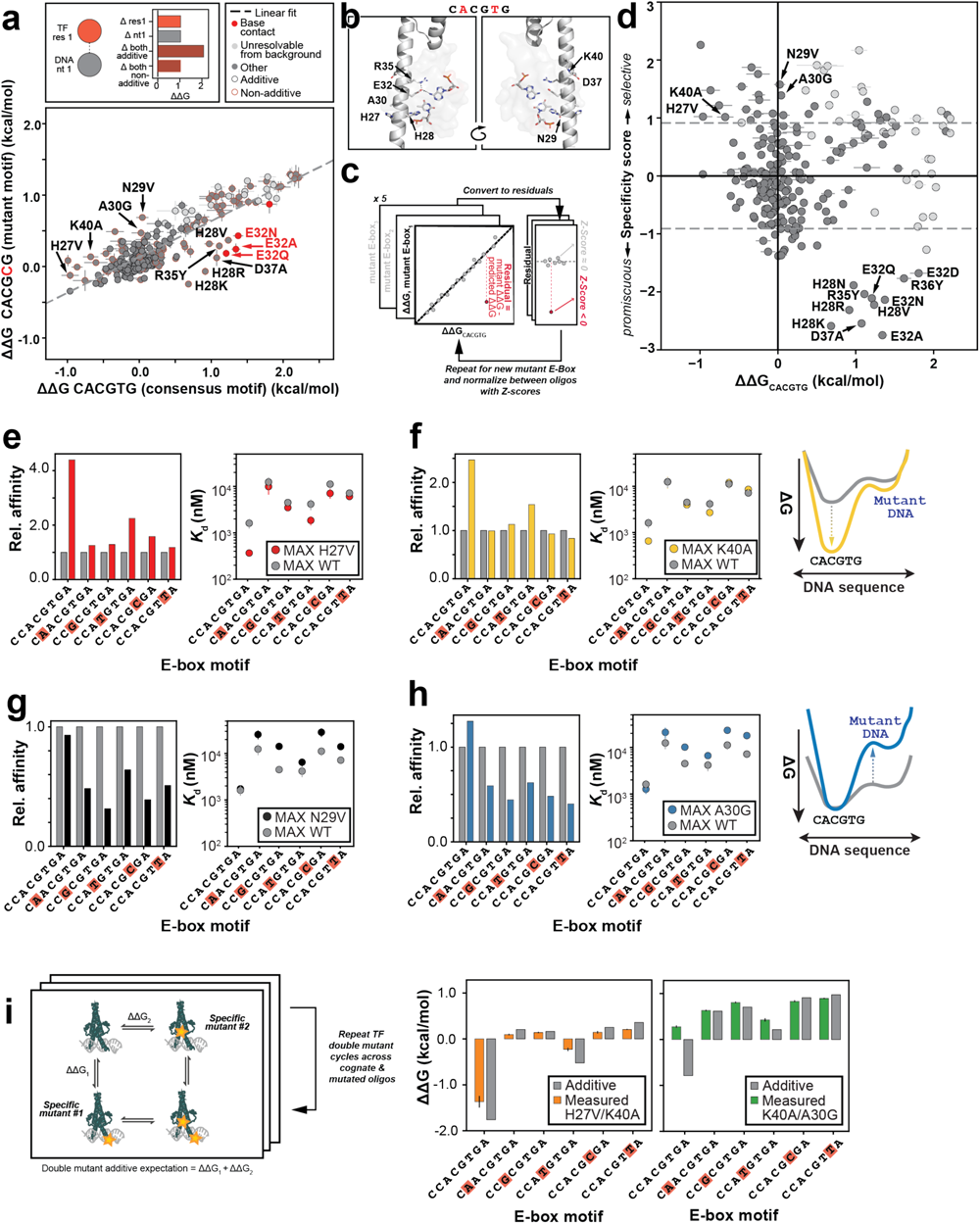
Mutations increase selectivity of MAX by enhancing cognate or weakening mutant DNA binding. **(A)** Cartoon depicting additive and epistatic energetic impacts for double mutants (*top*); pairwise comparisons between measured ΔΔG relative to WT for all MAX mutants interacting with low-affinity CACG**C**G versus cognate CACGTG. Light grey markers indicate mutations unresolvable from background for ≥1 DNA sequence; red markers indicate known crystallographic contacts to mutated nucleotide bases. Red marker edges indicate non-additive binding; dashed black line indicates linear regression. **(B)** Residues with epistatic energetic impacts shown on the MAX structure (1HLO). **(C)** Schematic illustrating calculation of “specificity scores” from double mutant cycles. **(D)** “Specificity scores” *vs*. ΔΔG_CACGTG_ for all MAX mutations. Light grey markers indicate mutations unresolvable from background for ≥1 DNA sequence; dashed grey line indicates thresholds for “specificity-altering” mutations. (**E-H**) Relative affinities (*left*) and median *K*_d_ ± SEM (*right*) across all E-box variants for WT and selected MAX mutants. **(I)** Cartoon depicting double mutant cycle analysis to probe for energetic coupling between selective TF mutations *(left*); results for 2 combinations of selective mutations (*right*).

While most (150) mutations additively alter binding to a low-affinity motif known to be bound by Myc/MAX heterodimers *in vivo* (CACG**C**G)^58^ relative to the cognate, 34 of the 81 non-additive mutants bind the mutated DNA with a reduced energetic penalty **(Fig. 4A, Table S6)**. These include mutations to structurally-predicted contacts, such as to E32 – which contacts the outer two nucleotide bases in the palindromic E-box motif (**CA**CG**TG**) – and residues that make stabilizing salt bridges with E32 like R35^59,60^. Some epistatic mutations, such as E32A, even alter the intrinsic sequence specificity of MAX to yield significant (*p*<0.05 via T-test) and absolute tighter binding to CACG**C**G; 10 additional mutations also alter sequence specificity to bind most tightly to other non-CACGTG E-box motifs (**Fig S34**). Other mutations non-additively bound CACGCG without an obvious structural rationale, such as solvent-facing D37 or H28 which canonically contact the 5’ guanine^44^ (**Fig. 4A-B**), suggesting that dynamic binding modes not apparent from low-energy bound structures contribute to sequence-specific binding (as explored further below).

### Mutations make MAX more selective by favoring the cognate or disfavoring non-cognate sequences

To identify mutations that increase selectivity for the cognate sequence relative to *many* mutated motifs, we computed residuals for each pairwise comparison between a mutated E-box and cognate sequence, and defined a ‘specificity score’ as the median of all residual Z-scores across each double mutant cycle comparison (**Figs. 4C, S35**). Mutants with negative specificity scores thus *decrease* the energetic penalty for binding to *mutant* motifs relative to the cognate (decreasing selectivity); as expected, many mutations to nucleotide base-contacting residues are among the most promiscuous mutations (*e*.*g*. H28, E32) (**Fig. 4D**). In contrast, mutants with positive specificity scores render MAX more selective. Strikingly, 22 MAX mutations increase selectivity for CACGTG (**Figs. 4D, S36, Methods**). Nearly all selectivity-increasing mutations lie in the DNA-contacting basic region or helix 1 (**Fig S37)** and are enriched for mutations at backbone-contacting residues (*p*=1*10^−3^ via Chi-squared test) and solvent-facing basic region residues (*p*=8*10^−7^ via Chi-squared test).

To understand how mutations change absolute affinities to increase selectivity, we examined specificity scores versus ΔΔG_CACGTG_ (**Fig. 4D**). Mutations in the upper-left quadrant – such as H27V and K40A – increase selectivity by disproportionately *increasing* affinity for the cognate sequence (*i*.*e*. “deepening the energetic well” for cognate binding) without altering affinity for many mutated sequences (**Fig. 4E-F**). Structural and mutational data suggests that these mutations often increase selectivity by stabilizing selective microstates (often by altering positioning of residues at the DNA interface). For example, given that all measured mutations at H27 position increase affinity for the cognate sequence **(Fig S38)** and to a lesser degree CA**T**GTG (the only mutated motif for which H28 mutations are additive) **(Fig S36, Table S6**), this suggests any substitutions at residue 27 may better position or reduce competition for protonation of H28 to promote selective E-box recognition.

Similarly, K40A does not directly contact DNA or DNA-contacting residues, but mutations at this position may disrupt a structurally predicted polar contact with D37^44^. As mutations to D37 reduce the energetic penalty for binding DNA sequences with mutations to the outer 2 bases in the E-box, disrupting this interaction may epistatically stabilize binding to all DNA sequences without mutations to these bases **(Fig 4F, S36)**. The aligned position in Pho4 contains an alanine (**Fig. 1B**), possibly contributing to Pho4’s selectivity via the same mechanism.

We note that no Pho4 mutations simultaneously increase affinity *and* selectivity for CACGTG (**Fig S39**). Instead, “affinity-altering” Pho4 substitutions simultaneously also increase affinity for mutated E-box sequences and are typically found within the basic region (rather than the dimerization region, as for MAX) (**Figs S40**). This Pho4 behavior is again consistent with a simple model of folding-and-binding where the same bound conformation recognizes all E-box variants, and mutations therefore proportionally change binding to all sequences.

MAX mutations in the upper middle of **Fig. 4D** increase selectivity by decreasing binding to non-CACGTG sequences (*i*.*e*. “raising the energetic wall” for non-cognate binding). For example, N29V (which ablates a structurally predicted polar contact with the DNA phosphate backbone^61^) does not alter affinity for CACGTG but *decreases* affinity for all measured mutated E-box sequences (**Fig 4G**). Similarly, mutating residue A30 (which faces the solvent on DNA-bound structures and is not adjacent to any other nucleotide base-contacting residues) to glycine leaves CACGTG binding unaltered but decreases affinities for all mutated E-boxes (**Fig 4H**). Mutations that “distinguish” between cognate and mutated motifs without any apparent sequence preference among mutated sequences suggest that the two alternative conformations hypothesized for MAX are either selective or non-specific, as explored below.

### Non-additivity of pairs of specificity-enhancing MAX mutations suggests existence of multiple binding conformations

To test for conformational partitioning between selective and nonspecific conformations, we examined mutant cycles between two protein residue pairs^62^ binding to many motifs (**Fig 4I, S41**). To exclude convolved effects of nearby mutations, we restricted interpretation to TF mutant pairs >15 Å apart in published structures (**Fig S42**). In this analysis, motif-independent binding additivity implies independent perturbations (such as rearrangement of local contacts that impact population of local microstates), while motif-dependent binding additivity implies distinct structural conformations such that the impact of a single mutation can be masked by altered macrostate occupancies.

Consistent with these expectations, the MAX H27V/K40A double mutant yielded additive energetic impacts relative to the single TF mutants for all measured E-box sequences (**Fig 4I**), with both mutations likely increasing selectivity by rearranging local contacts to independently enhance cognate binding. By contrast, the K40A/A30G double mutant yielded additive impacts for all motifs except for CACGTG, where impacts were less-than-additive (like A30G alone) (**Fig 4I)**. This non-additivity suggests that some mutations, like A30G, enhance motif selectivity by changing partitioning between multiple conformations with distinct sequence preferences: one that selectively binds CACGTG and another that promiscuously binds many mutated E-box sequences.

### Kinetic measurements provide insights into binding mechanism

Next, we asked if selective mutations caused changes in the MAX bound conformational ensemble, unbound ensemble, or both. As equilibrium binding measurements cannot resolve at which stages of the folding-and-binding reaction selective mutations alter microscopic transitions^63^, we developed a kinetic version of these microfluidic binding assays (k-STAMMP, derived from k-MITOMI^64^) (**Fig S43**). Specifically, we measured macroscopic dissociation rates (*k*_off,macroscopic_, hereon referred to as *k*_off_) (**Fig 5A**) and inferred apparent on-rates (inferred *k*_on,_= *k*_off_/*K*_d_, calculated assuming a macroscopic 2-state model in which TFs are either bound or not bound to DNA)^65^ for Pho4 and MAX variants interacting with 7 motif-variant DNA sequences^65^, totaling 610 total measured rate constants (**Fig S44-49**). Off-rates were well-fit by a single exponential for both Pho4 and MAX (**Fig 5A**), with measured rates typically varying by ∼2-fold between experiments.

**Figure 5:**
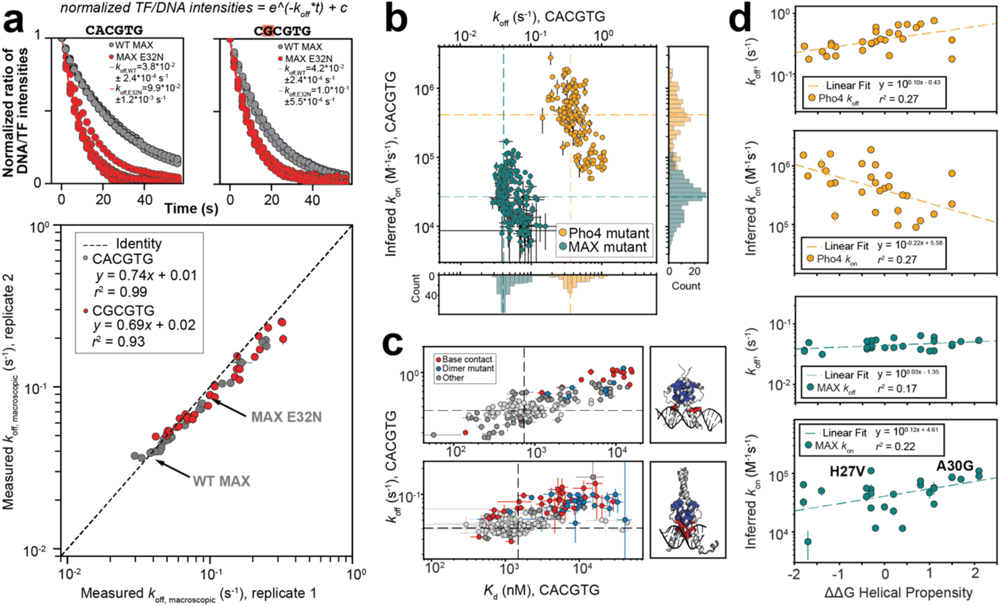
Binding kinetics suggest MAX and Pho4 differ in folding-and-binding transition. **(A)** Example dissociation traces for WT (grey) and E32N (red) variants of MAX (*top*); comparison of *k*_off, macroscopic_ measurements (median ± SEM) across two replicates of MAX and Pho4 variants dissociating from two sequences (*bottom*). Dashed black line indicates 1:1 relationship. **(B)** Inferred *k*_on_ versus measured *k*_off, macroscopic_ for all Pho4 (orange) and MAX (teal) mutants interacting with the cognate E-box (median ± SEM); dashed lines denote WT values. **(C)** *K*_d_ versus measured *k*_off, macroscopic_ for all Pho4 (*top*) and MAX (*bottom*) mutants interacting with the cognate E-box (median ± SEM); dashed lines denote WT values. Marker color indicates WT-like *k*_off_ values (*p* > 0.05, light grey) or mutations to known crystallographic DNA base contacts (red)/dimerization interface contacts (blue), imposed on Pho4 (1A0A) and MAX (1HLO) crystal structures. **(D)** Inferred *k*_on_ and measured *k*_off, macroscopic_ versus changes in helical propensity^53^ for non-DNA contacting basic region substitutions in MAX (teal) and Pho4 (orange); dashed line indicates linear fit.

Mutations to both Pho4 and MAX yielded larger changes to inferred on-rates than measured off-rates (7.8 and 5.6-fold change difference between fastest and slowest off-rates and 56.6 and 36.7-fold for on-rates for Pho4 and MAX, respectively) **(Fig. 5B**), consistent with recent work suggesting TF affinity and specificity is primarily governed by variation in association rates^20,66^. The subset of mutations that significantly changed dissociation rates tended to ablate or disrupt nucleotide (such as E32N), backbone, or dimer contacts **(Fig 5C)**, likely due to destabilization of the bound state(s).

### MAX and Pho4 differ in folding-and-binding transition, suggesting a conformation with reduced affinity for the cognate is more stable in MAX

For a folding-and-binding reaction with a single binding conformation, preformation of structure should increase *k*_on_ and decrease *k*_off_^67^. Consistent with this model, increasing helical propensity in Pho4 increased *k*_on,apparent,CACGTG_ and decreased *k*_off,CACGTG_ (**Fig. 5D**). In contrast, increasing helical propensity in MAX slightly decreased *k*_on, apparent_ and had little impact on *k*_off_ **(Fig. 5D)**. This observation is again consistent with the existence of multiple unbound binding-competent states in MAX such that the energetic impact of a mutation on cognate binding becomes uncoupled with intrinsic changes to helicity and instead alters conformational partitioning. Moreover, the observation that putatively helix-breaking mutations such as A30G speed up on-rate suggests that a weaker-binding conformation for CACGTG may be more stable. This is also consistent with stopped-flow kinetics data that suggest a conformational-change is the rate-limiting step for binding in MAX^68^.

### Selective mutations change binding landscape through different microscopic mechanisms

By the Hammond postulate, on-rates are more impacted by changes in the unbound state and off-rates by changes in the bound state. Investigating the inverse relationship between *k*_on_ and *k*_off_ across multiple DNA sequences (relative to WT) can therefore provide insight into which microscopic transition(s) are impacted^66^. MAX mutations to phosphate backbone-contacting residues that enhanced selectivity by weakening binding to non-cognate motifs (*e*.*g*. N29V, R60V; **Fig. 4G**) primarily altered *k*_off,apparent_ with little-to-no changes in inferred *k*_on,apparent_ (**Fig. 6A, S50**). Other mutations to non-DNA-contacting, solvent-facing residues that enhanced selectivity by selectively increasing affinity for the consensus motif (*e*.*g*. H27V, K40A) (**Fig. 4E-F**) primarily increased *k*_on,apparent_ for the cognate motif **(Fig. 6B, S50**) with little changes to off-rate across many measured motifs **(Fig. 6B**). This again implies that these mutations may change the unbound ensemble, increasing the rate of initial MAX DNA association in a DNA sequence-independent fashion. Finally, some selective mutations to solvent-facing residues (*e*.*g*. A30G) (**Fig. 4H**) altered both *k*_off,apparent_ and *k*_on,apparent_ (**Fig. 6C, S50**), suggesting changes to both bound and unbound states or to the transition state itself.

**Figure 6:**
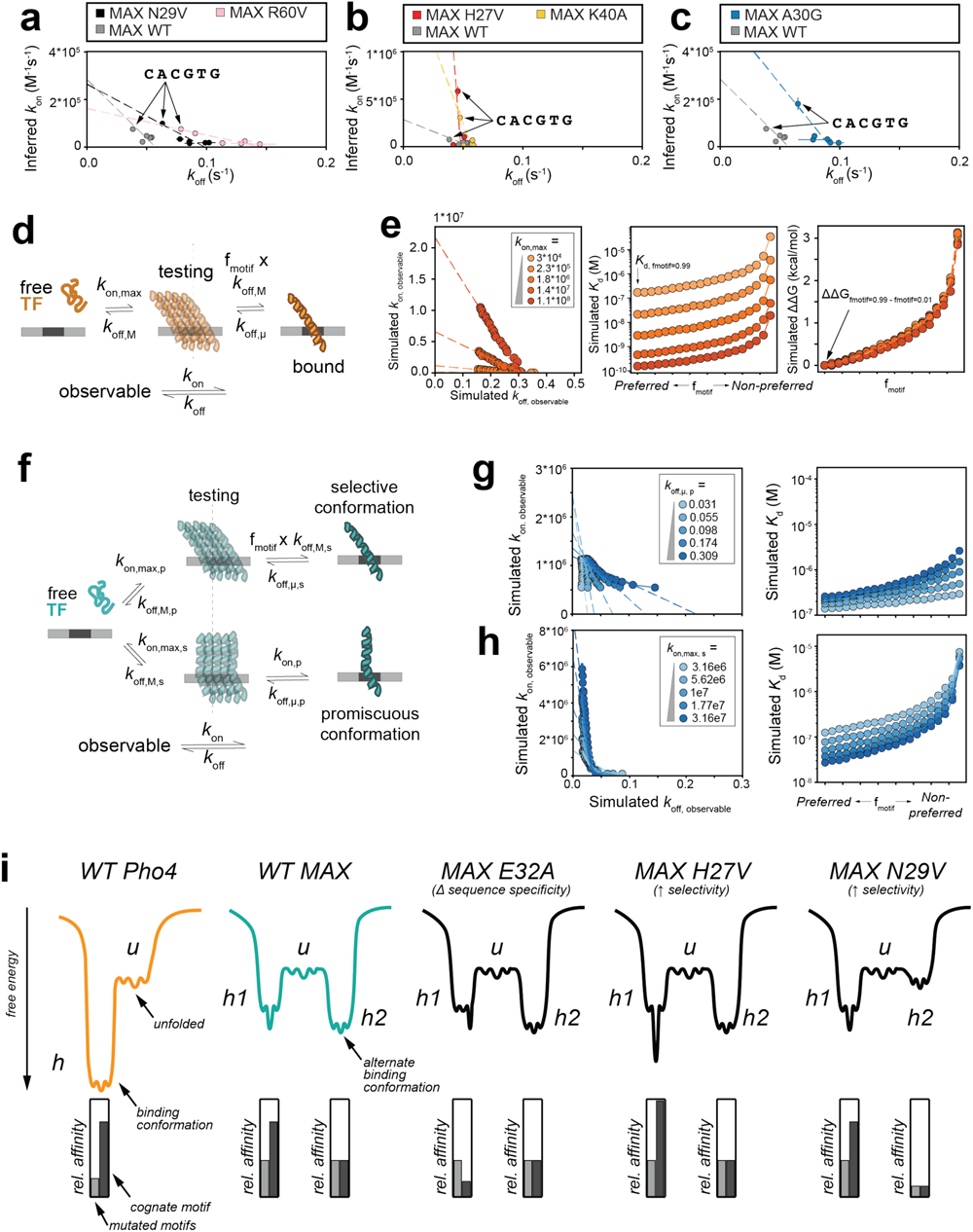
Selective mutations change the DNA binding landscape through different microscopic mechanisms. **(A-C)** Inferred *k*_on_ versus measured *k*_off, macroscopic_ for WT MAX and selective mutations across many E-box variants; dashed line indicates linear fit minimizing error in both x- and y-dimensions **(D)** Three-state model and associated microscopic rate constants for TF binding with a single bound conformation. **(E)** Simulated rate constants (*left*), affinities (*middle*), and differences in free energy of binding (relative to the most preferred sequence, *right*) as a function of microscopic on-rate and f_motif_ value with binding model illustrated in **(D)**. *k*on,max is the rate constant for transitioning between the *free* and *testing* states (representing a theoretical upper bound for the on-rate when all non-specific TF-DNA interactions result in specific binding), *k*off,μ is the rate constant for transitioning from the *bound* state to the *testing* state, and the probability of transitioning to the *bound* state depends on the likelihood of binding a given sequence (*f*_motif_) and the rate at which TFs transition from *testing* back to the *free* state (*f*_motif_ x *k*off,M). **(F)** Five-state model and associated microscopic rate constants for TF binding with multiple unbound and bound conformations with different intrinsic selectivities. Transitions to and from the selective and promiscuous ‘testing’ states are described by *k*_on,max,s_, *k*_off,M,s_, *k*_on,max,p_ and *k*_off,M,p_; transitions to and from the selective and promiscuous ‘bound’ states are described by the microscopic rate constants *f*_motif_ x *k*_off,M,s_, *k*_off,µ,s_, *k*_on,p_, and *k*_off,µ,s_. **(G-H)** Simulated rate constants (*left*) and affinities (*right*) as a function of microscopic on-rate and f_motif_ value with binding model illustrated in **(F)**. **(I)** Cartoon model illustrating idealized energetic landscapes for Pho4 (orange), MAX (teal), and three specificity-altering MAX mutations.

### Kinetic model with selective and promiscuous states reconstitutes measured changes

To test our proposed multi-state model of MAX binding and the microscopic origins of selective binding, we employed Gillespie simulations to model binding for a single TF and DNA molecule via multiple reaction schemes (**Fig. 6D-G**). For each reaction scheme, we sought to identify which, if any, changes in microscopic rate constants altered binding selectivity through similar kinetic and affinity pathways to those measured in selective MAX mutations.

First, we examined a 3-state model in which TFs are either unbound to DNA (‘free’), nonspecifically bound and ‘testing’ to see if a site underfoot represents the target site, or specifically bound (**Fig 6D**), identical to a scheme previously used to model *E. coli* LacI binding to various operator sequences^66^. Similar to observations for Pho4, systematically varying or co-varying rate constants globally shifted the binding landscape without changing selectivity (**Fig 6E, S51**); explicitly modeling folding-and-binding transitions also did not change selectivity (ΔΔG_f_motif=0.99 – fmotif=0.01_) (**Fig S52-53)**. These results are consistent with our experimental observations that Pho4 model mutations that alter helical propensity globally tune affinity (**Fig S39-40**) and hypothesis that explaining the mechanisms by which solvent-facing MAX mutations alter selectivity requires the existence of an additional state.

Consistent with MAX observations, changes to microscopic rate constants within a 5-state model in which proteins transition to 2 different helical^35^ ‘testing’ states that bind with different intrinsic selectivities (**Fig 6F**) yielded a variety of distinct affinity and selectivity effects (**Figs. 6G, S54**-**58**). Increasing the microscopic off-rate from the promiscuous state simultaneously increases macroscopic *k*_off,apparent_ and decreases macroscopic *k*_on,apparent_, mimicking the changes to binding kinetics observed for mutations that disrupt phosphate backbone contacts yet increase selectivity by reducing affinities for mutated E-box motifs (*e*.*g*. N29V and R60V, **Figs. 4E, 6G**). Similarly, increasing the microscopic on-rate to the specific state mimics effects seen for the solvent-facing H27V and K40A mutations that enhance selectivity by increasing cognate binding, increasing *k*_on,apparent_ while leaving *k*_off,apparent_ relatively unchanged (**Figs. 4F, 6H**). Thus, these mutations may increase energetic specificity by changing the unbound ensemble to “preconfigure” certain conformations with side-chains positioned for specific recognition. Combinations of changes to microscopic rate constants can even recapitulate more complex behaviors, such as those observed for A30G (**Fig S59)**. While we find that models with two differentially selective states (not models with a single binding conformation) are consistent with selective MAX mutant data, this toy model likely approximates two “macrostates” are each the sum of some large number of microstates in which residues at the DNA interface are differentially positioned within the folding landscape (**Fig 6I**). We conclude that consideration of multiple states with different intrinsic selectivity for the same set of sequences is necessary to explain kinetic and thermodynamic data for MAX.

## Discussion

Understanding selectivity – the quantitative difference in binding energy between preferred and non-preferred ligands – remains an unsolved biophysical question and unrealized engineering goal, with applications to binding generally beyond TF-nucleic acid interactions. Here, we investigated how mutations to MAX and Pho4, two structurally similar bHLH TFs with conserved DNA contacts yet different selectivity, alter binding to motif-variant DNA sequences. These measurements, in concert with kinetic and thermodynamic modeling, revealed putative non-contacting mutations in MAX that increased selectivity for the cognate motif via diverse molecular mechanisms: while some mutations likely stabilize selective microstates prior to binding (similar to mechanisms thought to drive antibody affinity maturation^69^), others change partitioning between different differentially selective macroscopic conformations (**Fig. 6I**). Pho4, in contrast, lacks evidence of appreciable alternate binding states, suggesting highly selective binding may be achieved with narrow folding funnels (lacking ability to access alternate conformations or rearrangements) (**Fig. 6I**). Overall, our results demonstrate that high-throughput measurements of mutational impacts on binding affinities and kinetics can reveal important properties about conformational ensembles difficult to resolve via other methods, and that these properties can dramatically impact the selectivity of otherwise highly similar proteins.

The observed selectivity differences between Pho4 and MAX may represent distinct evolutionary pressures stemming from their different biological roles and speed/specificity tradeoffs within different genome sizes^70^. Pho4 initiates transcription in response to phosphate stress^71^, while MAX acts as a heterodimerization node to control cell proliferation in concert with other TFs^72^. The observed decreased “mutational sensitivity” of MAX compared to Pho4 (**Fig. 2C**) may result from a need to preserve a wide variety of existing functions and reflect the fact that mutations in promiscuous binders may be more likely to yield functional binding proteins^73–76^. Finally, our observed non-additivity of selective mutations (**Fig 4I & S41**) suggest a rugged mutational landscape that complicates protein engineering efforts to combine favorable mutations to enhance selectivity^77,78^.

This work is aligned with many other investigations linking conformational ensembles to TF specificity, from bispecific binding to divergent motifs^5,38,39,79,80^ to structural characterization of selective and promiscuous complexes^81–84^. These selective and promiscuous conformations are not just static bound states; TFs undergo conformational rearrangements between these complexes with varying degrees of selectivity as part of the binding pathway^40,85,86^. Moreover, the ability to access different conformations – and therefore bind increasingly diverse sites – can originate from decreased global fold stability^76^.

This work highlights the need for new data detailing how selective mutations discriminate between not just a handful of motif-like sequences, but rather large landscapes of diverse sites. Obtaining these measurements will be essential for improved design of selective binders. While algorithms to design synthetic TF-like binders with user-specified sequence specificity^9,10^ are increasingly successful, attempts to improve selectivity – such as by mutating contacts involved in “non-specific” contacts like charged interactions with the phosphate backbone – yield scaffold-dependent success^9^ (**Fig 4G, 6A**). Our work suggests that prediction and design of selective binders (beyond TF-DNA interactions) will necessitate consideration of energy landscapes that govern both folding and specific recognition. Currently, many structure-based binding algorithms cannot capture this information; we predict that incorporating conformational dynamics will be essential for properly predicting and engineering molecular specificity.

## Materials and Methods

### Data acquisition and curve fitting

#### Fabrication of microfluidic molds and devices

Flow and control molds were fabricated as described previously^1,2^ and all design files are available on the Fordyce Lab website (http://www.fordycelab.com/microfluidic-design-files).

We fabricated two-layer MITOMI devices from these molds using polydimethylsiloxane (PDMS) polymer (RS Hughes, RTV615) in the Stanford Microfluidics Foundry. To fabricate the control layer, we combined 55 g of PDMS (1:5 ratio of cross-linker to base), mixed and degassed the components within a centrifugal mixer at 2000 and 2200 rpm, respectively, for 3 minutes each (THINKY). We then poured the mixture onto the molds, degassed them in a vacuum chamber for 45 minutes under house vacuum, and baked them in an 80°C convection oven for 40 minutes. We then cut control layers for individual devices from the cast PDMS and punched fluid inlet lines using a drill press (Technical Innovations) with a mounted catheter hole punch (SYNEO, CR0350255N20R4).

To fabricate the flow layer, we combined PDMS at a 1:20 ratio (cross-linker to base) and mixed and degassed the components within a centrifugal mixer at 2000 and 2200 rpm, respectively, for 3 minutes each. We then spin-cast the PDMS onto molds for 10 s at 500 rpm followed by 1850 rpm for 75 s. Spin-cast layers were allowed to relax on a flat surface at room temperature for 5 minutes before baking at 80°C for 40 minutes. We then manually aligned individual control layers to the partially cured flow layer and baked the aligned devices for an additional 40 minutes at 80°C. Bonded two-layer devices were cut from the flow mold with a scalpel and the flow-layer fluid inlet lines were punched as described above.

#### QuikChange Mutagenesis for MAX mutant library

##### MAX Plasmid

We generated a MAX plasmid carrying the full sequence of the MAX transcription factor with a c-terminal monomeric eGFP tag^3^ separated from the MAX coding sequence via a gly-ser linker (GGSGGGGS). We used Gibson assembly to clone the MAX-eGFP fusion into a purified, linearized PURExpress vector with ampicillin resistance. The construct was sequenced validated using Sanger sequencing prior to generating mutants.

##### Mutagenesis primer design

Primers encoding mutants were generated as described previously^4,5^ using a custom-made program, available at (https://github.com/FordyceLab/designQuikChangePrimers). Briefly, the program takes as input the DNA sequence encoding the MAX ORF sequence and a list of desired mutants (e.g. “A67D” for Ala 67 to Asp mutation), generates a set of candidate primers for each mutant, and returns suggested mutagenic primer pairs scored according to criteria previously published in the QuikChange manual. Primers were ordered in a 96-well plate format from IDT (Integrated DNA Technologies) at the 10nmol synthesis scale with standard desalting purification; the forward and reverse primers for each mutant were premixed in each well. For library design, pathogenic mutations and VUS were curated from clinical sequencing databases as of March 2021.

##### Plate-based QuikChange mutagenesis

Mutagenesis reactions were performed as previously reported^4^ in a 96-well plate format. Each well contained its own mutagenesis reaction. Reactions were performed using the QuikChange protocol as directed by the manufacturer (Agilent Technologies, New England Biolabs). Upon completion of mutagenesis, we digested any remaining methylated wildtype plasmid using Dpn1 (New England Biolabs, R0176L) for 1 hour at 37°C. We then transformed 1µL of each reaction into 5µL of competent *E. coli* DH5alpha cells (New England Biolabs, C2987I). Transformants were grown to saturation in 5-8mL of LB media supplemented with ampicillin (100µg/mL) and miniprepped (Qiagen) for Sanger sequencing. To validate successful mutagenesis, we aligned each sequence to the template ORF and ensured that only the intended mutation was present in the plasmid. We re-picked colonies in the event of errant mutations elsewhere in construct (eg. indels, additional mutations in plasmid), or poor sequencing quality.

#### Plasmid Array Printing

##### Plate preparation

Prior to printing plasmids, we transferred mini-prepped plasmid into 96-well plates. To standardize volumes of plasmids, the wells were evaporated to dryness. We resuspended each plasmid with 50uL of Milli-Q water. Plasmids were transferred from 96-well plates into 384-well plates using a Biomek FX Automated Workstation (Beckman Coulter, model A31843). Each plasmid was pipetted into 4 consecutive wells within the 384-well plate, and each well of the 384-well plate contained 10µL of plasmid. We recorded positions to keep track of empty wells for adding subsequent mutants manually.

We evaporated 384-well plates to dryness at room temperature and resuspended dried wells in print buffer formulated as below:

- 1% (10mg/mL) Bovine Serum Albumin (Sigma Life Science, B4287-25G)
- 200mM (11.65 mg/mL) NaCl (Sigma Life Science, 71376-1KG)
- 12mg/mL trehalose dihydrate (Sigma Life Science, T9531-25G)

All reagents were combined in Milli-Q water and mixed to dissolution at room temperature and sterile filtered to remove aggregates. To each well in the 384-well plate, we added 12-15µL of print buffer for arrayer printing. When not in use, we sealed plasmid plates with foil covers and stored them at -20°C. Prior to printing, plates were defrosted overnight at 4°C and centrifuged at 2000 RPM for 5-10 minutes. Over the course of subsequent prints, we added ∼3-5µL of Milli-Q water (or additional print buffer) as needed to ensure sufficient volumes of sample in plates for printing.

##### Plasmid printing & device alignment

We printed plasmids using a SciFlex Arrayer (SCIENION AG) using either the PDC50 or PDC70 nozzle (Type 1 coating). We generated a “field file” to map each well on a 384-well plate to positions within the printed plasmid array. To prevent cross-contamination between plasmids, the glass nozzle was washed with room temperature Milli-Q water in between spotting different plasmid samples. We printed plasmid arrays on epoxysilane-coated glass slides (ArrayIt SME2, SuperChip C50-5588-M20, or self-coated as previously described^6^). After drying arrays overnight at room temperature, we aligned microfluidic devices to “program” each chamber with its own plasmid spot. Prior to alignment, we pre-baked microfluidic devices at 80°C for 20-25 minutes using a hotplate (Torrey Pines Scientific) and allowed them to cool to room temperature. These devices were then baked for 4-4.5 hours at 95°C on a hotplate.

#### Preparation of DNA for fluorescence-based binding assays

We designed all DNA sequences for binding assays with a 3’ region complementary to a AlexaFluor-647 dye-conjugated primer (anneal temperature: 37°C) (See **Table S4**).

##### Double-stranded DNA preparation and dilution

We ordered all DNA sequences as single-stranded oligonucleotides from Integrated DNA Technologies (IDT) with standard desalting purification and shipped in ‘LabReady’ formulation (100µM in IDTE buffer, pH 8.0). We then duplexed these single-stranded DNA sequences by (1) annealing the universal AlexaFluor-647-labeled primer to the 3’ region of the oligonucleotide and (2) extension using the primer as a template using Klenow fragment, exo^-^, polymerase. Both steps (1) and (2) were performed as previously described^4^.

After the Klenow extension, we filtered the DNA reactions using a 0.45µm filter spin column. We subsequently equilibrated duplexed DNA in the final assay buffer (10mM Tris-HCl, 100mM NaCl, 1mM DTT, pH 7.5; aliquoted and filtered using 0.45 mM Steriflip vacuum (Millipore, SE1M179M6)) using 10K filter spin concentrator columns (Amicon Ultra, UFC501096). We added ∼100µL of the duplexed DNA to the filter spin columns, added 200µL of assay buffer, mixed by pipetting, and concentrated the reaction to 100µL by centrifugation (9000RPM for 8 minutes). We repeated this process 5 times, and subsequently eluted the equilibrated DNA via manufacturer’s instructions for the 10K filter spin concentrator column.

We serially diluted equilibrated DNA in final assay buffer as previously described^4^. For this dilution and the subsequent assay, the assay buffer was supplemented with 50µg/mL of UltraPure BSA (ThermoFisher, AM2618). To calibrate each step of the binding assay with a DNA concentration, we measured the highest concentration of DNA using a DeNovix to measure absorbance at 260nm.

For all experiments involving a mutation within the core-site, we also performed this procedure for the consensus DNA sequence 5’-C CACGTG A-3’. For these oligonucleotides, we measured binding isotherms for 5 DNA concentrations. For the sixth measurement, we introduced the duplexed and labeled reference DNA sequence at a high concentration (∼7-9µM) so that we could accurately quantify the saturation ratio with which to fit all binding isotherms.

#### Microscopy and instrumentation

We made measurements as previously described^4,5^ using a Nikon Ti-S microscope. Devices were controlled using a pneumatics manifold^7^. Custom scripting and automation enabled integrated control of both the microscope and the pneumatics manifold (https://github.com/FordyceLab/RunPack).

#### Measuring *K*_d_ values on-chip via STAMMP

Measuring *K*_d_ values on-chip have 3 major steps: (1) On-chip expression and purification of MAX mutants, (2) titration of fluorescently labeled DNA, (3) image analysis and calculation of *K*_d_ values.

##### On-chip expression & purification of MAX mutants

On-chip expression and purification require the immobilization of expressed proteins for subsequent assay steps. To accomplish this, we took devices aligned to printed plasmids and performed a series of passivation steps as described previously^4,5^ to immobilize biotinylated anti-GFP antibodies selectively underneath the ‘button valves’ of the STAMMP microfluidic device. As all TF variants are fused to a GFP, these antibodies will trap recruited TF variants for subsequent assays.

To express all TF variants simultaneously, we used PURExpress (NEB E6800L). Briefly, we equilibrated Parts A and B of PURExpress on ice until defrosted. For one device using 25µL total of PURExpress, we first incubated 10µL of Part A with 7.5 µL of Part B on ice for 45 minutes. Then, we added 1.5µL of recombinant RNAsin (Promega N2515) and 6µL of nuclease-free water (Promega P1193) and mixed by pipette until no phase separation was visible. We introduced PURExpress onto the device as previously described^4,5^ Devices were then placed on a pre-heated hotplate at 37°C for 45 minutes to express all proteins. We then placed devices on the scope and allowed the GFP to fold over the course of 45-60 minutes with the button valves on the device closed. After this was completed, we opened the button valves and recruited GFP-tagged protein to the antibodies for 20-30 minutes. We then closed the buttons to shield trapped TFs while we washed the device with PBS and TrypLE (ThermoFisher 12604-013) to remove nonspecifically bound TFs from the device walls. After this, we equilibrated the device with assay buffer to remove trace amounts of TrypLE and to equilibrate proteins in assay buffer, composed as follows unless otherwise specified:

- 20 mM Tris-HCl pH 7.5 (from 100 mM stock)
- 100 mM NaCl (from 100 mM stock)
- 1 mM DTT (from 1 M stock) (Sigma-Aldrich, D9779)
- 50 ug/mL ultrapure BSA (ThermoFisher, AM2618)

##### DNA Binding measurements

Binding measurements were performed as described previously^4^. Briefly, we introduced fluorescent DNA (prepared as described above) at 6 concentrations ranging between ∼60nM to ∼6µM on the device. For binding measurements with DNA sequences containing mutations within the core binding site, only five concentration points were measured. For the sixth and final concentration point, we measured DNA binding for the reference DNA sequence 5’-C CACGTG A-3’ at a high concentration to determine DNA to protein fluorescence intensity ratio denoting saturation of all binding sites for global fitting *K*_d_ values. For prewash Cy5 images, we imaged the device at multiple exposure times, ranging from 30 ms to 100 ms. We imaged postwash GFP images using an exposure time of 500 ms. For postwash Cy5 images, we used exposure times of either 1200 ms or 3000 ms to ensure we did not collect measurements at a saturating intensity.

##### Image analysis

Image analysis and calculation of *K*_d_ values were largely performed as previously described^4^. Briefly, images were stitched using in-house Python packages ImageStitcher (https://github.com/FordyceLab/ImageStitcher). These images were then analyzed using the ProcessingPack package (https://github.com/FordyceLab/ProcessingPack), largely as previously reported^4^.

Briefly, to quantify affinities for each TF mutant binding to a given DNA sequence, we acquired per-chamber calibration curves relating observed AlexaFluor-647 fluorescence to spectroscopically measured dsDNA concentrations (**Fig. S6**), converted intensities to DNA concentrations based on orthogonal measurements using a DeNovix instrument, and then fit concentration-dependent binding curves as described below.

To identify TF mutants with DNA binding statistically indistinguishable from background, we compared Cy5 intensities from TF-containing chambers with those from blank chambers by repeated measures ANOVA (providing a conservative estimate of mutants with detectable binding); we report measured *K*_d_s for these variants as a lower limit **(Fig. S7, Table S3)**.

##### Calculation of K_d_ values

To fit dissociation constants, we first measured the amount of DNA bound to surface-immobilized TF mutants over multiple concentrations and converted these to ratios of bound DNA intensities (Cy5 channel) over immobilized TF (eGFP channel). We then applied a global fit to the measured DNA/TF ratios and fit data from each individual chamber to single-site binding models^1,8^.

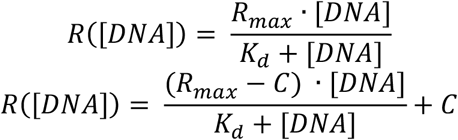

Here, R is the intensity of DNA/TF as a function of DNA concentration within the chamber, R_max_ is the constant shared across all chambers corresponding to the value at which all binding curves saturate (assuming an identical molecular stoichiometry), [DNA] corresponds to the concentration of free DNA within the chamber, and *K*_d_ is the dissociation constant for a particular chamber. We determined the R_max_ value by taking the median of the top 10% of DNA binding MAX mutants at the highest DNA concentration point in an experiment for the reference DNA sequence; for experiments with a mutated DNA sequence, we measured the highest DNA concentration point using a reference DNA sequence to prevent underestimation of R_max_.

In addition to fitting to a Langmuir isotherm, we fit our data to a modified single-site binding model with an offset value, C, to correct for variations in background intensities between experiments that can affect ratio values. The fitting method that minimized per-chamber RMSE of fits for each technical replicate was used for final determination and export of *K*_d_ values.

#### Measuring *k*_off_ values on-chip via k-STAMMP

At the end of a STAMMP binding assay, *k*_off_ values can also be optionally obtained. Measuring *k*_off_ values on-chip adds two additional steps to a STAMMP assay: (1) titration of unlabeled DNA and (2) image analysis and calculation of kinetic constants.

##### Dissociation measurements

Dissociation rate data was acquired after equilibrium binding procedures largely as previously described^9,10^. We first flushed each device with non-fluorescent (dark) competitor dsDNA oligonucleotides containing an E-box motif at a concentration of ∼0.9 μM diluted in PBS for 10 minutes with button valves closed after the acquisition of the “post-wash” image. These oligos were prepared with Klenow polymerase as described above but with unlabeled primers. The inclusion of non-fluorescent (dark) competitor at high concentrations during dissociation is critical to prevent rebinding of labeled material, which leads to systematic underestimation of dissociation rates^10^.

Next, after stopping flow of unlabeled competitor dsDNA and closing sandwich valves, we then opened the buttons for 2.0 seconds to allow dissociation of bound fluorescent DNA from surface-immobilized TF. Finally, we closed buttons, flushed the device, and imaged in both the Cy5 and eGFP channels to quantify loss of DNA binding and surface-bound TF, respectively. For each experiment, we iterated this process for 40 button duty cycle iterations.

##### Calculation of kinetic constants

After acquiring and processing images as described for STAMMP assays, kinetic constants (*k*_off_) were determined by first calculating the ratio (R) of “post-wash” DNA fluorescence (Alexa 647) to “post-wash” GFP fluorescence per chamber at each time point. This ratio was then used to fit a single exponential value:

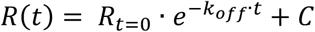

where R(t) is the fluorescence ratio as a function of time, k is the dissociation constant, and C is a constant term which accounts for background fluorescence or non-specific sticking of DNA. From these fitted *k*_off_ values, we can infer *k*_*<u>on</u>*_ through the definition of the dissociation constant from measured *K*_d_ and measured *k*_off_ as previously described^10,11^:

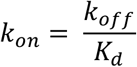

### Data interpretation

#### Calculation of fold-reduction in binding from MITOMI measurements

To calculate fold-reduction in binding for MAX and Pho4 from previous measurements^8^, we collected the measured affinities for all sequences with a Hamming distance of one away from the consensus motif (**Fig. 1E**), compared these binding affinities to the median of all consensus motif measurements (with variable flank nucleotides), and calculated and reported the 90^th^ percentile for fold-reduction in binding.

Thermodynamic modeling of coupled folding-and-binding equilibria

Thermodynamic model fitting and binding simulations were defined by the following variables:

1. *H*, the percentage of TF that is folded (helical) in solution.
2. *C*, the percentage of TF that is unfolded (coil) in solution.
3. *D*, the concentration of free DNA.
4. *Co* (for Complex), the bound TF-DNA complex.
5. *pT* (for total protein), the total amount of TF available in the reaction.
6. *dt* (for total DNA), the total amount of DNA available in the reaction.

These variables were used to construct the following equations and define equilibrium constants:

1. Mass balance equation for protein species, defined as:

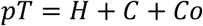
2. Mass balance equation for DNA, defined as:

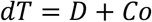
3. Equilibrium constant defining partitioning between folded and unfolded states in the unbound state:

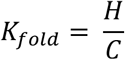
4. True binding equilibrium constant, where only the folded (helical) form can complex with DNA:

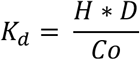

##### Developing a function for K_d,apparent_

In a STAMMP experiment, only the total amount of free DNA, total amount of immobilized TF, and fractional occupancy of bound TF-DNA complex is known; the distribution of folded/unfolded unbound states and true values underlying equilibrium constants is not known. Therefore, first we used the preceding 4 equations and sympy.solve to define: *Co* ( *pT, dT*; *K*_*fold*_, *K*_*d*_), the concentration of bound TF-DNA complexes as a function of total protein and DNA given the folding and binding equilibrium constants, as follows:

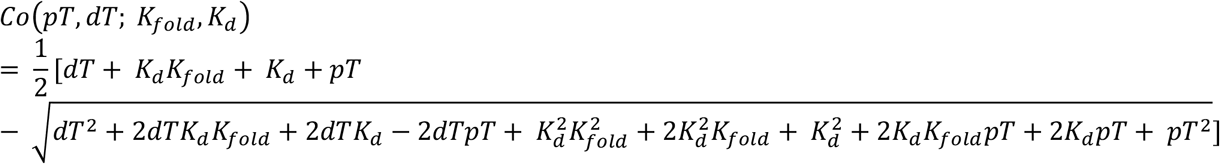

For all subsequent calculations, pT was defined as 50 nM, based on previous estimates for the concentration of immobilized protein on MITOMI microfluidic devices^5^. Apparent (measured) DNA-binding affinities were then obtained by: (1) calculating equilibrium occupancy of *Co* at *dT* spanning from 0 to 10 *μM*, analogous to the procedure for measuring binding affinities in STAMMP experiments, and then (2) defining the *dT* resulting in 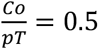 as the apparent DNA-binding *K*_*d,apparent*._

##### Defining changes in K_fold_ for helicity-altering TF mutations

TF mutations that alter the propensity to fold or unfold in the unbound state can also alter the observed DNA binding affinity. We assumed that all surveyed TF mutations only change the free energy difference between the folded and unfolded state, changing *K*_*fold*_ but not *K*_*d*_. The amount by which a TF mutation changes *K*_*fold*_ can then be defined as the change in helical propensity relative to WT TF, which changes the folding equilibrium as follows:

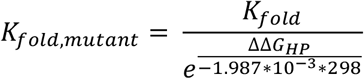

where ΔΔ*G*_*HP*_ is defined as the change in helical propensity, which defines the free energy difference for partitioning between unfolded and folded, helical states.

##### Fitting STAMMP data to derived thermodynamic model

To first develop intuition for how *K*_*d,true*,_ *K*_*fold*_, and ??G_HP_ alter the range of *K*_*d,measured*_ for both WT and mutant TFs, we determined how the expected linear free energy relationships between helical propensity-altering TF mutants and apparent DNA binding affinity impact *K*_*d,measured*_ for TF mutations with intrinsic changes in helical propensity spanning -2.0 to 2.5 kcal/mol (**Figures S13 and S14**).

Next, we fit *K*_*d,measured*_ for Max and Pho4 to the thermodynamic model defined for *Co*(*K*_*d*_, *K*_*fold*_, *dT, pT*) to extract *K*_*d,true*_ and *K*_*fold*_ for WT TF (**Figure S15**). To accomplish this, we first restricted our analysis to TF mutations in the basic region that do not make crystal contacts with DNA or at the dimerization interfaces, as these mutants presumably alter *K*_*d,measured*_ through mechanisms other than changes to *K*_*fold*_. Each of these TF mutations was then defined to have a ??G_HP_ in accordance with previously measured changes in Gibbs free energy for helix formation^12^. Next, (for both MAX and Pho4) we calculated the RMSE of log_10_ thermodynamic model-predicted *K*_*d,measured*_ for all measured TF mutations to the STAMMP-derived log_10_ *K*_*d,measured*_ for values of *K*_*d,true,WT*_ ranging from 1 to 103.5 nM and values of *K*_*fold,WT*_ where the fraction of unfolded TF in the unbound state ranged from 99 to 1 percent. The fitted values of *K*_*d,true,WT*_ and *K*_*fold,WT*_ were defined as those that minimized RMSE for the mutations and *K*_*d,measured*_s measured via STAMMP relative to the thermodynamic model of folding-and-binding. Code to reproduce all simulations and fitting procedures is available at https://osf.io/jmz8t.

#### Identification of “affinity-altering” mutations

To identify “affinity-altering” mutations, we: (1) calculated the free energy change (ΔΔG) imparted by each TF mutation on binding to each DNA sequence relative to WT MAX and computed the variance across this set of 6 ΔΔG values, (2) eliminated sequences with strongly differential effects on ΔΔGs by excluding MAX mutations in the top quartile of variance, and (3) excluded mutations unresolvable from background binding to any DNA sequence (**Fig. S7, Table S4**) and mutations that do not significantly alter affinities relative to WT MAX (*p* < 0.05 via independent T-test) in all measured E-Box sequences (**Figs. 3C, S29**).

#### Identification of non-additive TF+DNA mutation pairs

Identifying epistasis across the TF-DNA interface requires 4 affinity measurements: (1) WT TF binding a ‘reference’ DNA sequence, (2) mutant TF binding a ‘reference’ DNA sequence, (3) WT TF binding a ‘mutant’ DNA sequence, and (4) mutant TF binding a ‘mutant’ DNA sequence. We then determined if each pair was statistically significantly non-additive in *K*_d_ space, largely as previously reported^4^.

Briefly, to visualize the concentration-dependent binding behavior that would have been expected if the energetic effects of TF and oligonucleotide mutations were purely additive, we first calculated an expected ‘additive’ *K*_d_ value using the median reference *K*_d_ value (for WT TF interacting with the ‘reference’ oligonucleotide), the median *K*_d_ resulting from the relevant oligonucleotide mutation alone, and the median *K*_d_ resulting from the TF mutation alone as follows:

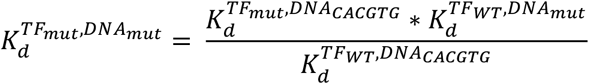

To determine whether the candidate TF mutant appeared epistatic with the DNA nucleotide mutation, we used measurements of (1) WT MAX and CACGTG, (2) WT MAX and mutant DNA, and (3) mutant MAX and WT DNA to generate a distribution of additive *K*_d_ measurements (n=500 simulated additive measurements). We then performed an independent T-test comparing the distribution of ‘additive’ affinities with the experimentally measured affinities for the double-mutant and used a p-value cut-off of 0.05 to define TF mutants that are epistatic with DNA mutants.

#### Identification of “selective” MAX mutations

To identify mutations that differentially increase selectivity, we computed residuals for each pairwise comparison between a mutated E-box motif and the CACGTG cognate, calculated Z-scores for each residual (to account for the fact that residual distributions vary with absolute affinity), and defined a ‘specificity score’ as the median of all Z-scores across each double mutant cycle comparison (**Figs. 4C, S35**), with ‘selective’ mutations exceeding a threshold defined by the standard deviation of the Gaussian fit to the residuals (**Fig S37)**. Mutations which were unresolvable from background in the cognate motif measurement (for which reported *K*_d_s are underestimated) were excluded from the list of reported ‘selective’ mutations, and candidate ‘selective’ mutations were inspected and culled by eye.

#### Gillespie model of TF binding kinetics with one or more binding conformations

Gillespie algorithms are stochastic simulations based on reaction rates that use discrete molecule counts and variable time steps^13^. Here, we simulated TF “energetic specificity” using Gillespie models of different binding pathways depicted in **Fig. 6D and 6F**. At each time step, we compute: (1) How long until the next reaction occurs? and (2) Which reaction happens?

First, we calculated reaction propensities (a) from reaction probabilities (c) and the number of reactants available for each reaction. Reaction probabilities can be derived from the kinetic rate constants as previously described^10^. For all simulations, microscopic rate parameters previously estimated from CTMC modeling were used as a starting estimation^10^. In this model, we initialized with 1 molecule of MAX and DNA and set the volume to 1.66*10^−12^ pL (chosen for simplicity so simulated s^-1^ values equal M s^-1^ on-rate constants).

Observed off-rates (macroscopic *k*_off_) were calculated as the median value across 3 replicates of the inverse of the average time it takes MAX to become fully dissociated once specifically bound; observed on-rates (macroscopic *k*_on_) were calculated as the median value across 3 replicates of the inverse of the average time it takes MAX to become specifically bound once dissociated in solution. The observed *K*_d_ is calculated as the ratio between macroscopic on- and off-rates.

To calculate “energetic specificity”, we calculated *k*_off_, *k*_on_, and *K*_d_ for a range of “motifs”, ranging from “strongly” to “weakly” bound sequences. “Motif strength” is defined in all models by *f*_*motif*_, an implicit parameterization of the probability of binding such that an increase in *f*_*motif*_ (tightly bound motifs) causes an increase in association rate and a decrease in off-rate, as previously described^10,14^. Given that all observed specificity-increasing mutations do not occur at conserved nucleotide contacting residues, we assumed that TF mutations do not change the intrinsic probability of recognizing a motif (*f*_*motif*_) but instead only alter microscopic rate parameters. Specificity was defined as the free energy difference between the “strongest” (*f*_*motif*_ = 0.99) and “weakest” (*f*_*motif*_ = 0.01) motif surveyed. Code to reproduce all simulations is available at https://osf.io/jmz8t.

##### Sensitivity analysis for 3-state model

Reaction likelihoods were defined according to the 3-state model consisting of unbound, testing, and bound states shown in **Fig. 6D**. Simulated *k*_on_, *k*_off,_ and *K*_d_ values across 20 “motifs” of strengths ranging from *f*_*motif*_ = 0.99 to 0.01 were obtained by coarsely varying 3 free microscopic rate constants *k*_*on, max*_, *k*_*off, max*_, and *k*_*off, u*_ across 4 orders of magnitude each in 10 step increments. For each combination of free parameters, we simulated 3 independent trajectories with 10^4^ reaction steps. The resulting free energy difference between the tightest and weakest surveyed motifs was calculated.

##### Sensitivity analysis for 4-state model

Reaction likelihoods were defined according to the 4-state model consisting of an unbound binding-incompetent conformation, an unbound binding-competent conformation, testing states, and bound states shown in **Fig. S55**. Simulated *k*_*on*_, *k*_*off*,_ and *K*_*d*_ values across 20 “motifs” of various strengths (from *f*_*motif*_ = 0.99 to 0.01) were obtained by coarsely varying 3 free microscopic rate constants *k*_*on, max*_, *k*_*off, max*_, and *k*_*off, u*_ across 4 orders of magnitude each in 10 step increments, and an additional equilibrium folding constant *K*_*fold*_ (defined as *k*_*on, fold*_/(1-*k*_*on, fold*_)) over 5 increments (spanning percent folded in the unbound state from 1 – 99%). For each combination of free parameters, we simulated 3 independent trajectories with 10^4^ reaction steps. The resulting free energy difference between the tightest and weakest surveyed motifs was calculated to report selectivity.

##### Sensitivity analysis for model with multiple bound conformations

Reaction likelihoods were defined according to a 5-state model of consisting of unbound TF, a selective testing and bound state, and a promiscuous testing and bound state (as shown in **Fig. 6F**). Simulated *k*_*on*_, *k*_*off*,_ and *K*_*d*_ values across 10 “motifs” of various strengths (from *f*_*motif*_ = 0.99 to 0.01) were obtained by coarsely varying 6 free microscopic rate constants *k*_*on, max, s*_, *k*_*off, max, s*_, *k*_*off, u, s*_, *k*_*on, max, p*_, *k*_*off, max, p*_, and *k*_*off, u, p*_ across 4 orders of magnitude each in 5 step increments. For each combination of free parameters, we simulated 3 independent trajectories with 3*10^3^ reaction steps. The resulting free energy difference between the tightest and weakest surveyed motifs was calculated to report on selectivity. For relevant parameter spaces, trajectories were re-simulated with 10^4^ steps across 20 “motifs” with 5 independent replicates.

