## Supplemental Information for "High-throughput thermodynamic and kinetic measurements of transcription factor/DNA mutations reveal how conformational heterogeneity can shape motif selectivity"

2022

### Contents

**1 Supplemental figures** **3**

**2 Supplemental tables** **68**

### 1 Supplemental figures

S5 TF expression levels and expression reproducibility across experimental replicates for binding experiments to oligonucleotides containing the consensus 3'-CACGTG-5' motif. **Left:** Box plots indicating absolute per-chamber expression levels for chambers with (green) and without (grey) printed plasmid for each experimental replicate. **Right:** Scatter plots showing pairwise comparisons of measured fluorescence intensities of surface-immobilized eGFP-tagged mutants across experiments. Grey dashed line indicates linear regression with annotated $r^2$ values; red dashed lines indicate identity line. Solid red lines indicate intensity thresholds used for binary classification of mutants that did and did not express. . . . . . . . . . . . . . . . . . . . . . . . . . . . 17

S6 Calibration curves relating measured fluorescence intensities and DNA concentrations for high-affinity CACGTG consensus sequence binding experiments. **(A)** Chamber calibration curves for representative sample chambers containing 2 MAX constructs (WT and R35L) across 3 experiments showing individual points (colored by device chamber) and associated linear fits (black dashed lines). **(B)** Heatmaps showing linear fit slope as a function of chamber position within device. Slopes vary by approximately 2-fold across the device, with lowest slopes corresponding to outer edges of the microscope field-of-view. **(C)** Goodness-of-fit distributions for calibration curves. 18

S7 Determining mutants at the lower limit of assay detection across experimental replicates with different dynamic ranges. **(A)** Images of device chambers showing measured DNA intensities for two MAX variants and an empty chamber lacking arrayed plasmid, at increasing [DNA]. Image contrast was adjusted uniformly for visibility. **(B)** Distribution of button-bound Cy5 intensities for many device chambers across different DNA concentrations for WT MAX, dead mutant MAX R35L, and empty chambers across three reference sequence binding replicates. Empty chambers lacking arrayed plasmid (and therefore expressed TF) are used to determine the effective lower limit of detection for binding to the 5'- C CACGTG A-3' reference sequence. Here, WT MAX is resolvable from background Cy5 intensity (p-value from repeated measures ANOVA test = [0.0074, 0.0073, 0.0108]), while weakly DNA-binding MAX R35L mutant is not (p-value from repeated measures ANOVA test = [0.1143, 0.0858, 0.0842]). Mutations which were indistinguishable from background in all 3 technical replicates are listed in Table S3. **(C)** Distribution of p-values for repeated measures ANOVA test for each MAX mutation versus button-bound GFP intensities. Weak-binding mutations such as MAX R35L are indistinguishable from background even though they are highly expressed, while low expression of a given TF variant can reduce the ability to resolve bound Cy5 intensity from background. Grey dashed lines indicate intensity thresholds used to identify mutants that did and did not express; red dashed lines indicate significance thresholds used to classify background-resolvable TF mutants. . . . . . . . . . . . . . . . . . . . . . . . . . . . 19
S8 Pairwise comparisons of per-mutant **(A)** $K_d$s (top row) and **(B)** $\Delta\Delta G$s (bottom row) for all TF mutants across 3 experiments for reference DNA sequence 5'-C CACGTG A-3'. Points indicate median affinities (± SEM) for each TF mutant. $\Delta\Delta G$s reflect differences in binding energy relative to WT MAX. Linear fits are indicated by grey dashed lines; identity lines are indicated by red dashed lines. Light grey points indicate mutant $K_d$ measurements that were not statistically significantly different from background binding in all replicates, listed in Table S3. . . . . . . . . . . . . . 20
S9 Comparison of normalized MAX mutant eGFP intensities (y axis) versus measured $K_d$s across 3 technical replicates binding to DNA sequences containing the consensus motif 5'-CACGTG-3'. Markers denote per-chamber fitted $K_d$s and immobilized eGFP intensities; dashed line indicates linear regression with annotated $r^2$ values. . 21
S10 Distribution of the number of experimental replicates per MAX mutant aggregated across devices for the 5'-C CACGTG A-3' oligonucleotide. . . . . . . . . . . . . . . . 21
S11 Measured binding affinities ($K_d$s) for MAX mutations classified by mutation type. **(A)** Measured $K_d$s for all mutations classified by their location within MAX. **(B)** Measured $K_d$s for mutations classified by substitution type. In both panels, black markers indicate binding to the cognate CACGTG motif that is statistically significantly different from that of WT MAX (using Bonferroni-corrected p¡0.05); grey markers denote mutants that are WT-like or indistinguishable from background. . . 22

S12 Measured concentration-dependent binding and associated Langmuir isotherm fits for aligned mutations with differential impacts in Pho4 (bottom) and MAX (top). Each plot shows fluorescence intensity ratios for bound DNA over immobiliized TF normalized to the global fit asymptote $R_{max}$ as a function of soluble DNA concentration; grey markers denote WT proteins and teal and orange markers denote MAX and Pho4 mutants, respectively. Solid lines indicate Langmuir isotherm fits for the median measured $K_d$ across all technical replicates. . . . . . . . . . . . . . . . . . . . 23
S13 Simulated impact of TF mutations altering helical propensity on apparent (measured) $K_d$s for a TF that is mostly unfolded in solution. (**A**) Schematic showing 2-state folding-upon-binding equilibrium simulations for WT TF that is predominantly unfolded in the unbound state. (**B**) Resultant apparent DNA-binding isotherms when WT TF $K_d = 1$ and WT TF $k_u = \frac{0.95}{0.05}$ for WT and a helix-breaking mutant with $\Delta\Delta G_{HP} = 2.0$ kcal/mol. Red dashed line denotes the DNA concentration at which half of the TFs are bound, the definition of the observed $K_d$. **C** Thermodynamic model of expected measured $K_d$ for many mutations altering helical propensity. . . . 24
S14 Simulated impact of TF mutations altering helical propensity on apparent (measured) $K_d$s for a TF that is mostly folded (helical) in solution. (**A**) Schematic showing 2-state folding-upon-binding equilibrium simulations for WT TF that is predominantly folded in the unbound state. (**B**) Resultant apparent DNA-binding isotherms when WT TF $K_d = 1$ and WT TF $k_u = \frac{0.05}{0.95}$ for WT and a helix-breaking mutant with $\Delta\Delta G_{HP} = 2.0$ kcal/mol. Red dashed line denotes the DNA concentration at which half of the TFs are bound, the definition of the observed $K_d$. **C** Thermodynamic model of expected measured $K_d$ for many mutations altering helical propensity. If the WT TF is already predominantly folded at equilibrium, mutations that alter helical propensity do not change measured DNA-binding affinity. . . . . . 25
S15 Heatmaps illustrating folding-and-binding thermodynamic model fit for mutations non-contacting basic regions plotted in **Fig. 2E** for (**A**) MAX and (**B**) Pho4. Color indicates RMSE between model-predicted apparent binding affinity ($log_{10}$ $K_{d,apparent}$) and true measured affinity ($log_{10}$ $K_{d,measured}$)) as a function of intrinsic affinity and fraction folded in solution. Lowest RMSE fit for Pho4 data suggests that WT Pho4 binds DNA with a true DNA-binding $K_d$ of 227 $nM$ and is 81 percent unfolded in the unbound form. Lowest RMSE fit for MAX data suggests that WT MAX binds DNA with a true DNA-binding $K_d$ of 1178 $nM$ and is 1 percent unfolded in the unbound form. . . . . . . . . . . . . . . . . . . . . . . . . . . . . . . . . . 26
S16 TF expression levels and expression reproducibility across experimental replicates for binding experiments to oligonucleotides containing the mutated E-Box motif 3'-**A**ACGTG-5'. **Left:** Box plots indicating absolute per-chamber expression levels for chambers with (green) and without (grey) printed plasmid for each experimental replicate. **Right:** Scatter plots showing pairwise comparisons of measured fluorescence intensities of surface-immobilized eGFP-tagged mutants across experiments. Grey dashed line indicates linear regression with annotated $r^2$ values; red dashed lines indicate identity line. Solid red lines indicate intensity thresholds used to identify mutants that did and did not express. . . . . . . . . . . . . . . . . . . . . . . . . 27

S17 TF expression levels and expression reproducibility across experimental replicates for binding experiments to oligonucleotides containing the mutated E-Box motif 3'-C**G**CGTG-5'. **Left:** Box plots indicating absolute per-chamber expression levels for chambers with (green) and without (grey) printed plasmid for each experimental replicate. **Right:** Scatter plots showing pairwise comparisons of measured fluorescence intensities of surface-immobilized eGFP-tagged mutants across experiments. Grey dashed line indicates linear regression with annotated $r^2$ values; red dashed lines indicate identity line. Solid red lines indicate intensity thresholds used to identify mutants that did and did not express. . . . . . . . . . . . . . . . . . . . . . . . . 27
S18 TF expression levels and expression reproducibility across experimental replicates for binding experiments to oligonucleotides containing the mutated E-Box motif 3'-CA**T**GTG-5'. **Left:** Box plots indicating absolute per-chamber expression levels for chambers with (green) and without (grey) printed plasmid for each experimental replicate. **Right:** Scatter plots showing pairwise comparisons of measured fluorescence intensities of surface-immobilized eGFP-tagged mutants across experiments. Grey dashed line indicates linear regression with annotated $r^2$ values; red dashed lines indicate identity line. Solid red lines indicate intensity thresholds used to identify mutants that did and did not express. . . . . . . . . . . . . . . . . . . . . . . . . 28
S19 TF expression levels and expression reproducibility across experimental replicates for binding experiments to oligonucleotides containing the mutated E-Box motif 3'-CACG**C**G-5'. **Left:** Box plots indicating absolute per-chamber expression levels for chambers with (green) and without (grey) printed plasmid for each experimental replicate. **Right:** Scatter plots showing pairwise comparisons of measured fluorescence intensities of surface-immobilized eGFP-tagged mutants across experiments. Grey dashed line indicates linear regression with annotated $r^2$ values; red dashed lines indicate identity line. Solid red lines indicate intensity thresholds used to identify mutants that did and did not express. . . . . . . . . . . . . . . . . . . . . . . . . 28
S20 TF expression levels and expression reproducibility across experimental replicates for binding experiments to oligonucleotides containing the mutated E-Box motif 3'-CACGT**T**-5'. **Left:** Box plots indicating absolute per-chamber expression levels for chambers with (green) and without (grey) printed plasmid for each experimental replicate. **Right:** Scatter plots showing pairwise comparisons of measured fluorescence intensities of surface-immobilized eGFP-tagged mutants across experiments. Grey dashed line indicates linear regression with annotated $r^2$ values; red dashed lines indicate identity line. Solid red lines indicate intensity thresholds used to identify mutants that did and did not express. . . . . . . . . . . . . . . . . . . . . . . . . 29
S21 Pairwise comparison of per-mutant **(A)** $K_d$s (top row) and **(B)** $\Delta\Delta G$s (bottom row) for all TF mutants across 3 experiments for mutant DNA sequence 5-C **A**ACGTG A-3. Points indicate median affinities (± SEM) for each TF mutant. $\Delta\Delta G$s reflect relative differences in binding energy relative to WT MAX. Linear fits are indicated by grey dashed lines; identity lines are indicated by red dashed lines. Light grey points indicate mutant $K_d$ measurements that are not statistically significantly different from background binding in all replicates, listed in Table S3. . . . . . . . . . 30

S22 Pairwise comparison of per-mutant **(A)** $K_d$s (top row) and **(B)** $\Delta\Delta G$s (bottom row) for all TF mutants across 3 experiments for mutant DNA sequence 5'-C C**G**CGTG A-3'. Points indicate median affinities (± SEM) for each TF mutant. $\Delta\Delta G$s reflect relative differences in binding energy relative to WT MAX. Linear fits are indicated by grey dashed lines; identity lines are indicated by red dashed lines. Light grey points indicate mutant $K_d$ measurements that are not statistically significantly different from background binding in all replicates, listed in Table S3. . . . . . . . . . . 31
S23 Pairwise comparison of per-mutant **(A)** $K_d$s (top row) and **(B)** $\Delta\Delta G$s (bottom row) for all TF mutants across 3 experiments for mutant DNA sequence 5'-C CA**T**GTG A-3'. Points indicate median affinities (± SEM) for each TF mutant. $\Delta\Delta G$s reflect relative differences in binding energy relative to WT MAX. Linear fits are indicated by grey dashed lines; identity lines are indicated by red dashed lines. Light grey points indicate mutant $K_d$ measurements that are not statistically significantly different from background binding in all replicates, listed in Table S3. . . . . . . . . . . 32
S24 Pairwise comparison of per-mutant **(A)** $K_d$s (top row) and **(B)** $\Delta\Delta G$s (bottom row) for all TF mutants across 3 experiments for mutant DNA sequence 5'-C CACG**C**G A-3'. Points indicate median affinities (± SEM) for each TF mutant. $\Delta\Delta G$s reflect relative differences in binding energy relative to WT MAX. Linear fits are indicated by grey dashed lines; identity lines are indicated by red dashed lines. Light grey points indicate mutant $K_d$ measurements that are not statistically significantly different from background binding in all replicates, listed in Table S3. . . . . . . . . . . 33
S25 Pairwise comparison of per-mutant **(A)** $K_d$s (top row) and **(B)** $\Delta\Delta G$s (bottom row) for all TF mutants across 2 experiments for mutant DNA sequence 5'-C CACGT**T** A-3'. Points indicate median affinities (± SEM) for each TF mutant. $\Delta\Delta G$s reflect relative differences in binding energy relative to WT MAX. Linear fits are indicated by grey dashed lines; identity lines are indicated by red dashed lines. Light grey points indicate mutant $K_d$ measurements that are not statistically significantly different from background binding in all replicates, listed in Table S3. . . . . . . . . . . 34
S26 Comparison of normalized MAX mutant eGFP intensities (y axis) versus measured $K_d$s across 3 technical replicates binding to DNA sequences containing the mutations to the mutated E-box motifs **(A)** 5'-**A**ACGTG-3', **(B)** 5'-C**G**CGTG-3', **(C)** 5'-CA**T**GTG-3', **(D)** 5'-CACG**C**G-3', and **(E)** 5'-CACGT**T**-3'. Markers denote per-chamber fitted $K_d$s and immobilized eGFP intensities; dashed line indicates linear regression with annotated $r^2$ values. . . . . . . . . . . . . . . . . . . . . . . . . . . . . 35
S27 Distribution of the number of experimental replicates per MAX mutant aggregated across devices for the binding experiments conducted with oligonucleotides containing mutations to the consensus motif 5'-C CACGTG A-3'. . . . . . . . . . . . . . . . . . . 36
S28 Median $K_d$s for all MAX mutations sorted by linear residue position within the MAX sequence for all measured oligonucleotides. Points indicate median affinities (± SEM) for each TF mutant + DNA binding interaction. . . . . . . . . . . . . . . . . . . . . . 37

S29 Classification of affinity-altering substitutions in MAX. **Left:** Histogram of variance in $\Delta\Delta G_{peroligo}$ between all measured oligonucleotides. Red line indicates upper bound for variance of affinity altering mutations (variance $\leq 0.04$), excluding mutations with the highest quartile of variance in measured $\Delta\Delta G$s. **Right:** Mean $\Delta\Delta G_{peroligo}$ versus variance in $\Delta\Delta G_{peroligo}$ for all MAX mutations. Red markers indicate putative "affinity altering" mutations. Light grey points indicate mutations un-resolvable from background in at least one oligonucleotide. . . . . . . . . . . . . 38
S30 Pairwise comparison between measured $\Delta\Delta G$s for MAX mutants interacting with a low-affinity mutant sequence (5′-C **A**ACGTG A-3′) versus the reference sequence (5′-C CACGTG A-3′). Each marker indicates the median $\Delta\Delta G$ ($\pm$ SEM) for a given TF mutant across all chambers in all replicates. Grey dashed line indicates linear regression $y = 0.42 * x + 0.17$; red dashed line indicates identity line. Markers corresponding to TF mutations to known crystallographic contacts to nucleotide bases (H28 and E32) are colored in red. Non-additive mutations in $K_d$ space are indicated by red outlines; see **Table S6**. . . . . . . . . . . . . . . . . . . . . . . . . . . 39
S31 Pairwise comparison between measured binding affinities for MAX mutants interacting with a low-affinity mutant sequence (5′-C C**G**CGTG A-3′) versus the reference sequence (5′-C CACGTG A-3′). Each marker indicates the median $\Delta\Delta G$ ($\pm$ SEM) for a given TF mutant across all chambers in all replicates. Grey dashed line indicates linear regression $y = 0.62 * x + 0.19$; red dashed line indicates identity line. Markers corresponding to TF mutations to known crystallographic contacts to nucleotide bases (H28 and E32) are colored in red. Non-additive mutations in $K_d$ space are indicated by red outlines; see **Table S6**. . . . . . . . . . . . . . . . . . . . . . . . . . 40
S32 Pairwise comparison between measured binding affinities for MAX mutants interacting with a low-affinity mutant sequence (5′-C CA**T**GTG A-3′) versus the reference sequence (5′-C CACGTG A-3′). Each marker indicates the median $\Delta\Delta G$ ($\pm$ SEM) for a given TF mutant across all chambers in all replicates. Grey dashed line indicates linear regression $y = 0.71 * x + 0.02$; red dashed line indicates identity line. Markers corresponding to TF mutations to known crystallographic contacts to nucleotide bases (R36) are colored in red. Non-additive mutations in $K_d$ space are indicated by red outlines; see **Table S6**. . . . . . . . . . . . . . . . . . . . . . . . . . . 41
S33 Pairwise comparison between measured binding affinities for MAX mutants interacting with a low-affinity mutant sequence (5′-C CACGT**T** A-3′) versus the reference sequence (5′-C CACGTG A-3′). Each marker indicates the median $\Delta\Delta G$ ($\pm$ SEM) for a given TF mutant across all chambers in all replicates. Grey dashed line indicates linear regression $y = 0.58 * x + 0.19$; red dashed line indicates identity line. Markers corresponding to TF mutations to known crystallographic contacts to nucleotide bases (H28 and E32) are colored in red. Non-additive mutations in $K_d$ space are indicated by red outlines; see **Table S6**. . . . . . . . . . . . . . . . . . . . . . . . . . 42
S34 Mutations that alter sequence specificity such that a motif besides CACGTG is lowest in affinity. Bar plots indicate $\Delta\Delta G$ ($\pm$ SEM) for each mutation defined as $K_{d,mutant+mutantE-Box}$ relative to the $K_{d,mutant+CACGTG}$. Asterisks indicate significance (p-value $< 0.05$ in independent T-test comparing measured $K_d$s for the MAX mutant binding CACGTG versus the MAX mutant binding alternate motifs). 43

S35 Residual Z-score for each consensus oligonucleotide versus mutant oligonucleotide double mutant cycle comparison as a function of MAX amino acid sequence. Each marker indicates the residual Z-score of a MAX mutation computed from a double mutant cycle comparing $\Delta\Delta G$ of a mutated E-box motif to the cognate motif. Marker colors indicate the identity of the mutated oligonucleotide in the double mutant cycle comparison. . . . . . . . . . . . . . . . . . . . . . . . . . . . . . . . . . . . . . . . . 44
S36 Bar plots indicate $\Delta\Delta G$ ($\pm$ SEM) for each selective mutation relative to WT MAX in different oligonucleotide binding interactions. Asterisks indicate non-additive binding relative to WT MAX and CACGTG. Binding interactions that are unresolvable from background are annotated with red arrows, indicating that the plotted $\Delta\Delta G$ represents a lower bound estimate of the true difference in binding energies. . . . . 45
S37 Definition and location within structure of specificity-increasing MAX mutations, listed in **Figure S36**. **A** Histogram of median residual Z-scores for all measured MAX mutations, with threshold for determining specificity altering mutations indicated by the dashed red line (|median residual Z-score|$>$ 0.9). **B** Location of specificity increasing mutations (median residual Z-score $>$ 0.9) indicated on the MAX structure (PDB ID: 1HLO) in red. . . . . . . . . . . . . . . . . . . . . . . . . . . . . . . 46
S38 Median $K_d$ $\pm$ SEM across all E-box motifs for WT and all measured substitutions at H27. . . . . . . . . . . . . . . . . . . . . . . . . . . . . . . . . . . . . . . . . . . . 47
S39 **(A)** Median residual Z-score ("specificity score") versus $\Delta\Delta G_{CACGTG}$ for MAX (teal) and Pho4 (orange). Each point represents one TF mutation within the basic region. Size of marker indicates the relative conservation of the mutated residue. **(B)** Quantification of distribution of mutations in each quadrant for both MAX (teal) and Pho4 (orange). **(C)** Relative affinity (to WT) and measured $K_d$s for comparable MAX and Pho4 mutations. Markers indicate $K_d$s (median $\pm$ SEM) for WT TF (grey), MAX A30V (teal) and Pho4 H257V (orange). **(D)** Relative affinity (to WT) and measured $K_d$s for comparable MAX and Pho4 mutations. Markers indicate $K_d$s (median $\pm$ SEM) for WT TF (grey), MAX H27A (teal) and Pho4 S254A (orange). . 48
S40 **(A)** Mean energetic impact of a TF mutation on DNA binding across all measured DNA sequences ($\langle\Delta\Delta G_{DNA}\rangle$) versus variance in the energetic impact of a TF mutation on DNA binding across all measured DNA sequences (variance of $\Delta\Delta G_{DNA}$). Each point represents a unique Pho4 mutation. Dotted lines indicate 2*SEM difference from WT $\langle\Delta\Delta G_{DNA}\rangle$ for all oligonucleotides. Red points are mutations that can be confidently called as affinity-altering mutations within the measurement variance, defined as mutations where(variance of $\Delta\Delta G_{DNA}$) is less than $\langle\Delta\Delta G_{DNA}\rangle$. **(B)** Mutations where the absolute value of $\langle\Delta\Delta G_{DNA}\rangle > 0.5$ kcal/mol, colored in red on the Pho4 crystal structure (PDB ID: 1A0A). . . . . . . . . . . . . . . . . . . 49
S41 Pairwise comparisons of per-mutant $K_d$s for a small library of MAX (teal) and Pho4 (orange) mutants across 2 experiments. Points indicate median affinities ($\pm$ SEM) for each TF mutant. Linear fits are indicated by grey dashed lines; identity lines are indicated by red dashed lines. Light grey points indicate mutant $K_d$ measurements that are not statistically significantly different from background binding in all replicates, listed in Table S3. . . . . . . . . . . . . . . . . . . . . . . . . . . . . . . . 50
S42 Expected additive $\Delta\Delta G$ (calculated as summed $\Delta\Delta G$ of each mutation alone) versus measured $\Delta\Delta G$ of MAX H27V/A30G double mutant across all E-box motifs. . . . . 51

S43 Experimental workflow illustrating iterative trapping and dissociation of fluorescently-labelled DNA to quantify binding off-rates with imaging steps indicated. The addition of high-affinity unlabeled competitor DNA during dissociation prevents rebinding. 52
S44 Pairwise comparisons of per-mutant **A** $K_d$s and **B** apparent $\Delta\Delta G$s for Pho4 mutants across 3 experiments. Markers indicate values (median ± SEM) for each mutant. Linear fits are indicated by grey dashed lines; identity lines are indicated by red dashed lines. . . . . 53
S45 Pairwise comparisons of per-mutant **A** $K_d$s and **B** apparent $\Delta\Delta G$s for MAX mutants across 3 experiments. Markers indicate rates (median ± SEM) for each mutant. Linear fits are indicated by grey dashed lines; identity lines are indicated by red dashed lines. Light grey points indicate mutant $K_d$ measurements that are not statistically significantly different from background binding in all replicates, listed in Table S3. . . . . 54
S46 Pairwise comparisons of per-mutant **A** $k_{off}$s and **B** apparent $k_{on}$s for Pho4 mutants across 3 experiments. Markers indicate rates (median ± SEM) for each mutant. Linear fits are indicated by grey dashed lines; identity lines are indicated by red dashed lines. . . . . 55
S47 Pairwise comparisons of per-mutant **A** $k_{off}$s and **B** apparent $k_{on}$s for MAX mutants across 3 experiments. Markers indicate rates (median ± SEM) for each mutant. Linear fits are indicated by grey dashed lines; identity lines are indicated by red dashed lines. Light grey points indicate mutant $K_d$ measurements that are not statistically significantly different from background binding in all replicates, listed in Table S3. Identities of select outlier mutations are annotated. . . . . 56
S48 Pairwise comparisons of per-mutant $k_{off}$s for a small library of MAX (teal) and Pho4 (orange) mutants across 2 experiments. Markers indicate off-rates (median ± SEM) for each TF mutant. Linear fits are indicated by grey dashed lines; identity lines are indicated by red dashed lines. Light grey points indicate mutant $K_d$ measurements that are not statistically significantly different from background binding in all replicates, listed in Table S3. . . . . 57
S49 Pairwise comparisons of per-mutant apparent $k_{on}$s for a small library of MAX (teal) and Pho4 (orange) mutants across 2 experiments. Markers indicate on-rates (median ± SEM) for each TF mutant. Linear fits are indicated by grey dashed lines; identity lines are indicated by red dashed lines. Light grey points indicate mutant $K_d$ measurements that are not statistically significantly different from background binding in all replicates, listed in Table S3. . . . . 58
S50 Values (median ± SEM) of measured off- (top) and on-rate (bottom) constants for selective MAX mutants interacting with many E-box variants. Mutations shown in **(A)** speed up off-rate for all DNA sequences, mutations in **(B)** speed up on-rate mostly for the cognate motif, and the MAX mutation in **(C)** exhibits both behaviors. 59

S51 Ternary heat maps demonstrating that the 3-state model with one binding conformation shown in **6D** cannot explain differences in intrinsic selectivity for the same set of sequences. Simulated binding affinities for the most preferred sequence (defined as $f_{motif} = 0.99$) **(A)** and least preferred sequence (defined as $f_{motif} = 0.01$) **(B)** define binding selectivity **(C)**), the free energy difference between binding. All simulated affinities and free energies are shown as a function of microscopic rate constants $k_{off,\mu}$, $k_{off,M}$, and $k_{on,max}$. Each value represents the median across 3 simulation trajectories. . . . . . . . . . . . . . . . . . . . . . . . . . . . . . . . . . . . . . . 59
S52 Example schematic showing 4-state model and associated rate constants describing folding-and-binding with a single binding-competent folded conformation. . . . . . . 60
S53 Ternary heat maps demonstrating that the 4-state model with one binding conformation shown in **S52** cannot explain differences in intrinsic selectivity for the same set of sequences. Each ternary heat map shows selectivity (the energetic difference between the most ($f_{motif} = 0.99$) and least ($f_{motif} = 0.01$) preferred sequences) as a function of microscopic rate constants $k_{off,\mu}$, $k_{off,M}$, and $k_{on,max}$, varying the degree of pre-folded structure in the unbound state (shown in order of least to most residual structure, **(A)** $\approx 25$, **(B)** $\approx 50$, **(C)** $\approx 75$, and **(D)** $\approx 99$ percent folded) between each plot. Each value represents the median across 3 simulation trajectories. 61
S54 Ternary heat maps establishing that the 5-state model with distinct conformations model shown in **6F** can explain differences in intrinsic selectivity for the same set of sequences. Each ternary heat map shows selectivity (the energetic difference between the most ($f_{motif} = 0.99$) and least ($f_{motif} = 0.01$) preferred sequences) as a function of microscopic rate constants $k_{off,\mu,s}$, $k_{off,M,s}$, $k_{on,max,s}$, $k_{off,\mu,p}$, and $k_{off,M,p}$. Values on all ternary heat maps represent medians across 3 simulation trajectories when $k_{on,max,p} = 3 * 10^3$. . . . . . . . . . . . . . . . . . . . . . . . . . . . . . . . 62
S55 Ternary heat maps establishing that the 5-state model with distinct conformations model shown in **6F** can explain differences in intrinsic selectivity for the same set of sequences. Each ternary heat map shows selectivity (the energetic difference between the most ($f_{motif} = 0.99$) and least ($f_{motif} = 0.01$) preferred sequences) as a function of microscopic rate constants $k_{off,\mu,s}$, $k_{off,M,s}$, $k_{on,max,s}$, $k_{off,\mu,p}$, and $k_{off,M,p}$. Values on all ternary heat maps represent medians across 3 simulation trajectories when $k_{on,max,p} = 3 * 10^4$. . . . . . . . . . . . . . . . . . . . . . . . . . . . . . . . 63
S56 Ternary heat maps establishing that the 5-state model with distinct conformations model shown in **6F** can explain differences in intrinsic selectivity for the same set of sequences. Each ternary heat map shows selectivity (the energetic difference between the most ($f_{motif} = 0.99$) and least ($f_{motif} = 0.01$) preferred sequences) as a function of microscopic rate constants $k_{off,\mu,s}$, $k_{off,M,s}$, $k_{on,max,s}$, $k_{off,\mu,p}$, and $k_{off,M,p}$. Values on all ternary heat maps represent medians across 3 simulation trajectories when $k_{on,max,p} = 3 * 10^5$. . . . . . . . . . . . . . . . . . . . . . . . . . . . . . . . 64
S57 Ternary heat maps establishing that the 5-state model with distinct conformations model shown in **6F** can explain differences in intrinsic selectivity for the same set of sequences. Each ternary heat map shows selectivity (the energetic difference between the most ($f_{motif} = 0.99$) and least ($f_{motif} = 0.01$) preferred sequences) as a function of microscopic rate constants $k_{off,\mu,s}$, $k_{off,M,s}$, $k_{on,max,s}$, $k_{off,\mu,p}$, and $k_{off,M,p}$. Values on all ternary heat maps represent medians across 3 simulation trajectories when $k_{on,max,p} = 3 * 10^6$. . . . . . . . . . . . . . . . . . . . . . . . . . . . . . . . 65

S58 Ternary heat maps establishing that the 5-state model with distinct conformations model shown in **6F** can explain differences in intrinsic selectivity for the same set of sequences. Each ternary heat map shows selectivity (the energetic difference between the most ($f_{motif} = 0.99$) and least ($f_{motif} = 0.01$) preferred sequences) as a function of microscopic rate constants $k_{off,\mu,s}$, $k_{off,M,s}$, $k_{on,max,s}$, $k_{off,\mu,p}$, and $k_{off,M,p}$. Values on all ternary heat maps represent medians across 3 simulation trajectories when $k_{on,max,p} = 3 * 10^7$. . . . . . . . . . . . . . . . . . . . . . . . . . . . . . . . 66
S59 Co-varying $k_{on,max,specific}$ and $k_{off,\mu,specific}$ can recapitulate measured MAX A30G data in both kinetic (**A**), affinity (**B**), and free energy (**C**) space. . . . . . . . . . . 67

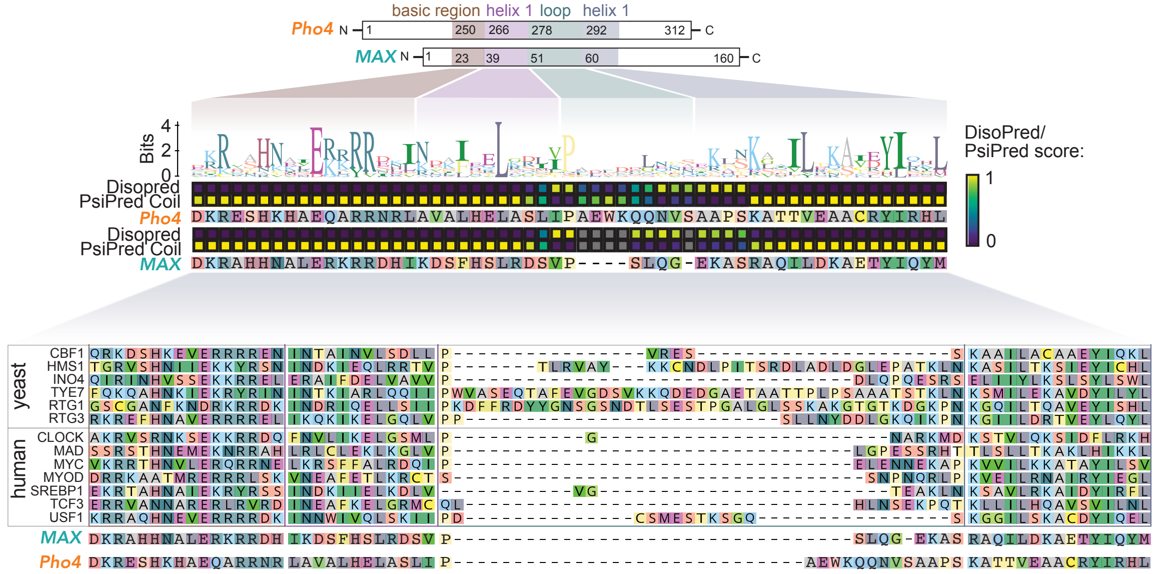

Figure S1: Protein sequence alignment for bHLH proteins across yeast and humans (bottom), with predicted secondary structure and domain organization for model bHLH TFs Pho4 and MAX annotated (top).

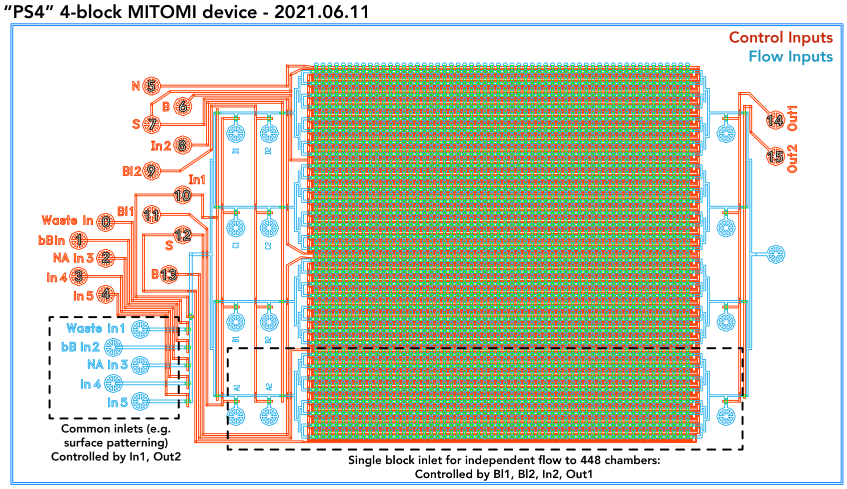

Figure S2: Schematic of 2-layer microfluidic device with 4 blocks of chambers with independently accessible inlet and outlet ports. Control channels are shown in red, flow channels are shown in blue, and inlet ports are labeled by their function.

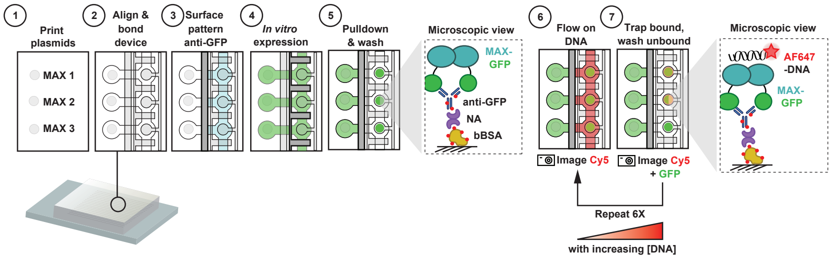

Figure S3: Experimental workflow illustrating device alignment, surface chemistry to pattern antibody for subsequent TF immobilization, expression of GFP-tagged TFs via *in vitro* transcription and translation, adsorption and purification of TF variants, and introduction of DNA at multiple concentrations to quantify binding with iterative imaging steps indicated.

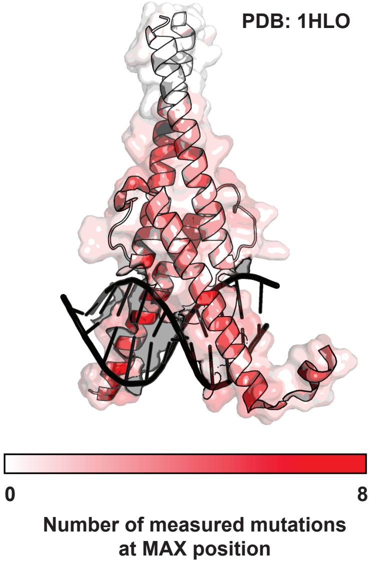

Figure S4: Number of measured mutations at each position in MAX mutational library superimposed on DNA-bound MAX crystal structure (1HLO).

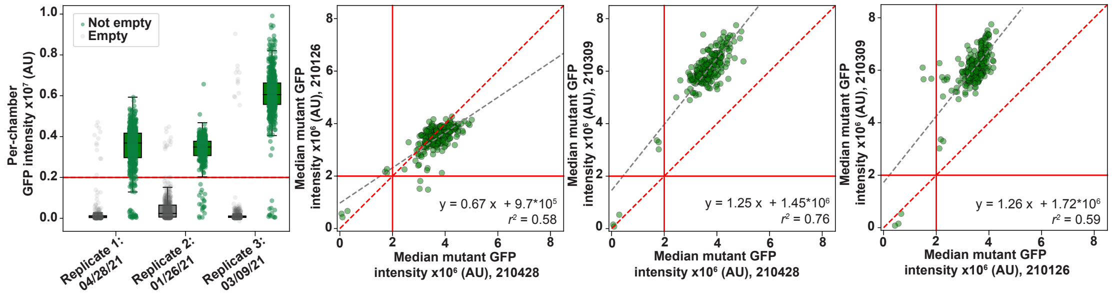

Figure S5: TF expression levels and expression reproducibility across experimental replicates for binding experiments to oligonucleotides containing the consensus 3'-CACGTG-5' motif. **Left:** Box plots indicating absolute per-chamber expression levels for chambers with (green) and without (grey) printed plasmid for each experimental replicate. **Right:** Scatter plots showing pairwise comparisons of measured fluorescence intensities of surface-immobilized eGFP-tagged mutants across experiments. Grey dashed line indicates linear regression with annotated $r^2$ values; red dashed lines indicate identity line. Solid red lines indicate intensity thresholds used for binary classification of mutants that did and did not express.

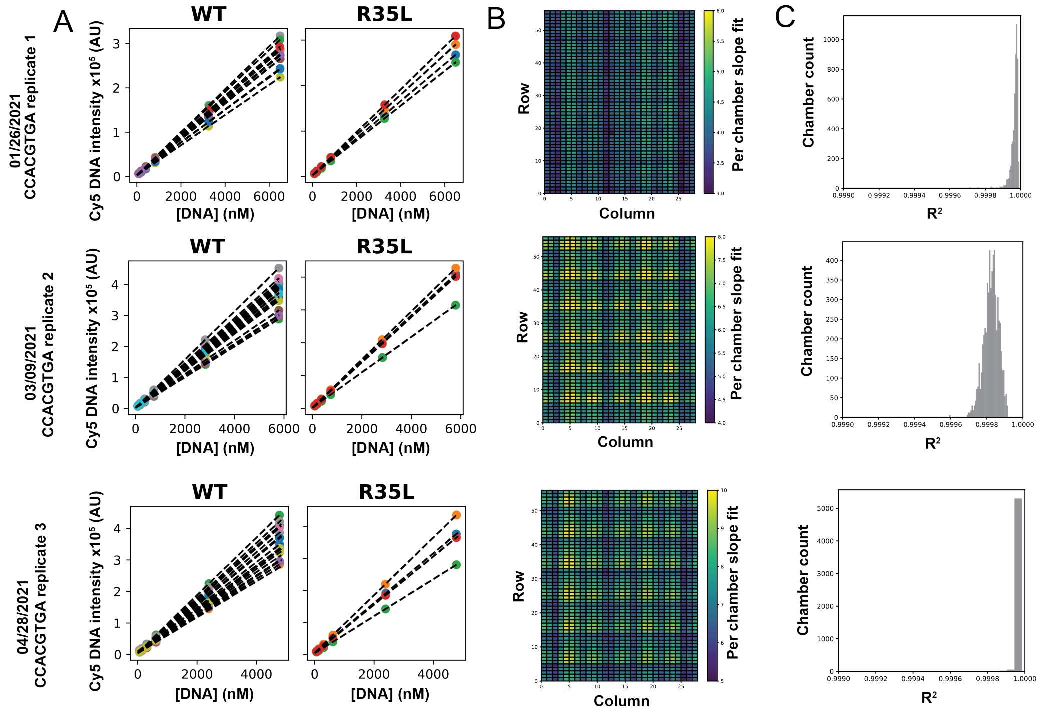

Figure S6: Calibration curves relating measured fluorescence intensities and DNA concentrations for high-affinity CACGTG consensus sequence binding experiments. **(A)** Chamber calibration curves for representative sample chambers containing 2 MAX constructs (WT and R35L) across 3 experiments showing individual points (colored by device chamber) and associated linear fits (black dashed lines). **(B)** Heatmaps showing linear fit slope as a function of chamber position within device. Slopes vary by approximately 2-fold across the device, with lowest slopes corresponding to outer edges of the microscope field-of-view. **(C)** Goodness-of-fit distributions for calibration curves.

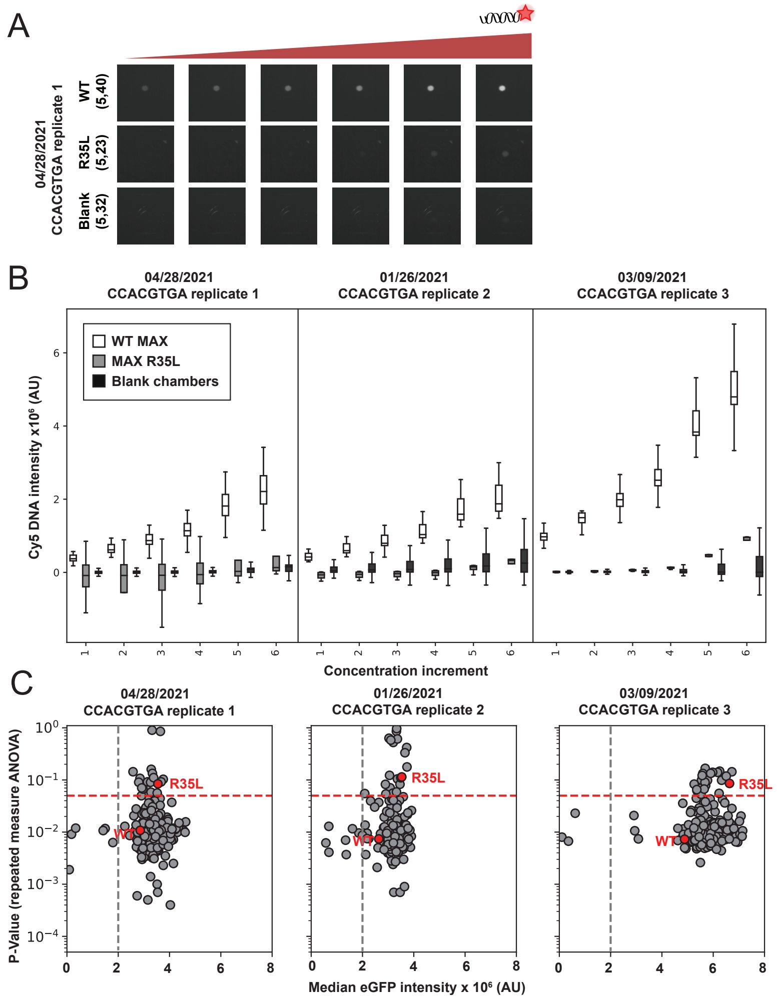

Figure S7: Determining mutants at the lower limit of assay detection across experimental replicates with different dynamic ranges. **(A)** Images of device chambers showing measured DNA intensities for two MAX variants and an empty chamber lacking arrayed plasmid, at increasing [DNA]. Image contrast was adjusted uniformly for visibility. **(B)** Distribution of button-bound Cy5 intensities for many device chambers across different DNA concentrations for WT MAX, dead mutant MAX R35L, and empty chambers across three reference sequence binding replicates. Empty chambers lacking arrayed plasmid (and therefore expressed TF) are used to determine the effective lower limit of detection for binding to the 5'- C CACGTG A-3' reference sequence. Here, WT MAX is resolvable from background Cy5 intensity (p-value from repeated measures ANOVA test = [0.0074, 0.0073, 0.0108]), while weakly DNA-binding MAX R35L mutant is not (p-value from repeated measures ANOVA test = [0.1143, 0.0858, 0.0842]). Mutations which were indistinguishable from background in all 3 technical replicates are listed in Table S3. **(C)** Distribution of p-values for repeated measures ANOVA test for each MAX mutation versus button-bound GFP intensities. Weak-binding mutations such as MAX R35L are indistinguishable from background even though they are highly expressed, while low expression of a given TF variant can reduce the ability to resolve bound Cy5 intensity from background. Grey dashed lines indicate intensity thresholds used to identify mutants that did and did not express; red dashed lines indicate significance thresholds used to classify background-resolvable TF mutants.

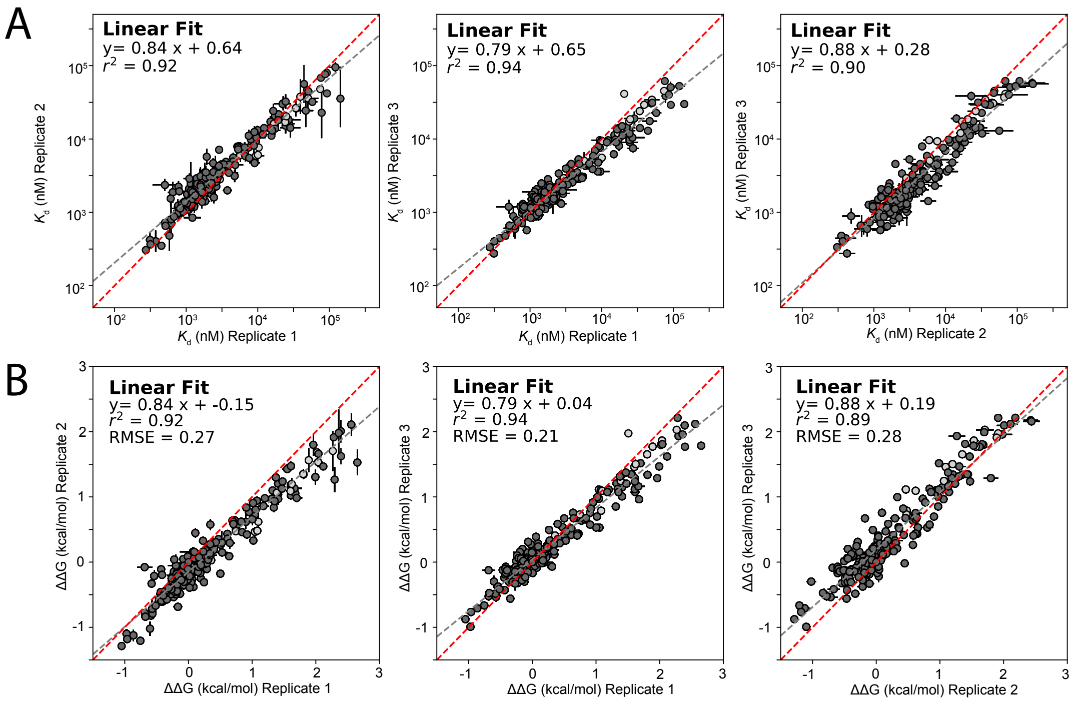

Figure S8: Pairwise comparisons of per-mutant **(A)** $K_d$s (top row) and **(B)** $\Delta\Delta G$s (bottom row) for all TF mutants across 3 experiments for reference DNA sequence 5'-C CACGTG A-3'. Points indicate median affinities (± SEM) for each TF mutant. $\Delta\Delta G$s reflect differences in binding energy relative to WT MAX. Linear fits are indicated by grey dashed lines; identity lines are indicated by red dashed lines. Light grey points indicate mutant $K_d$ measurements that were not statistically significantly different from background binding in all replicates, listed in Table S3.

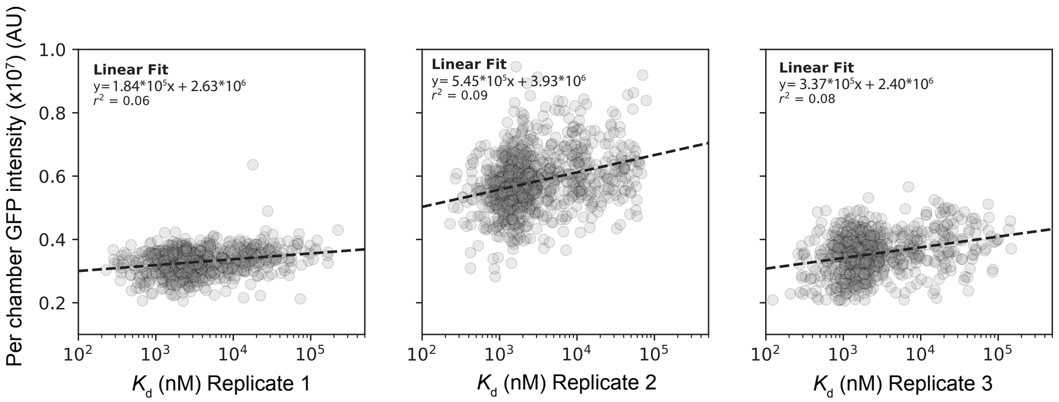

Figure S9: Comparison of normalized MAX mutant eGFP intensities (y axis) versus measured $K_d$s across 3 technical replicates binding to DNA sequences containing the consensus motif 5'-CACGTG-3'. Markers denote per-chamber fitted $K_d$s and immobilized eGFP intensities; dashed line indicates linear regression with annotated $r^2$ values.

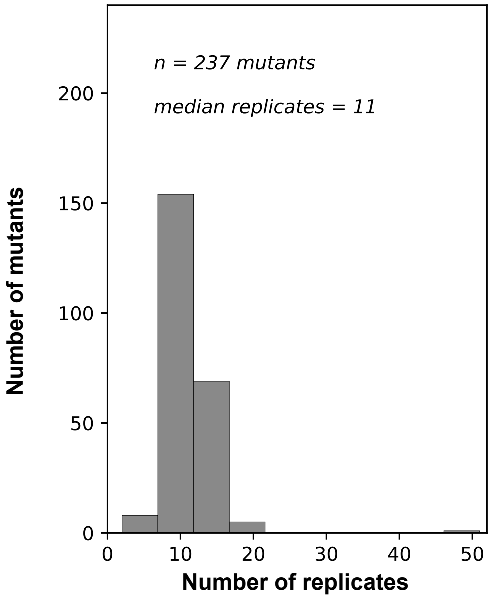

Figure S10: Distribution of the number of experimental replicates per MAX mutant aggregated across devices for the 5'-C CACGTG A-3' oligonucleotide.

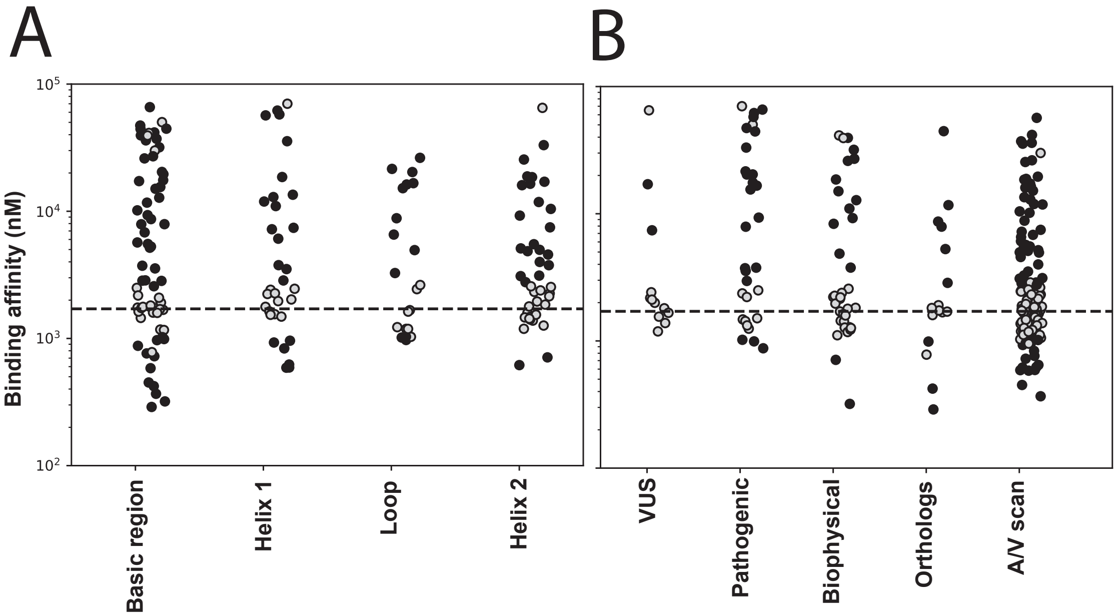

Figure S11: Measured binding affinities ($K_d$s) for MAX mutations classified by mutation type. **(A)** Measured $K_d$s for all mutations classified by their location within MAX. **(B)** Measured $K_d$s for mutations classified by substitution type. In both panels, black markers indicate binding to the cognate CACGTG motif that is statistically significantly different from that of WT MAX (using Bonferroni-corrected p¡0.05); grey markers denote mutants that are WT-like or indistinguishable from background.

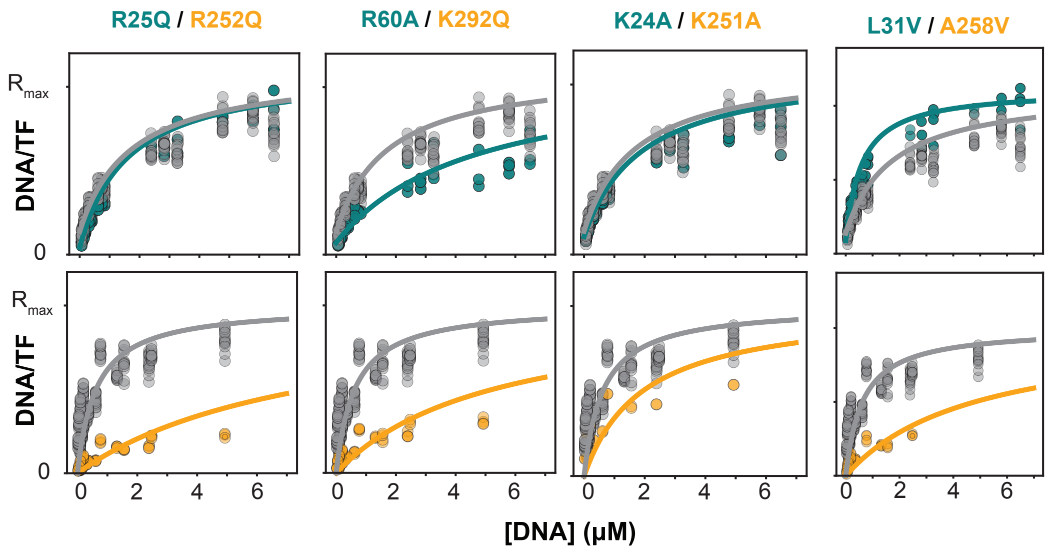

Figure S12: Measured concentration-dependent binding and associated Langmuir isotherm fits for aligned mutations with differential impacts in Pho4 (bottom) and MAX (top). Each plot shows fluorescence intensity ratios for bound DNA over immobiliized TF normalized to the global fit asymptote $R_{max}$ as a function of soluble DNA concentration; grey markers denote WT proteins and teal and orange markers denote MAX and Pho4 mutants, respectively. Solid lines indicate Langmuir isotherm fits for the median measured $K_d$ across all technical replicates.

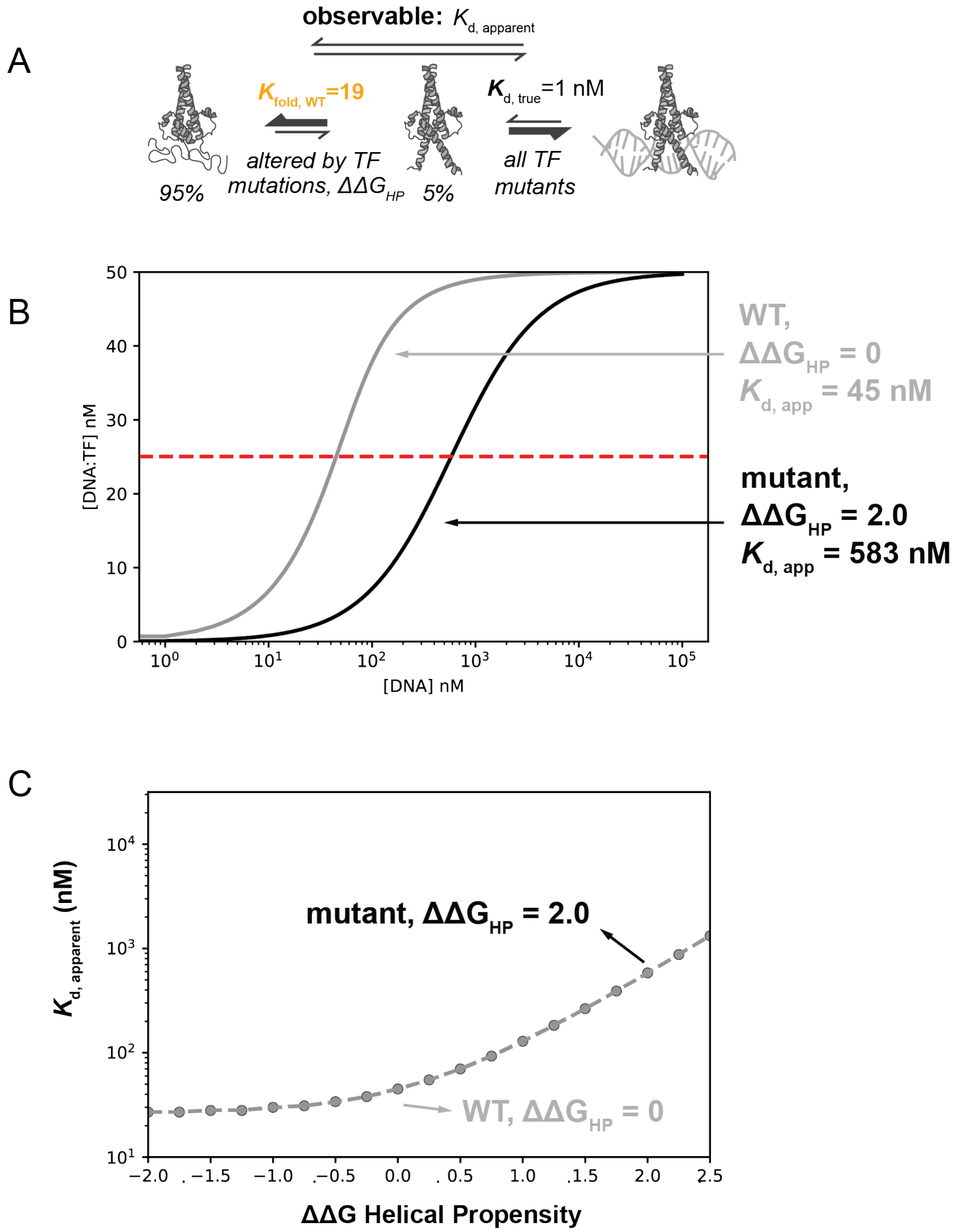

Figure S13: Simulated impact of TF mutations altering helical propensity on apparent (measured) $K_d$s for a TF that is mostly unfolded in solution. (**A**) Schematic showing 2-state folding-upon-binding equilibrium simulations for WT TF that is predominantly unfolded in the unbound state. (**B**) Resultant apparent DNA-binding isotherms when WT TF $K_d = 1$ and WT TF $k_u = \frac{0.95}{0.05}$ for WT and a helix-breaking mutant with $\Delta\Delta G_{HP} = 2.0$ kcal/mol. Red dashed line denotes the DNA concentration at which half of the TFs are bound, the definition of the observed $K_d$. **C** Thermodynamic model of expected measured $K_d$ for many mutations altering helical propensity.

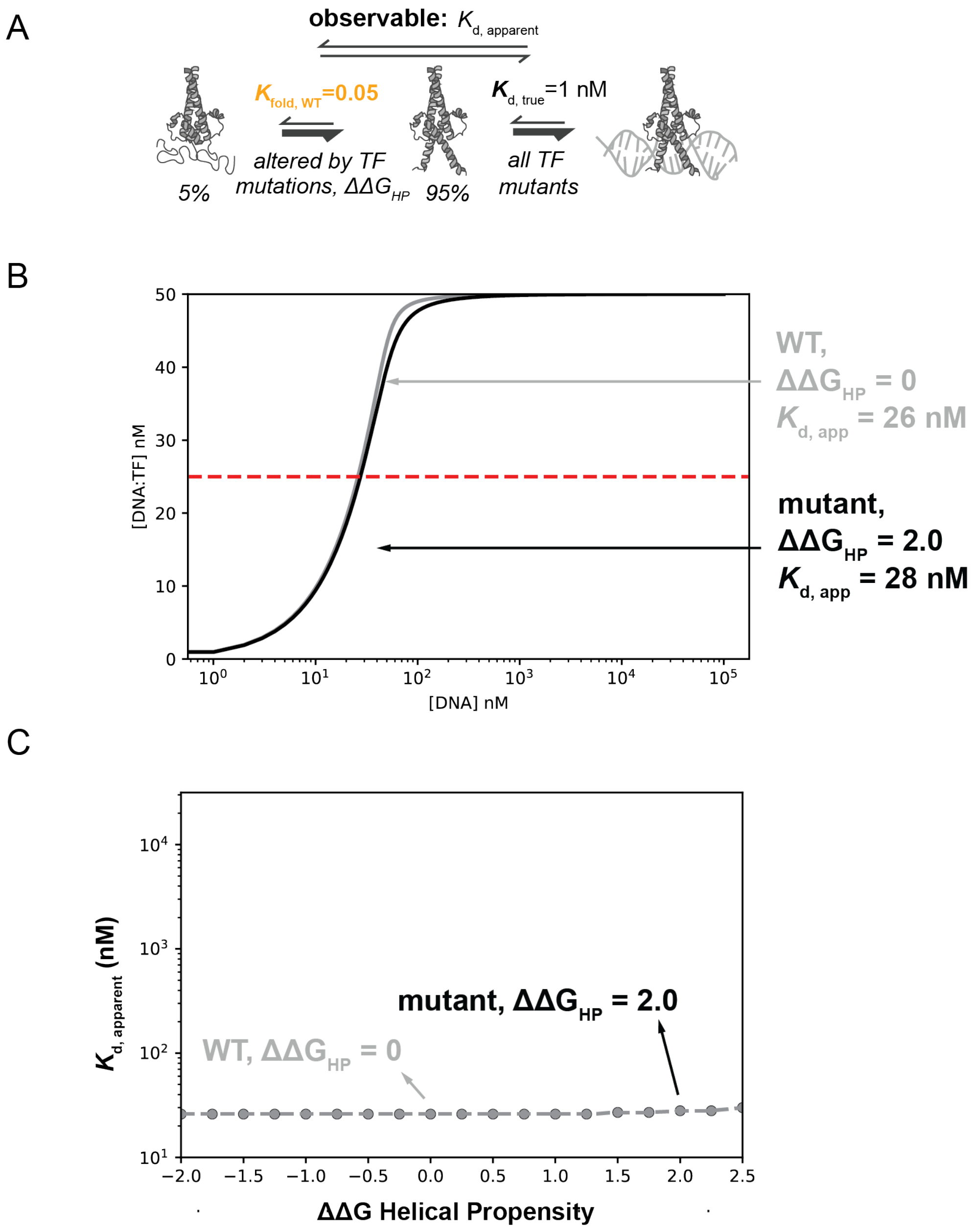

Figure S14: Simulated impact of TF mutations altering helical propensity on apparent (measured) $K_d$s for a TF that is mostly folded (helical) in solution. (**A**) Schematic showing 2-state folding-upon-binding equilibrium simulations for WT TF that is predominantly folded in the unbound state. (**B**) Resultant apparent DNA-binding isotherms when WT TF $K_d = 1$ and WT TF $k_u = \frac{0.05}{0.95}$ for WT and a helix-breaking mutant with $\Delta\Delta G_{HP} = 2.0$ kcal/mol. Red dashed line denotes the DNA concentration at which half of the TFs are bound, the definition of the observed $K_d$. **C** Thermodynamic model of expected measured $K_d$ for many mutations altering helical propensity. If the WT TF is already predominantly folded at equilibrium, mutations that alter helical propensity do not change measured DNA-binding affinity.

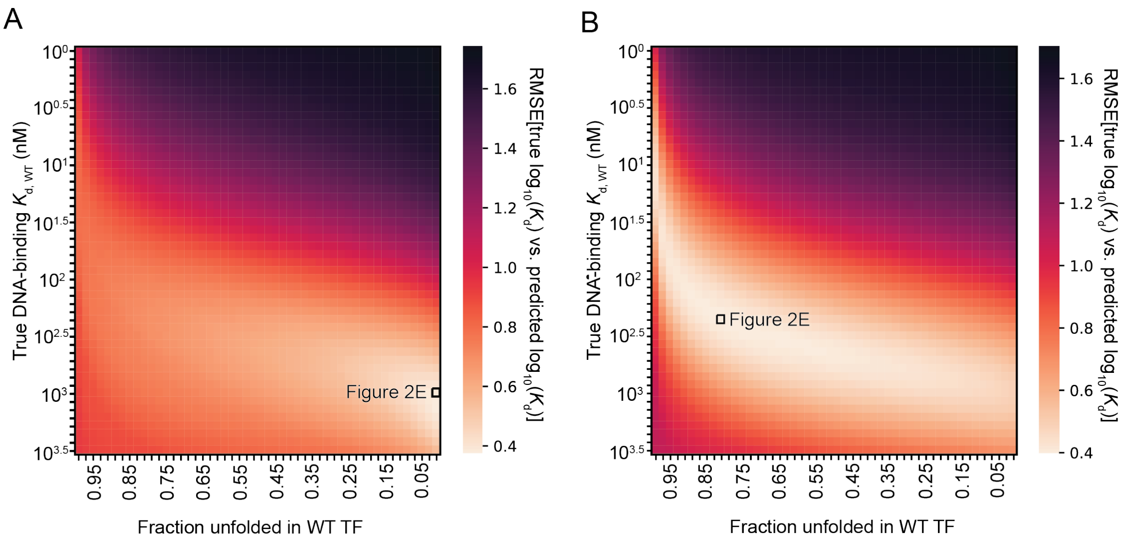

Figure S15: Heatmaps illustrating folding-and-binding thermodynamic model fit for mutations non-contacting basic regions plotted in **Fig. 2E** for (**A**) MAX and (**B**) Pho4. Color indicates RMSE between model-predicted apparent binding affinity ($log_{10}\ K_{d,apparent}$) and true measured affinity ($log_{10}\ K_{d,measured}$)) as a function of intrinsic affinity and fraction folded in solution. Lowest RMSE fit for Pho4 data suggests that WT Pho4 binds DNA with a true DNA-binding $K_d$ of 227 $nM$ and is 81 percent unfolded in the unbound form. Lowest RMSE fit for MAX data suggests that WT MAX binds DNA with a true DNA-binding $K_d$ of 1178 $nM$ and is 1 percent unfolded in the unbound form.

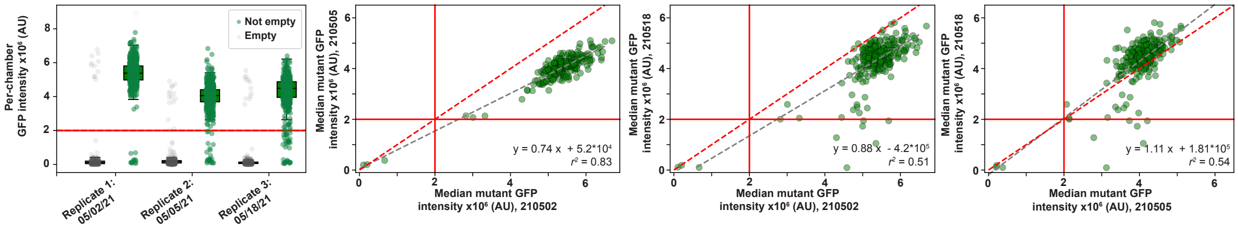

Figure S16: TF expression levels and expression reproducibility across experimental replicates for binding experiments to oligonucleotides containing the mutated E-Box motif 3'-**A**ACGTG-5'. **Left:** Box plots indicating absolute per-chamber expression levels for chambers with (green) and without (grey) printed plasmid for each experimental replicate. **Right:** Scatter plots showing pairwise comparisons of measured fluorescence intensities of surface-immobilized eGFP-tagged mutants across experiments. Grey dashed line indicates linear regression with annotated $r^2$ values; red dashed lines indicate identity line. Solid red lines indicate intensity thresholds used to identify mutants that did and did not express.

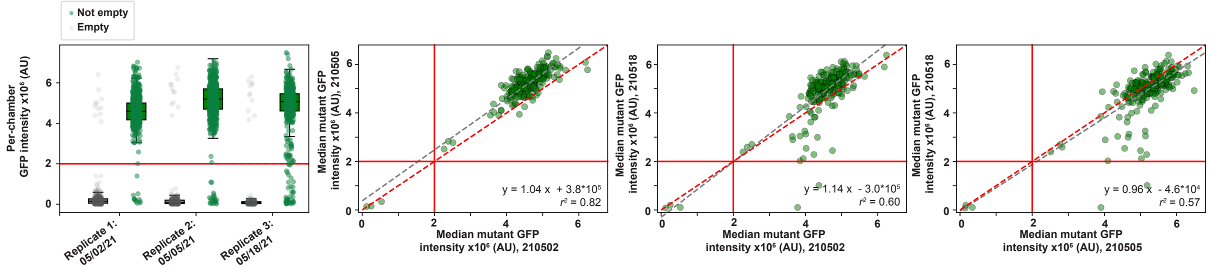

Figure S17: TF expression levels and expression reproducibility across experimental replicates for binding experiments to oligonucleotides containing the mutated E-Box motif 3'-C**G**CGTG-5'. **Left:** Box plots indicating absolute per-chamber expression levels for chambers with (green) and without (grey) printed plasmid for each experimental replicate. **Right:** Scatter plots showing pairwise comparisons of measured fluorescence intensities of surface-immobilized eGFP-tagged mutants across experiments. Grey dashed line indicates linear regression with annotated $r^2$ values; red dashed lines indicate identity line. Solid red lines indicate intensity thresholds used to identify mutants that did and did not express.

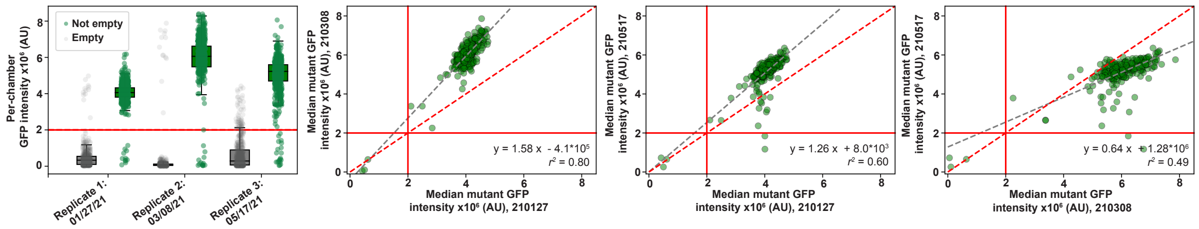

Figure S18: TF expression levels and expression reproducibility across experimental replicates for binding experiments to oligonucleotides containing the mutated E-Box motif 3'-CA**T**GTG-5'. **Left:** Box plots indicating absolute per-chamber expression levels for chambers with (green) and without (grey) printed plasmid for each experimental replicate. **Right:** Scatter plots showing pairwise comparisons of measured fluorescence intensities of surface-immobilized eGFP-tagged mutants across experiments. Grey dashed line indicates linear regression with annotated $r^2$ values; red dashed lines indicate identity line. Solid red lines indicate intensity thresholds used to identify mutants that did and did not express.

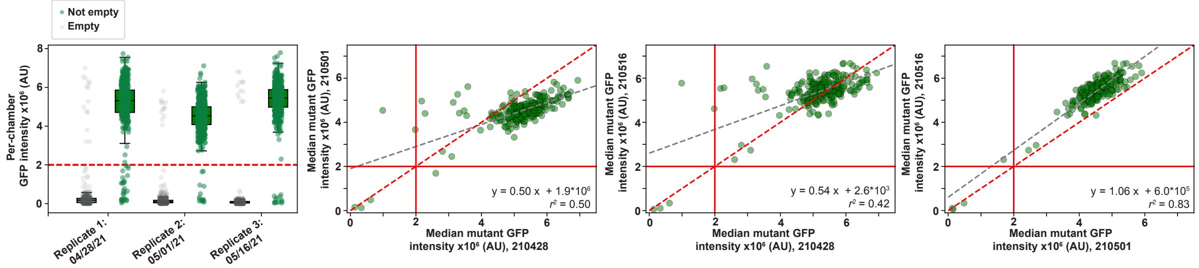

Figure S19: TF expression levels and expression reproducibility across experimental replicates for binding experiments to oligonucleotides containing the mutated E-Box motif 3'-CACG**C**G-5'. **Left:** Box plots indicating absolute per-chamber expression levels for chambers with (green) and without (grey) printed plasmid for each experimental replicate. **Right:** Scatter plots showing pairwise comparisons of measured fluorescence intensities of surface-immobilized eGFP-tagged mutants across experiments. Grey dashed line indicates linear regression with annotated $r^2$ values; red dashed lines indicate identity line. Solid red lines indicate intensity thresholds used to identify mutants that did and did not express.

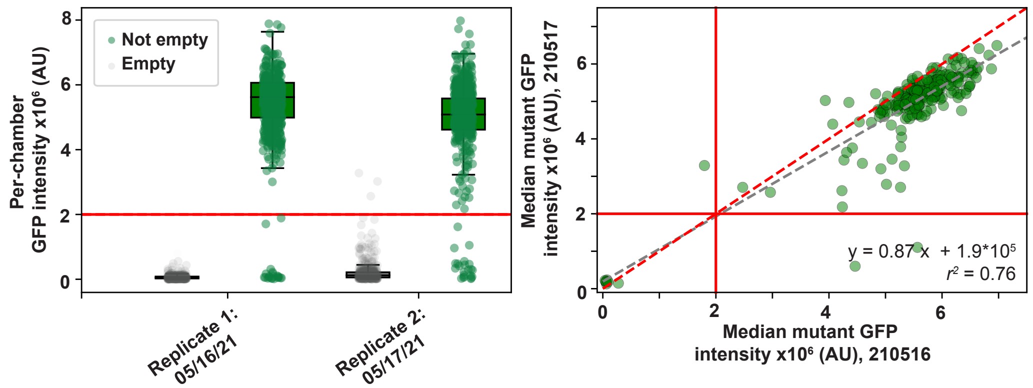

Figure S20: TF expression levels and expression reproducibility across experimental replicates for binding experiments to oligonucleotides containing the mutated E-Box motif 3'-CACGT**T**-5'. **Left:** Box plots indicating absolute per-chamber expression levels for chambers with (green) and without (grey) printed plasmid for each experimental replicate. **Right:** Scatter plots showing pairwise comparisons of measured fluorescence intensities of surface-immobilized eGFP-tagged mutants across experiments. Grey dashed line indicates linear regression with annotated $r^2$ values; red dashed lines indicate identity line. Solid red lines indicate intensity thresholds used to identify mutants that did and did not express.

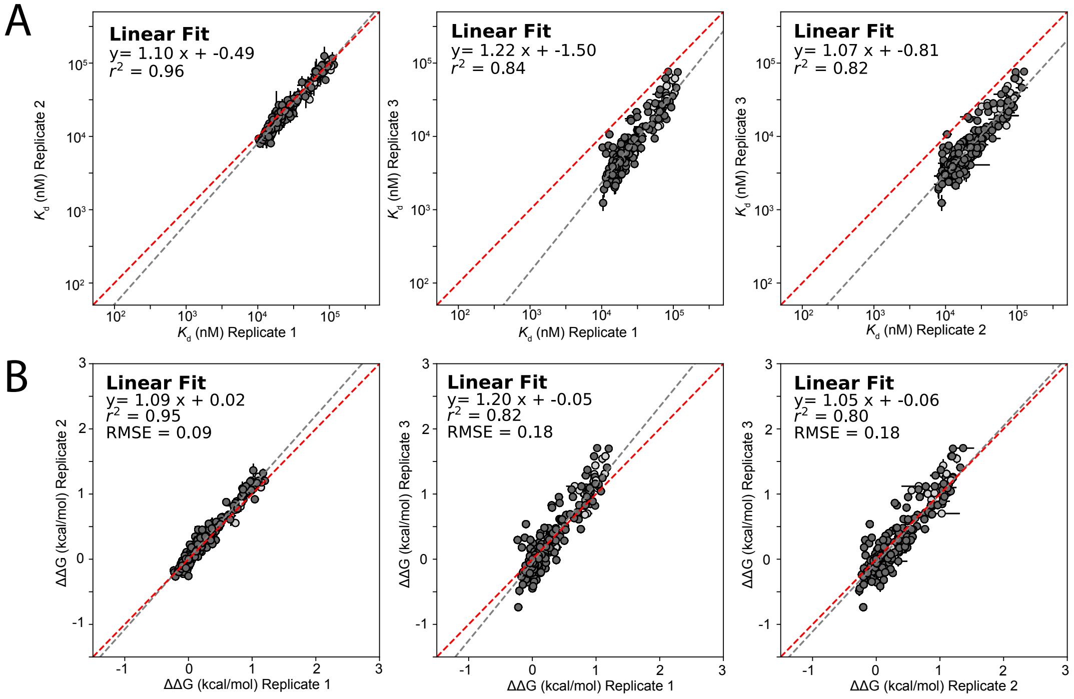

Figure S21: Pairwise comparison of per-mutant **(A)** $K_d$s (top row) and **(B)** $\Delta\Delta G$s (bottom row) for all TF mutants across 3 experiments for mutant DNA sequence 5-C **A**ACGTG A-3. Points indicate median affinities (± SEM) for each TF mutant. $\Delta\Delta G$s reflect relative differences in binding energy relative to WT MAX. Linear fits are indicated by grey dashed lines; identity lines are indicated by red dashed lines. Light grey points indicate mutant $K_d$ measurements that are not statistically significantly different from background binding in all replicates, listed in Table S3.

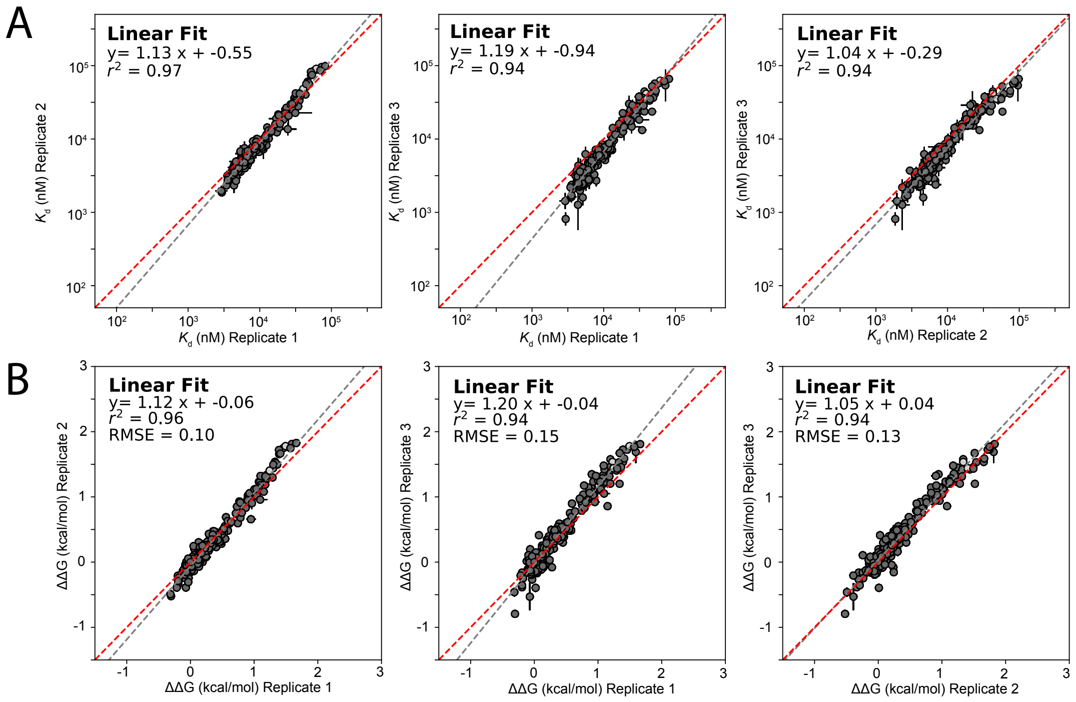

Figure S22: Pairwise comparison of per-mutant **(A)** $K_d$s (top row) and **(B)** $\Delta\Delta G$s (bottom row) for all TF mutants across 3 experiments for mutant DNA sequence 5'-C C**G**CGTG A-3'. Points indicate median affinities (± SEM) for each TF mutant. $\Delta\Delta G$s reflect relative differences in binding energy relative to WT MAX. Linear fits are indicated by grey dashed lines; identity lines are indicated by red dashed lines. Light grey points indicate mutant $K_d$ measurements that are not statistically significantly different from background binding in all replicates, listed in Table S3.

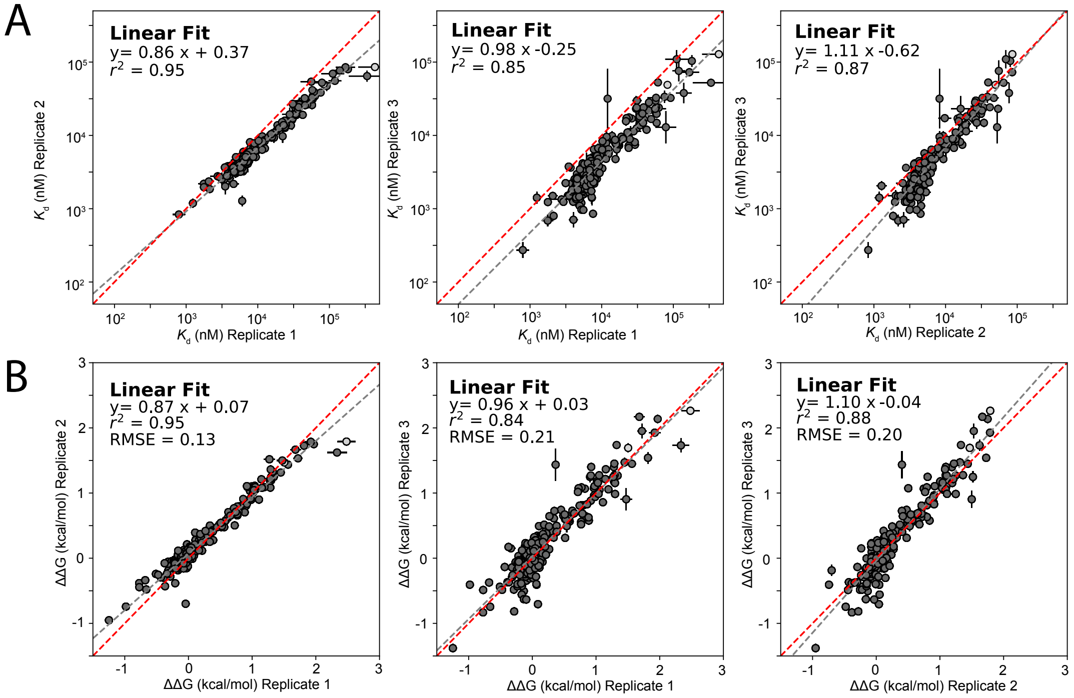

Figure S23: Pairwise comparison of per-mutant **(A)** $K_d$s (top row) and **(B)** $\Delta\Delta G$s (bottom row) for all TF mutants across 3 experiments for mutant DNA sequence 5'-C CA**T**GTG A-3'. Points indicate median affinities (± SEM) for each TF mutant. $\Delta\Delta G$s reflect relative differences in binding energy relative to WT MAX. Linear fits are indicated by grey dashed lines; identity lines are indicated by red dashed lines. Light grey points indicate mutant $K_d$ measurements that are not statistically significantly different from background binding in all replicates, listed in Table S3.

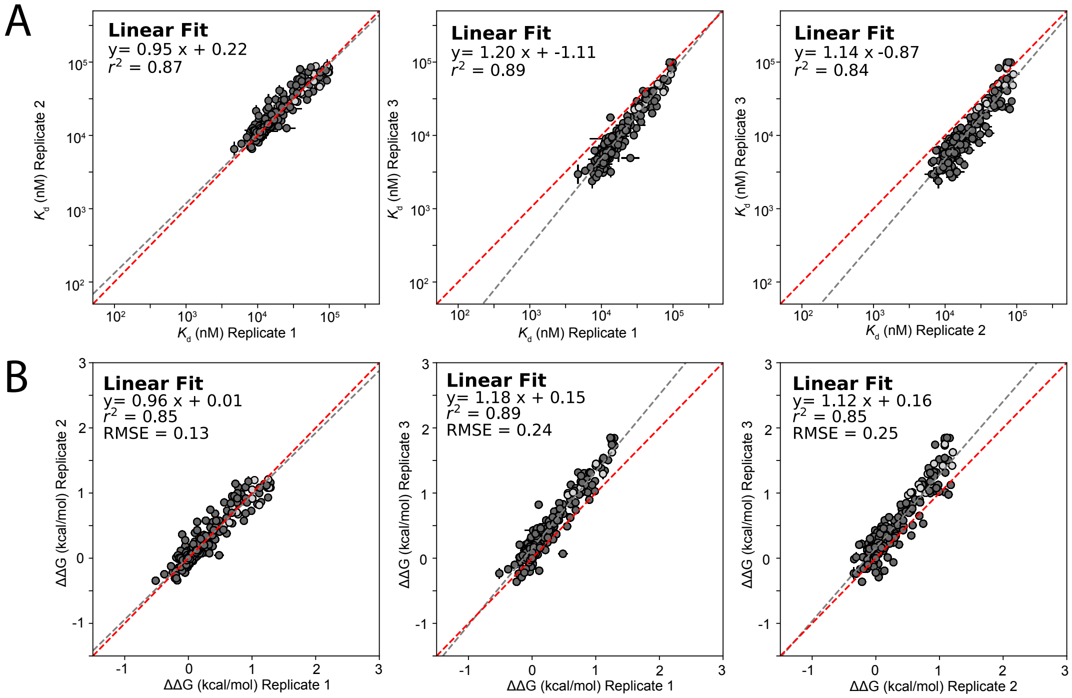

Figure S24: Pairwise comparison of per-mutant **(A)** $K_d$s (top row) and **(B)** $\Delta\Delta G$s (bottom row) for all TF mutants across 3 experiments for mutant DNA sequence 5'-C CACG**C**G A-3'. Points indicate median affinities ($\pm$ SEM) for each TF mutant. $\Delta\Delta G$s reflect relative differences in binding energy relative to WT MAX. Linear fits are indicated by grey dashed lines; identity lines are indicated by red dashed lines. Light grey points indicate mutant $K_d$ measurements that are not statistically significantly different from background binding in all replicates, listed in Table S3.

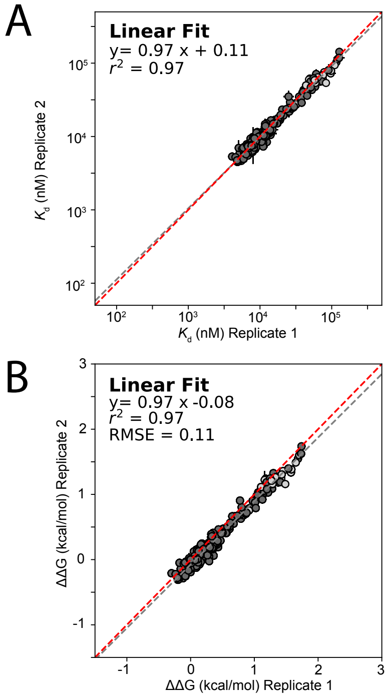

Figure S25: Pairwise comparison of per-mutant **(A)** $K_d$s (top row) and **(B)** $\Delta\Delta G$s (bottom row) for all TF mutants across 2 experiments for mutant DNA sequence 5'-C CACGT**T** A-3'. Points indicate median affinities (± SEM) for each TF mutant. $\Delta\Delta G$s reflect relative differences in binding energy relative to WT MAX. Linear fits are indicated by grey dashed lines; identity lines are indicated by red dashed lines. Light grey points indicate mutant $K_d$ measurements that are not statistically significantly different from background binding in all replicates, listed in Table S3.

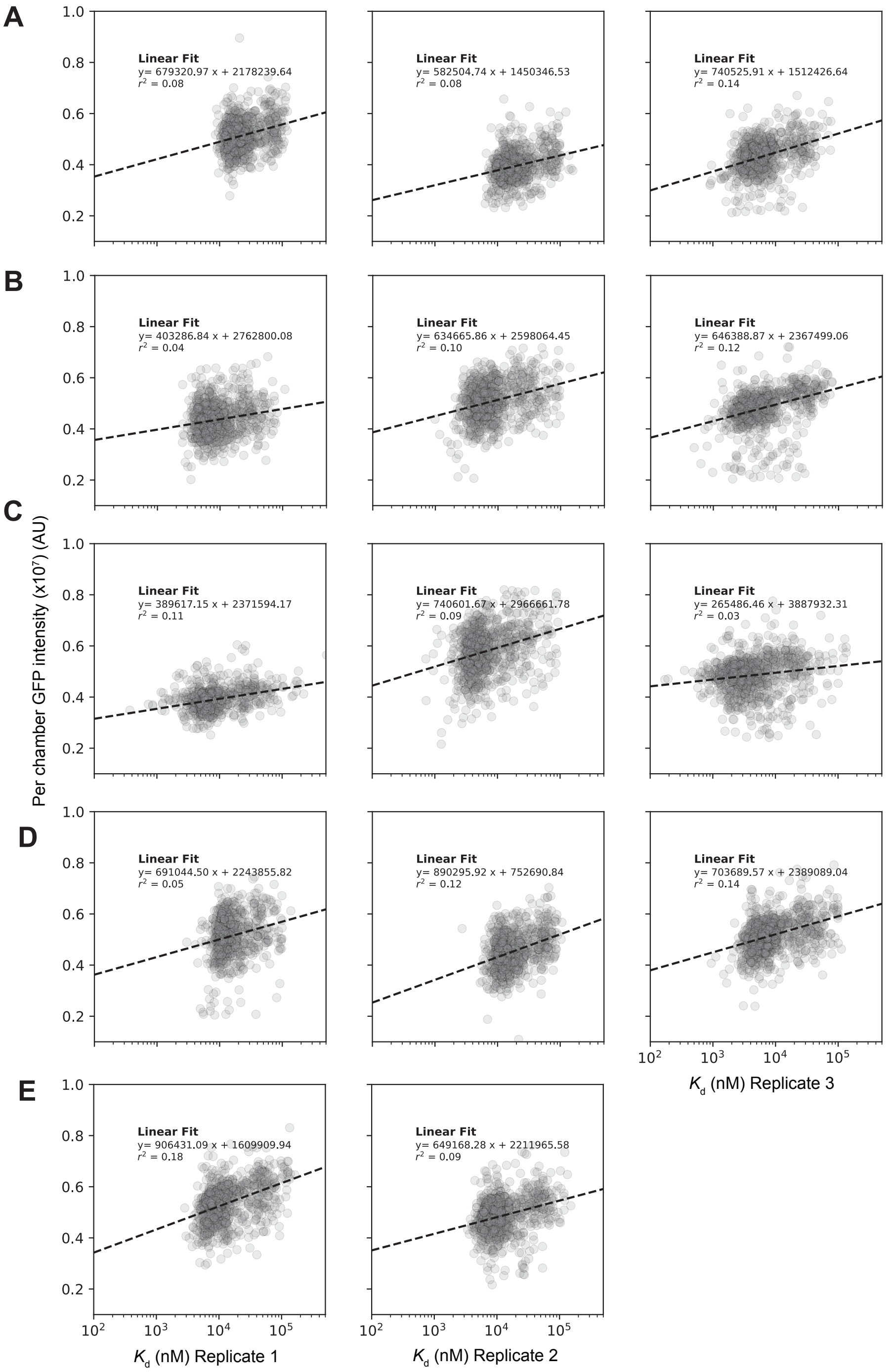

Figure S26: Comparison of normalized MAX mutant eGFP intensities (y axis) versus measured $K_d$s across 3 technical replicates binding to DNA sequences containing the mutations to the mutated E-box motifs **(A)** 5'-**A**ACGTG-3', **(B)** 5'-C**G**CGTG-3', **(C)** 5'-CA**T**GTG-3', **(D)** 5'-CACG**C**G-3', and **(E)** 5'-CACGT**T**-3'. Markers denote per-chamber fitted $K_d$s and immobilized eGFP intensities; dashed line indicates linear regression with annotated $r^2$ values.

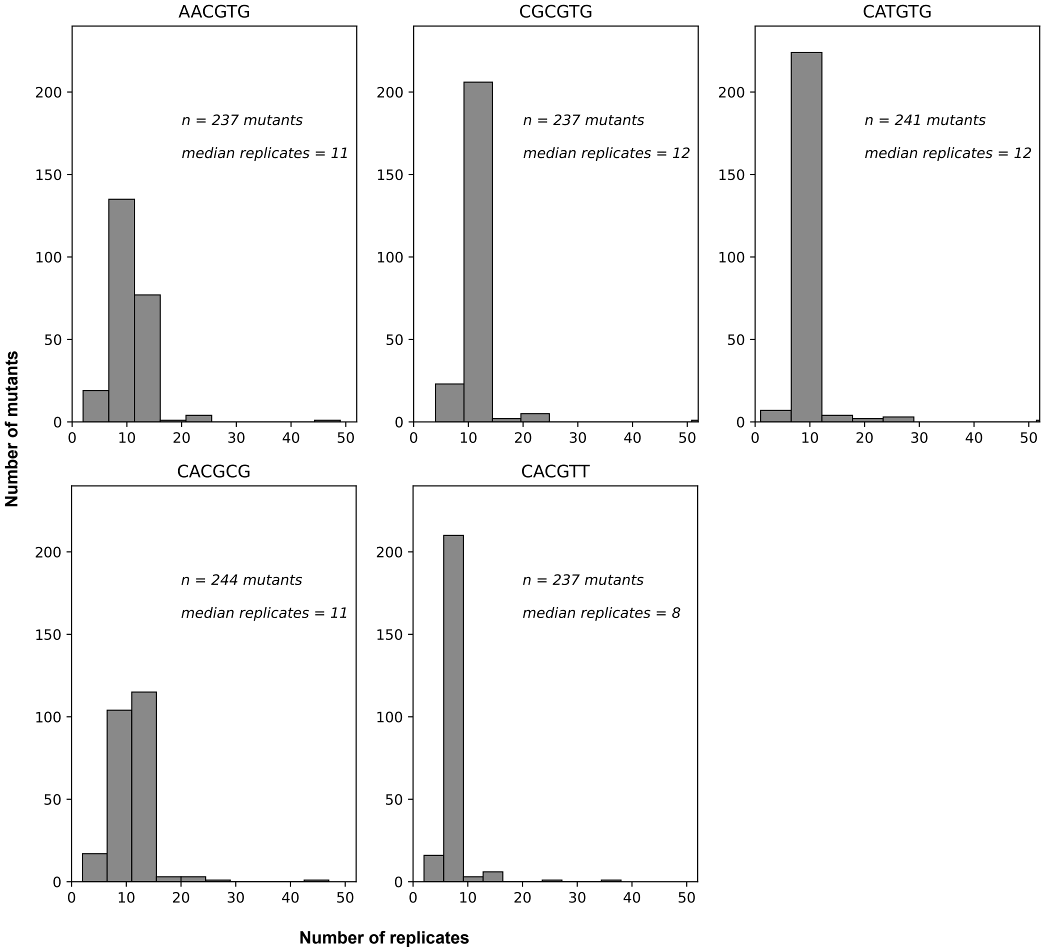

Figure S27: Distribution of the number of experimental replicates per MAX mutant aggregated across devices for the binding experiments conducted with oligonucleotides containing mutations to the consensus motif 5'-C CACGTG A-3'.

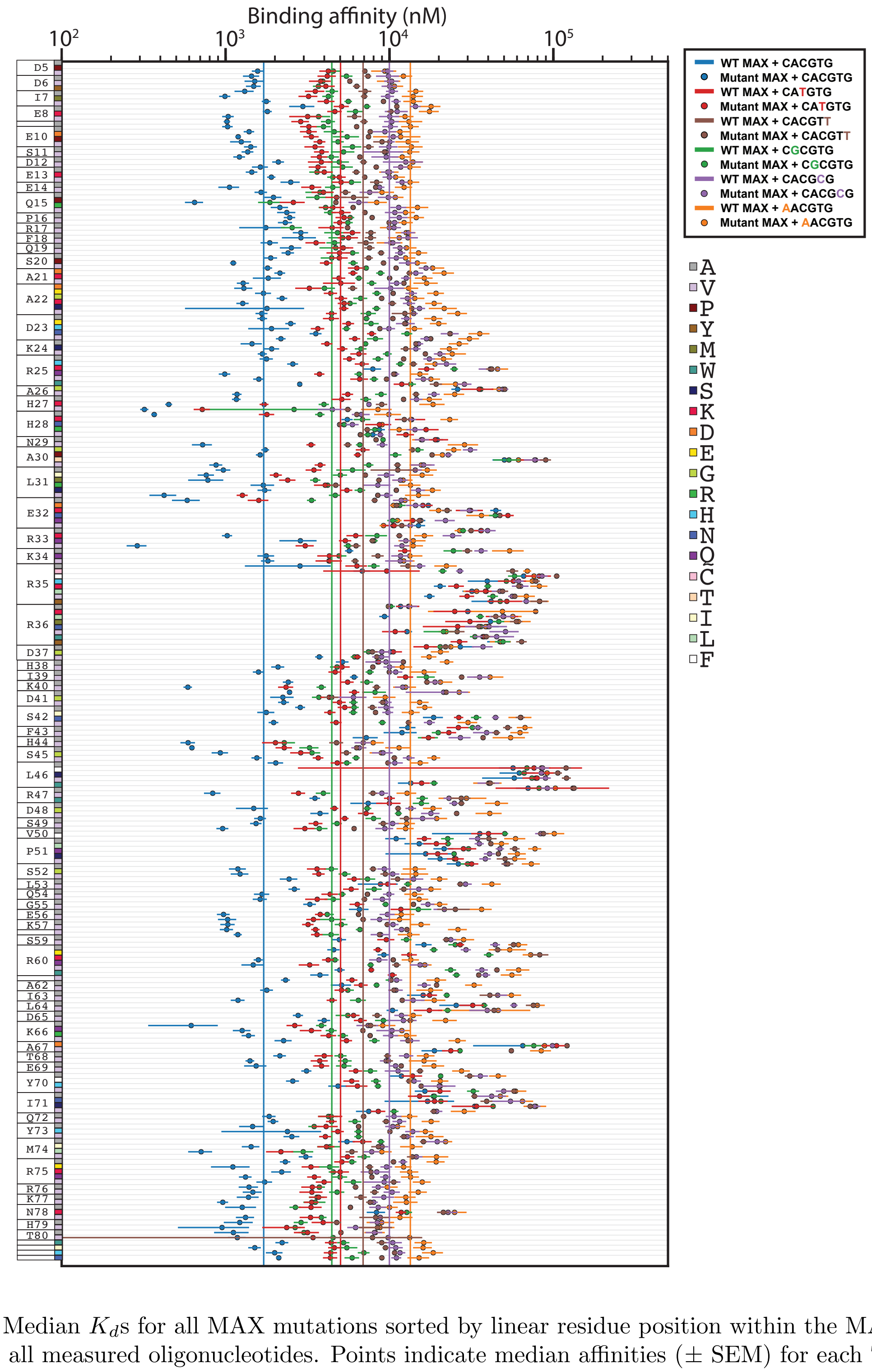

Figure S28: Median $K_d$s for all MAX mutations sorted by linear residue position within the MAX sequence for all measured oligonucleotides. Points indicate median affinities (± SEM) for each TF mutant + DNA binding interaction.

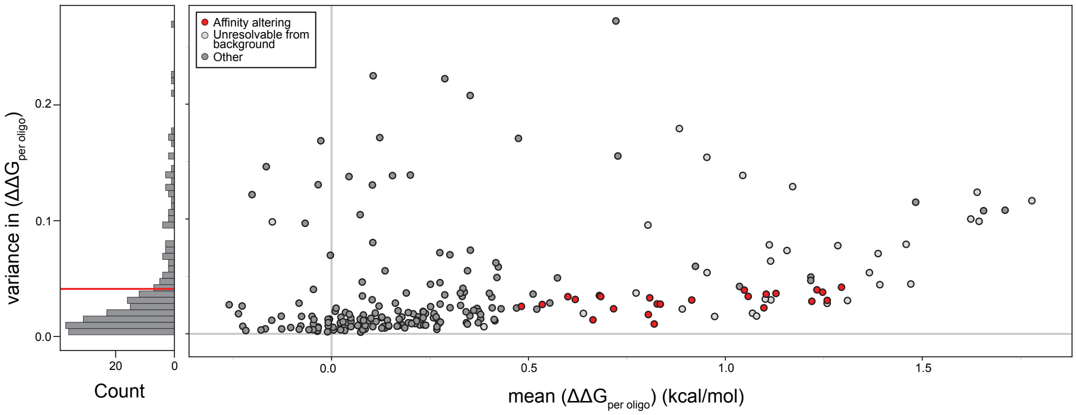

Figure S29: Classification of affinity-altering substitutions in MAX. **Left:** Histogram of variance in $\Delta\Delta G_{peroligo}$ between all measured oligonucleotides. Red line indicates upper bound for variance of affinity altering mutations (variance $\leq 0.04$), excluding mutations with the highest quartile of variance in measured $\Delta\Delta G$s. **Right:** Mean $\Delta\Delta G_{peroligo}$ versus variance in $\Delta\Delta G_{peroligo}$ for all MAX mutations. Red markers indicate putative "affinity altering" mutations. Light grey points indicate mutations un-resolvable from background in at least one oligonucleotide.

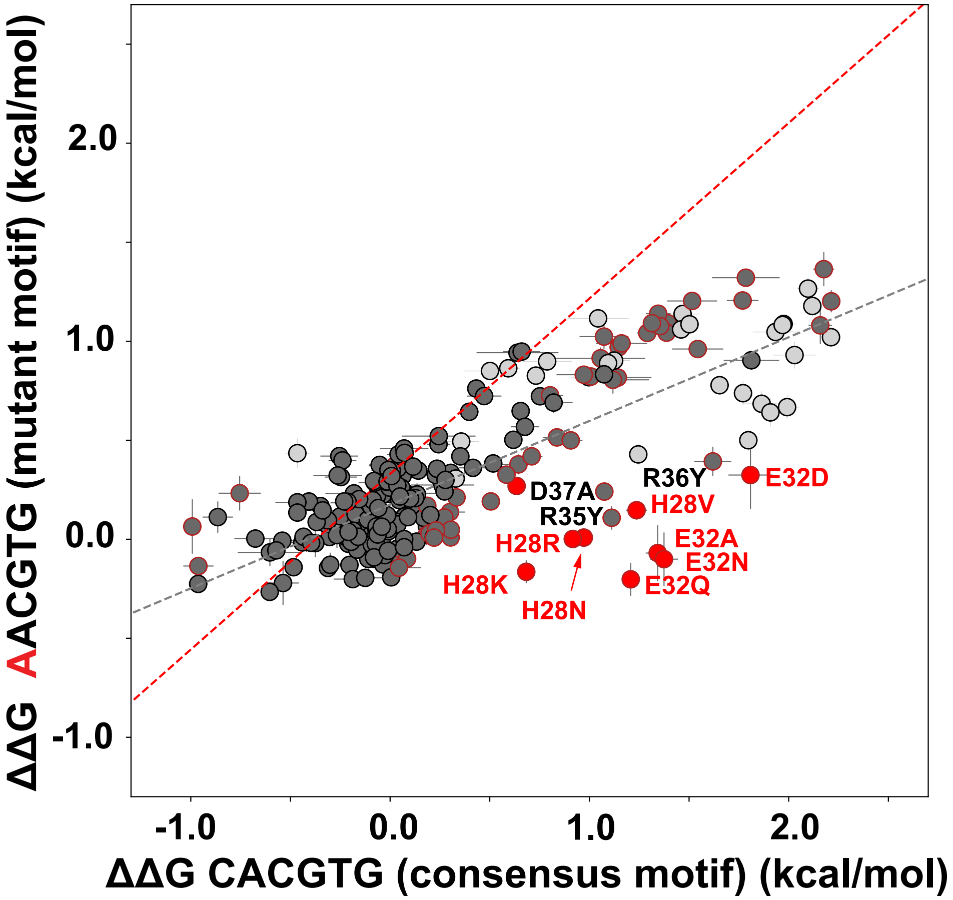

Figure S30: Pairwise comparison between measured $\Delta\Delta G$s for MAX mutants interacting with a low-affinity mutant sequence (5′-C **A**ACGTG A-3′) versus the reference sequence (5′-C CACGTG A-3′). Each marker indicates the median $\Delta\Delta G$ (± SEM) for a given TF mutant across all chambers in all replicates. Grey dashed line indicates linear regression $y = 0.42 * x + 0.17$; red dashed line indicates identity line. Markers corresponding to TF mutations to known crystallographic contacts to nucleotide bases (H28 and E32) are colored in red. Non-additive mutations in $K_d$ space are indicated by red outlines; see **Table S6**.

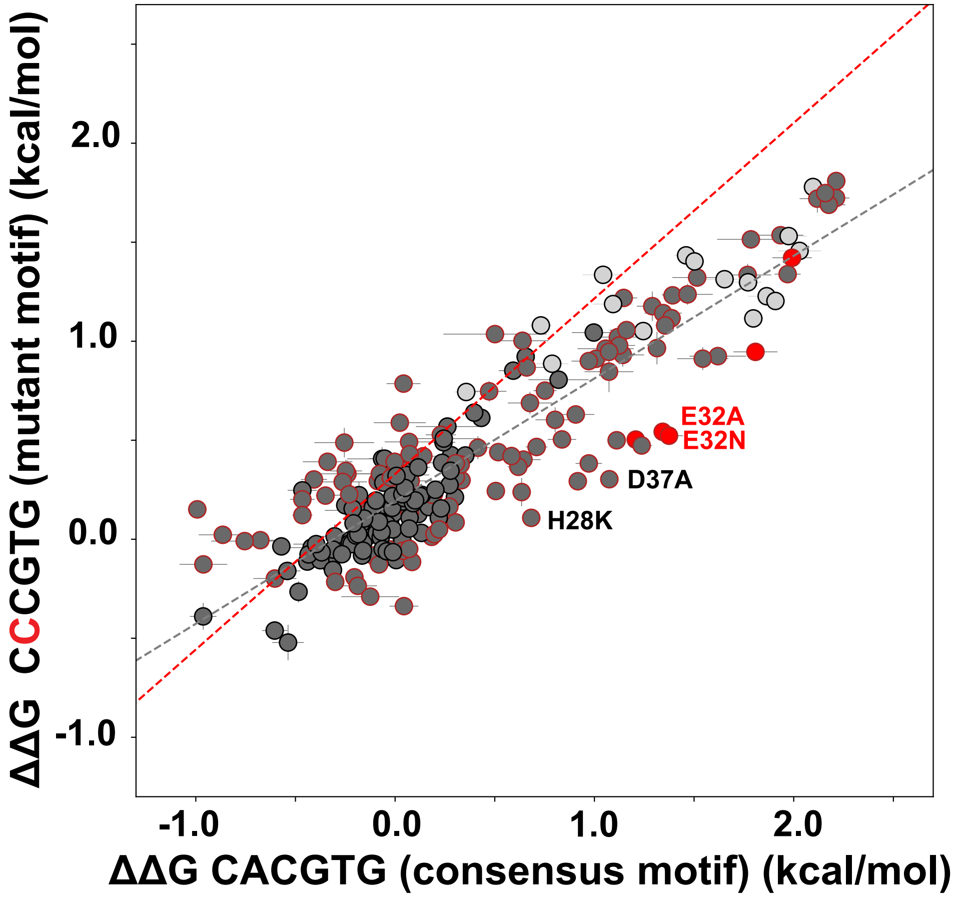

Figure S31: Pairwise comparison between measured binding affinities for MAX mutants interacting with a low-affinity mutant sequence (5′-C C**G**CGTG A-3′) versus the reference sequence (5′-C CACGTG A-3′). Each marker indicates the median $\Delta\Delta G$ (± SEM) for a given TF mutant across all chambers in all replicates. Grey dashed line indicates linear regression $y = 0.62 * x + 0.19$; red dashed line indicates identity line. Markers corresponding to TF mutations to known crystallographic contacts to nucleotide bases (H28 and E32) are colored in red. Non-additive mutations in $K_d$ space are indicated by red outlines; see **Table S6**.

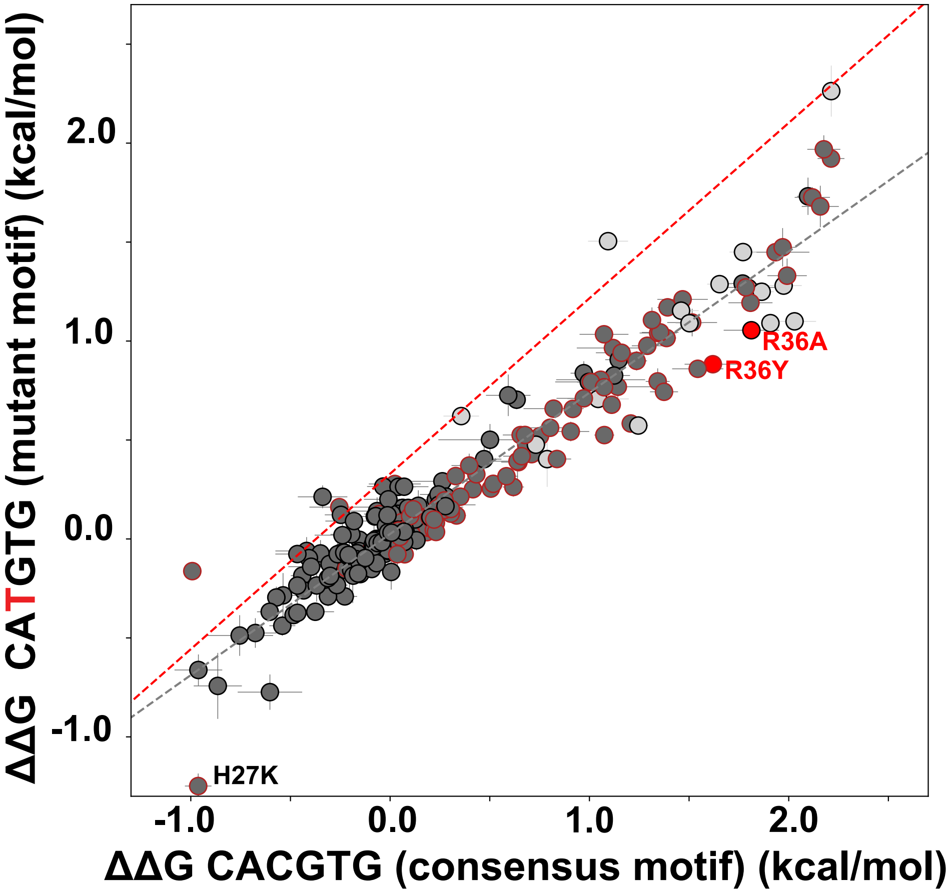

Figure S32: Pairwise comparison between measured binding affinities for MAX mutants interacting with a low-affinity mutant sequence (5′-C CA**T**GTG A-3′) versus the reference sequence (5′-C CACGTG A-3′). Each marker indicates the median $\Delta\Delta G$ (± SEM) for a given TF mutant across all chambers in all replicates. Grey dashed line indicates linear regression $y = 0.71 * x + 0.02$; red dashed line indicates identity line. Markers corresponding to TF mutations to known crystallographic contacts to nucleotide bases (R36) are colored in red. Non-additive mutations in $K_d$ space are indicated by red outlines; see **Table S6**.

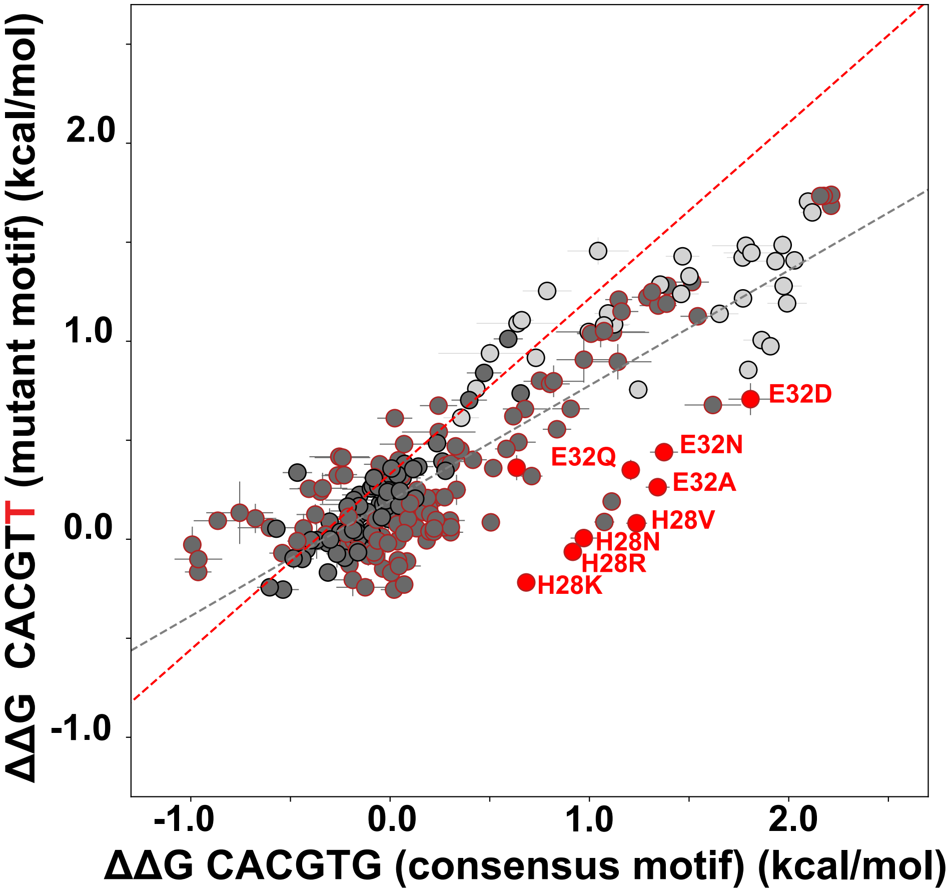

Figure S33: Pairwise comparison between measured binding affinities for MAX mutants interacting with a low-affinity mutant sequence (5′-C CACGT**T** A-3′) versus the reference sequence (5′-C CACGTG A-3′). Each marker indicates the median $\Delta\Delta G$ (± SEM) for a given TF mutant across all chambers in all replicates. Grey dashed line indicates linear regression $y = 0.58 * x + 0.19$; red dashed line indicates identity line. Markers corresponding to TF mutations to known crystallographic contacts to nucleotide bases (H28 and E32) are colored in red. Non-additive mutations in $K_d$ space are indicated by red outlines; see **Table S6**.

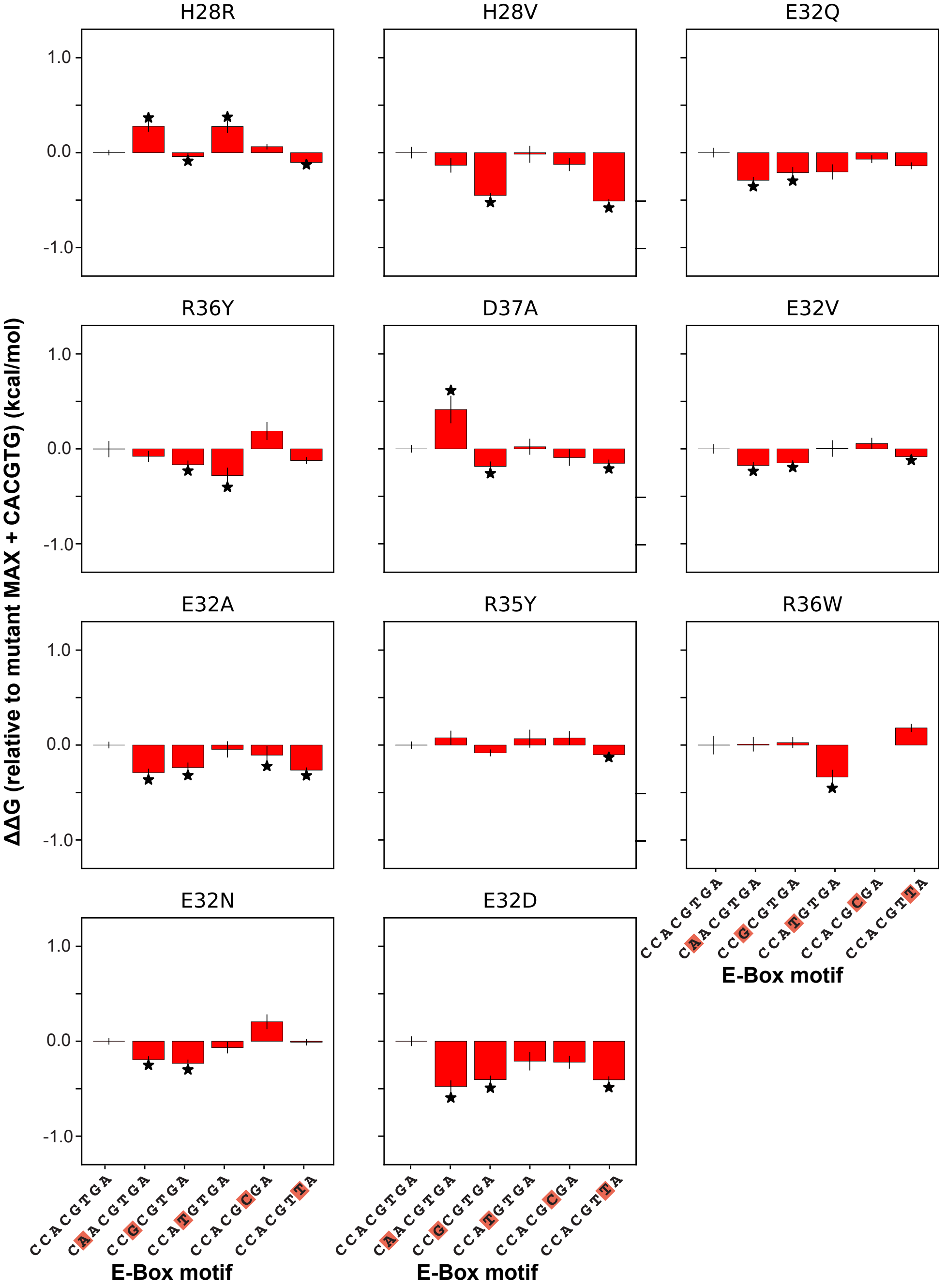

Figure S34: Mutations that alter sequence specificity such that a motif besides CACGTG is lowest in affinity. Bar plots indicate $\Delta\Delta G$ ($\pm$ SEM) for each mutation defined as $K_{d,mutant+mutantE-Box}$ relative to the $K_{d,mutant+CACGTG}$. Asterisks indicate significance (p-value $< 0.05$ in independent T-test comparing measured $K_d$s for the MAX mutant binding CACGTG versus the MAX mutant binding alternate motifs).

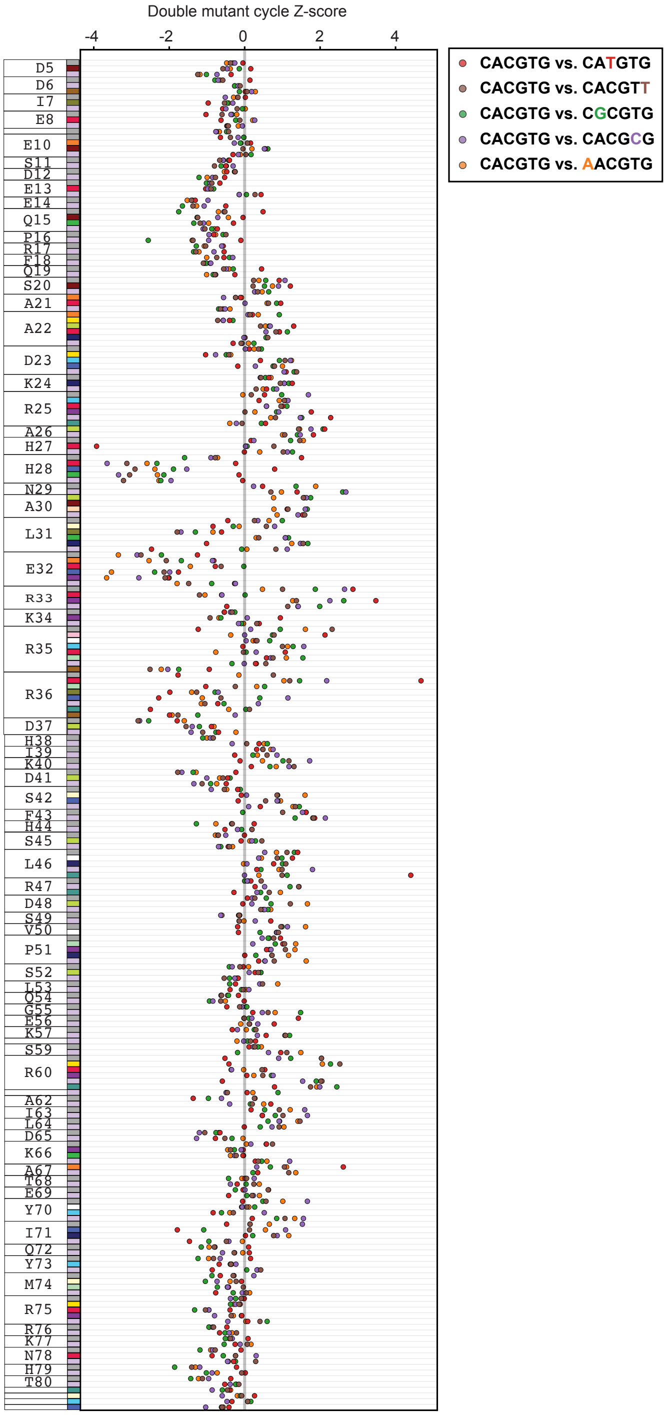

Figure S35: Residual Z-score for each consensus oligonucleotide versus mutant oligonucleotide double mutant cycle comparison as a function of MAX amino acid sequence. Each marker indicates the residual Z-score of a MAX mutation computed from a double mutant cycle comparing $\Delta\Delta G$ of a mutated E-box motif to the cognate motif. Marker colors indicate the identity of the mutated oligonucleotide in the double mutant cycle comparison.

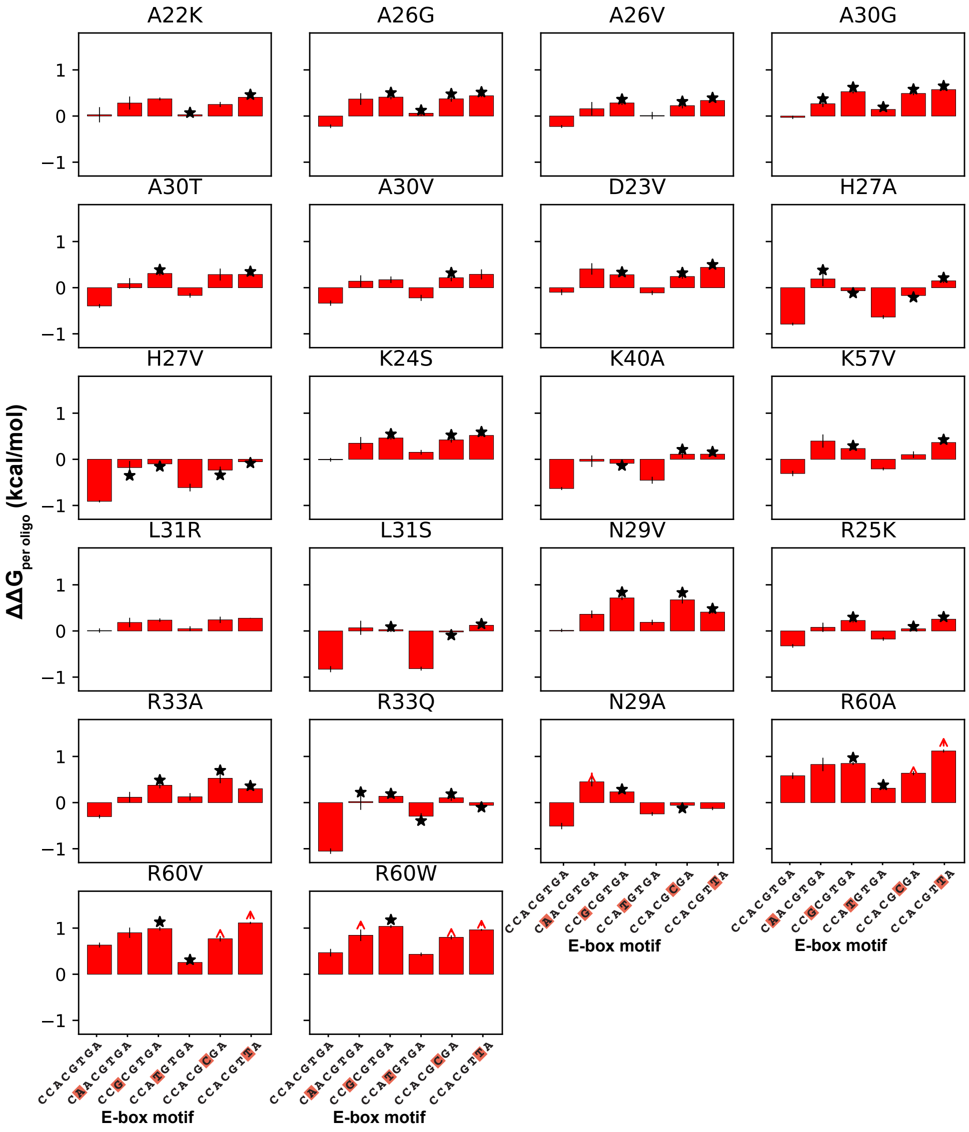

Figure S36: Bar plots indicate $\Delta\Delta G$ ($\pm$ SEM) for each selective mutation relative to WT MAX in different oligonucleotide binding interactions. Asterisks indicate non-additive binding relative to WT MAX and CACGTG. Binding interactions that are unresolvable from background are annotated with red arrows, indicating that the plotted $\Delta\Delta G$ represents a lower bound estimate of the true difference in binding energies.

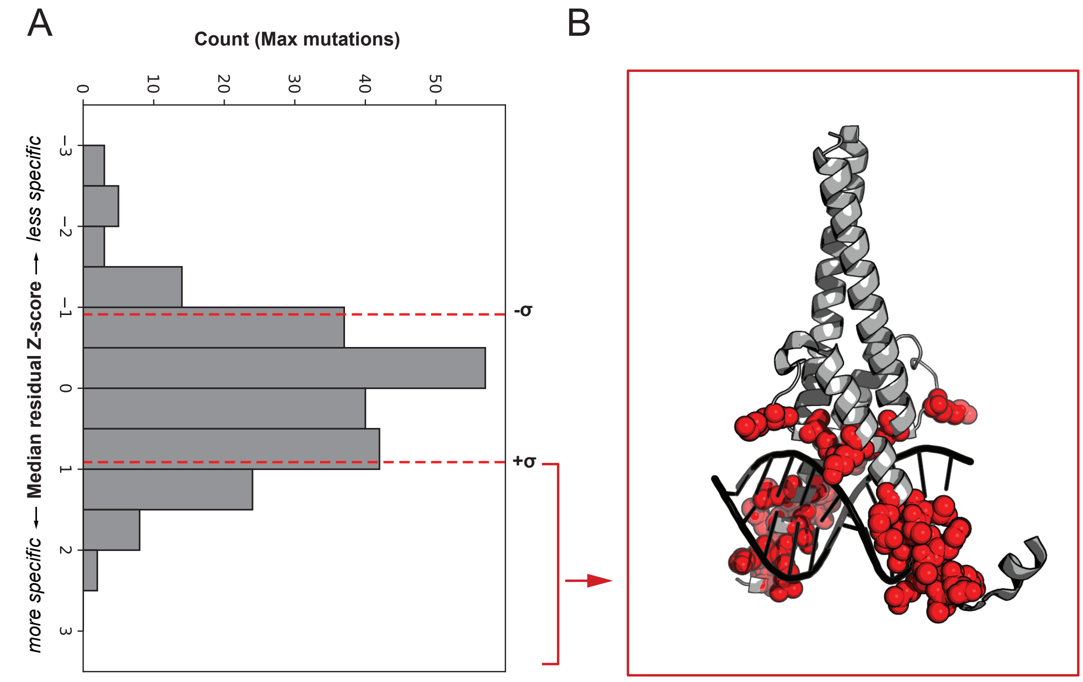

Figure S37: Definition and location within structure of specificity-increasing MAX mutations, listed in **Figure S36**. **A** Histogram of median residual Z-scores for all measured MAX mutations, with threshold for determining specificity altering mutations indicated by the dashed red line (|median residual Z-score| $> 0.9$). **B** Location of specificity increasing mutations (median residual Z-score $> 0.9$) indicated on the MAX structure (PDB ID: 1HLO) in red.

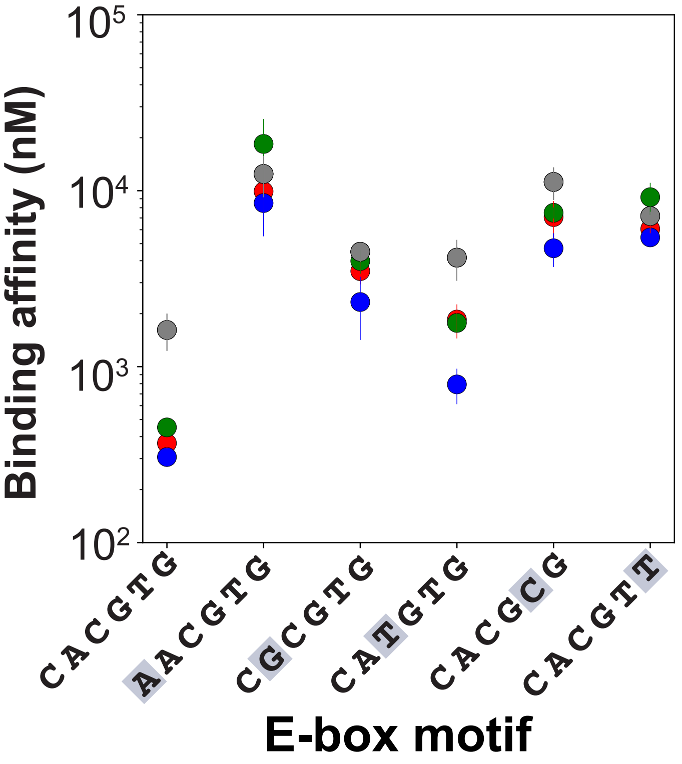

Figure S38: Median $K_d$ ± SEM across all E-box motifs for WT and all measured substitutions at H27.

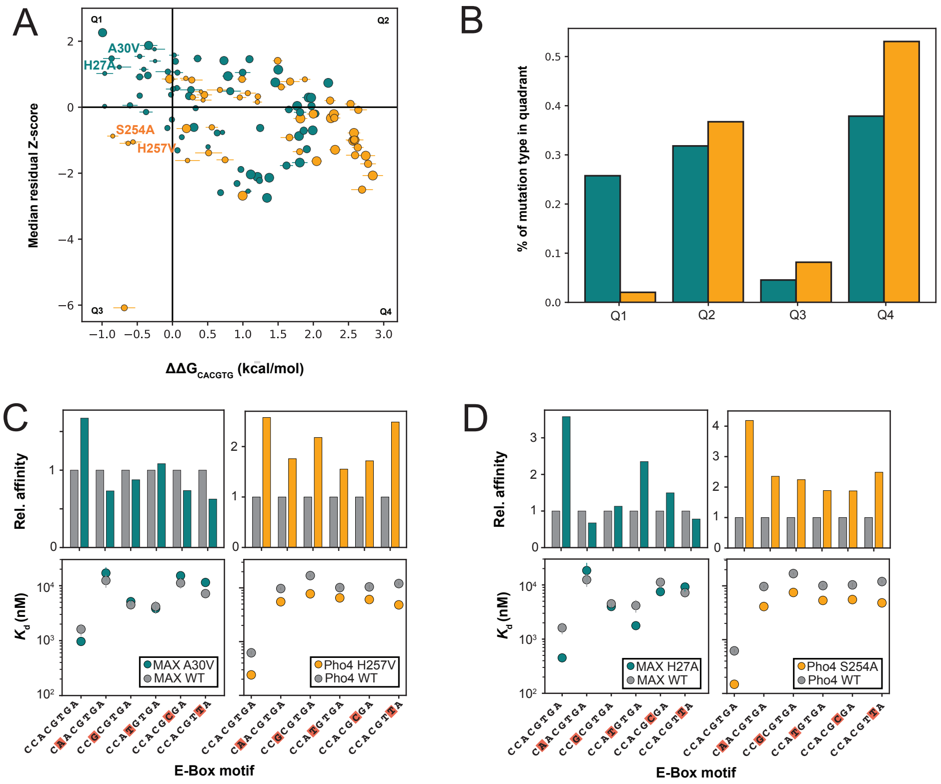

Figure S39: **(A)** Median residual Z-score ("specificity score") versus $\Delta\Delta G_{CACGTG}$ for MAX (teal) and Pho4 (orange). Each point represents one TF mutation within the basic region. Size of marker indicates the relative conservation of the mutated residue. **(B)** Quantification of distribution of mutations in each quadrant for both MAX (teal) and Pho4 (orange). **(C)** Relative affinity (to WT) and measured $K_d$s for comparable MAX and Pho4 mutations. Markers indicate $K_d$s (median ± SEM) for WT TF (grey), MAX A30V (teal) and Pho4 H257V (orange). **(D)** Relative affinity (to WT) and measured $K_d$s for comparable MAX and Pho4 mutations. Markers indicate $K_d$s (median ± SEM) for WT TF (grey), MAX H27A (teal) and Pho4 S254A (orange).

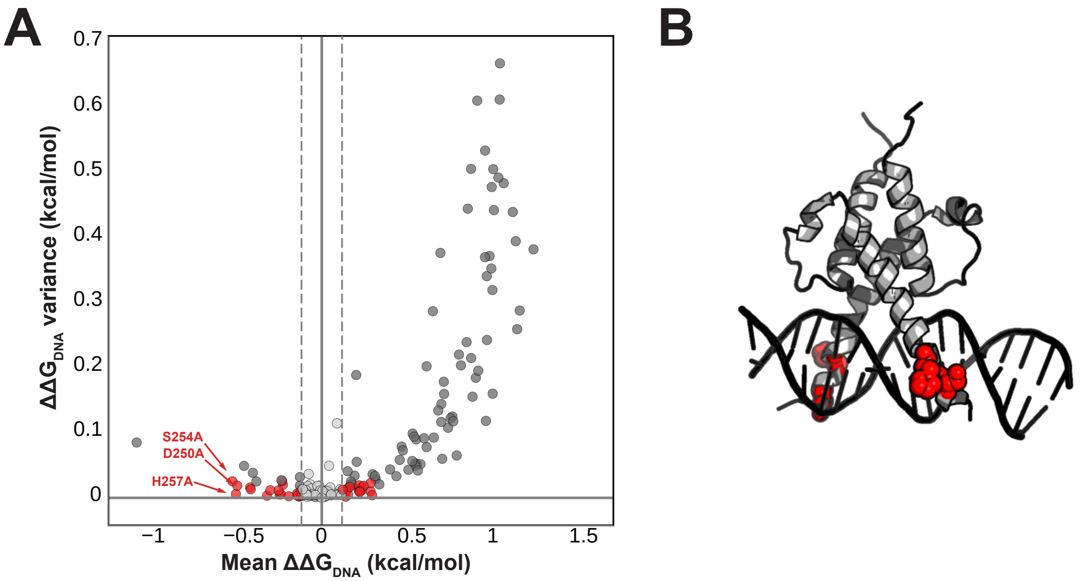

Figure S40: **(A)** Mean energetic impact of a TF mutation on DNA binding across all measured DNA sequences ($\langle \Delta\Delta G_{DNA} \rangle$) versus variance in the energetic impact of a TF mutation on DNA binding across all measured DNA sequences (variance of $\Delta\Delta G_{DNA}$). Each point represents a unique Pho4 mutation. Dotted lines indicate 2*SEM difference from WT $\langle \Delta\Delta G_{DNA} \rangle$ for all oligonucleotides. Red points are mutations that can be confidently called as affinity-altering mutations within the measurement variance, defined as mutations where(variance of $\Delta\Delta G_{DNA}$) is less than $\langle \Delta\Delta G_{DNA} \rangle$. **(B)** Mutations where the absolute value of $\langle \Delta\Delta G_{DNA} \rangle > 0.5$ kcal/mol, colored in red on the Pho4 crystal structure (PDB ID: 1A0A).

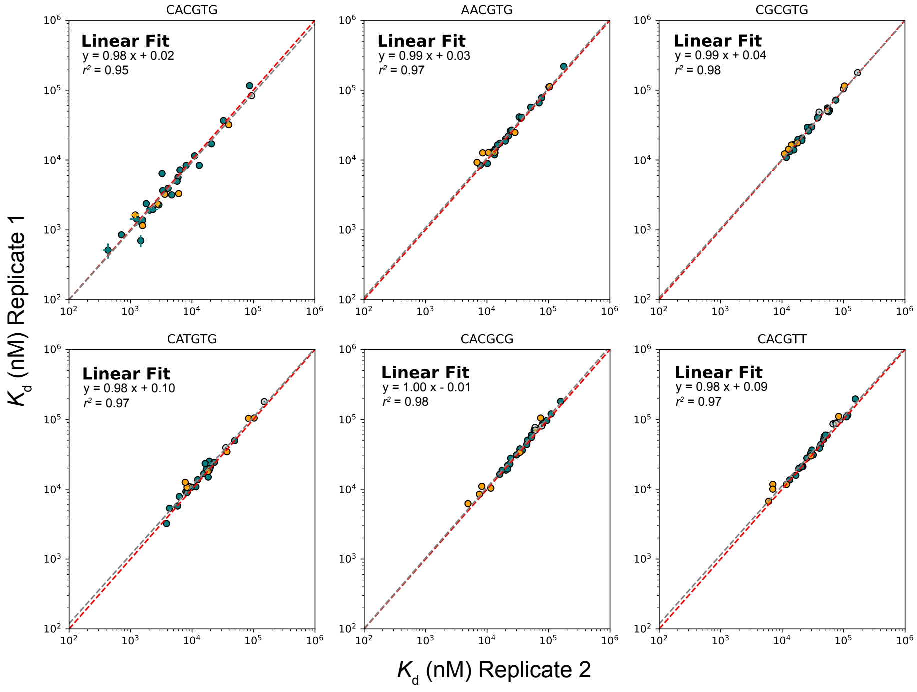

Figure S41: Pairwise comparisons of per-mutant $K_d$s for a small library of MAX (teal) and Pho4 (orange) mutants across 2 experiments. Points indicate median affinities (± SEM) for each TF mutant. Linear fits are indicated by grey dashed lines; identity lines are indicated by red dashed lines. Light grey points indicate mutant $K_d$ measurements that are not statistically significantly different from background binding in all replicates, listed in Table S3.

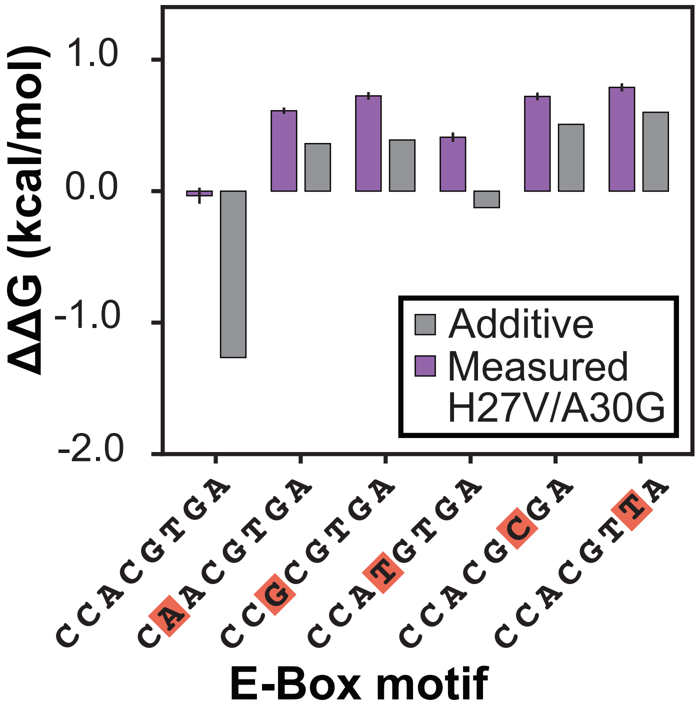

Figure S42: Expected additive $\Delta\Delta G$ (calculated as summed $\Delta\Delta G$ of each mutation alone) versus measured $\Delta\Delta G$ of MAX H27V/A30G double mutant across all E-box motifs.

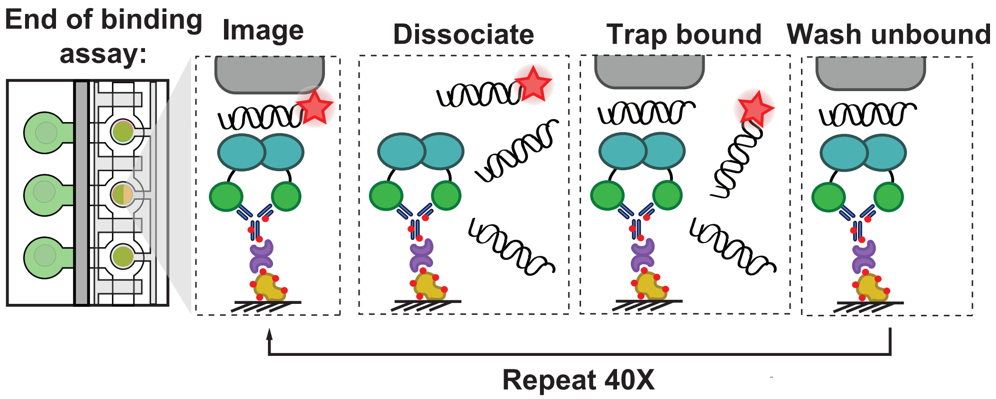

Figure S43: Experimental workflow illustrating iterative trapping and dissociation of fluorescently-labelled DNA to quantify binding off-rates with imaging steps indicated. The addition of high-affinity unlabeled competitor DNA during dissociation prevents rebinding.

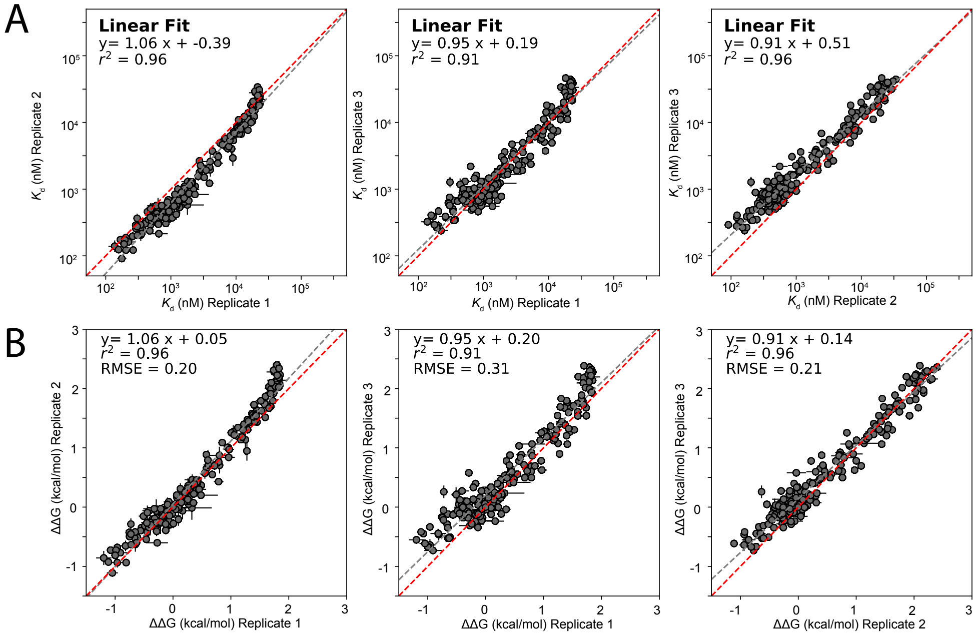

Figure S44: Pairwise comparisons of per-mutant **A** $K_d$s and **B** apparent $\Delta\Delta G$s for Pho4 mutants across 3 experiments. Markers indicate values (median ± SEM) for each mutant. Linear fits are indicated by grey dashed lines; identity lines are indicated by red dashed lines.

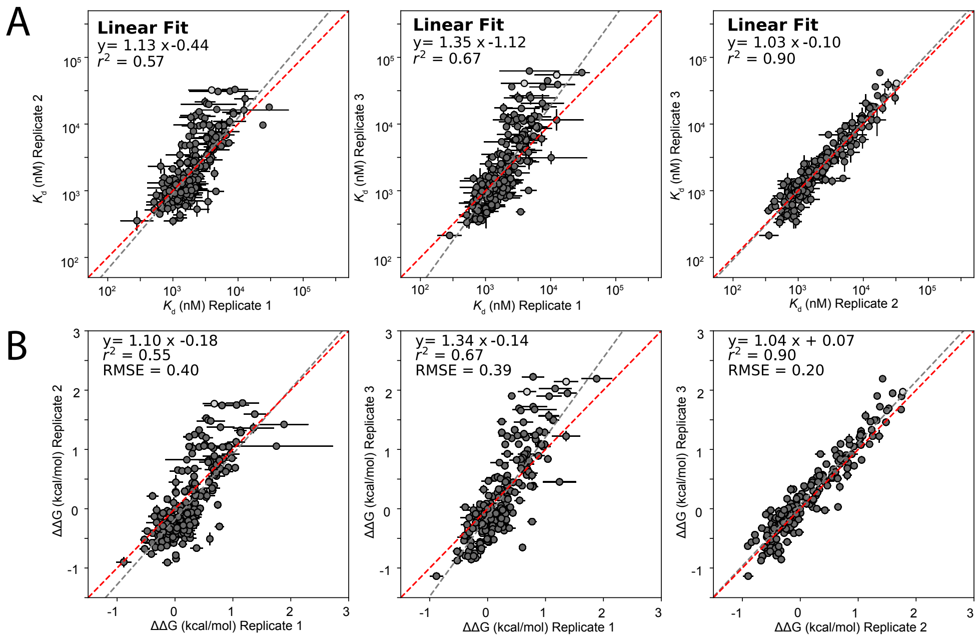

Figure S45: Pairwise comparisons of per-mutant **A** $K_d$s and **B** apparent $\Delta\Delta G$s for MAX mutants across 3 experiments. Markers indicate rates (median ± SEM) for each mutant. Linear fits are indicated by grey dashed lines; identity lines are indicated by red dashed lines. Light grey points indicate mutant $K_d$ measurements that are not statistically significantly different from background binding in all replicates, listed in Table S3.

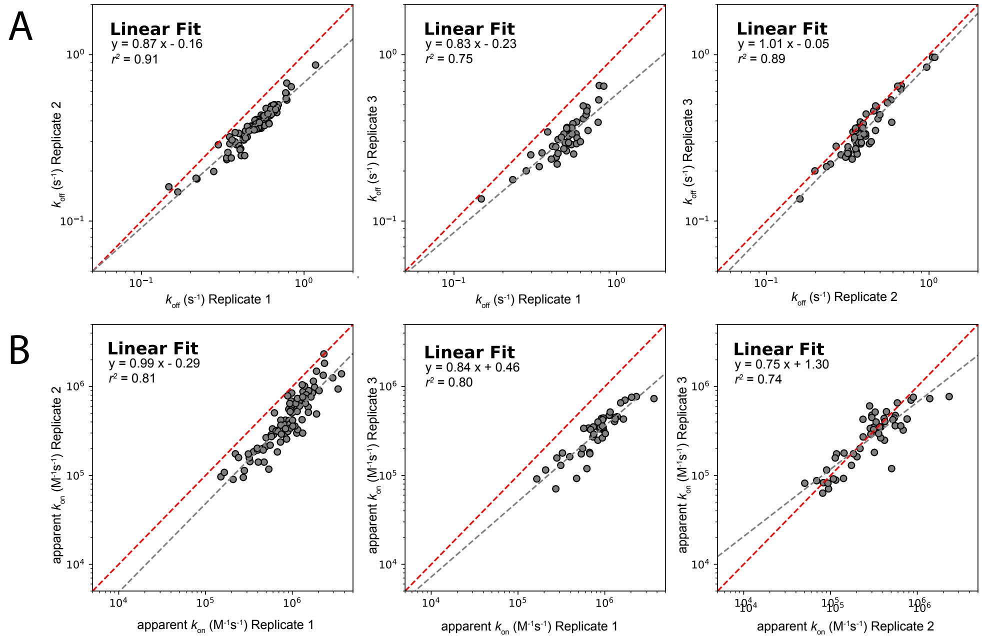

Figure S46: Pairwise comparisons of per-mutant **A** $k_{off}$s and **B** apparent $k_{on}$s for Pho4 mutants across 3 experiments. Markers indicate rates (median ± SEM) for each mutant. Linear fits are indicated by grey dashed lines; identity lines are indicated by red dashed lines.

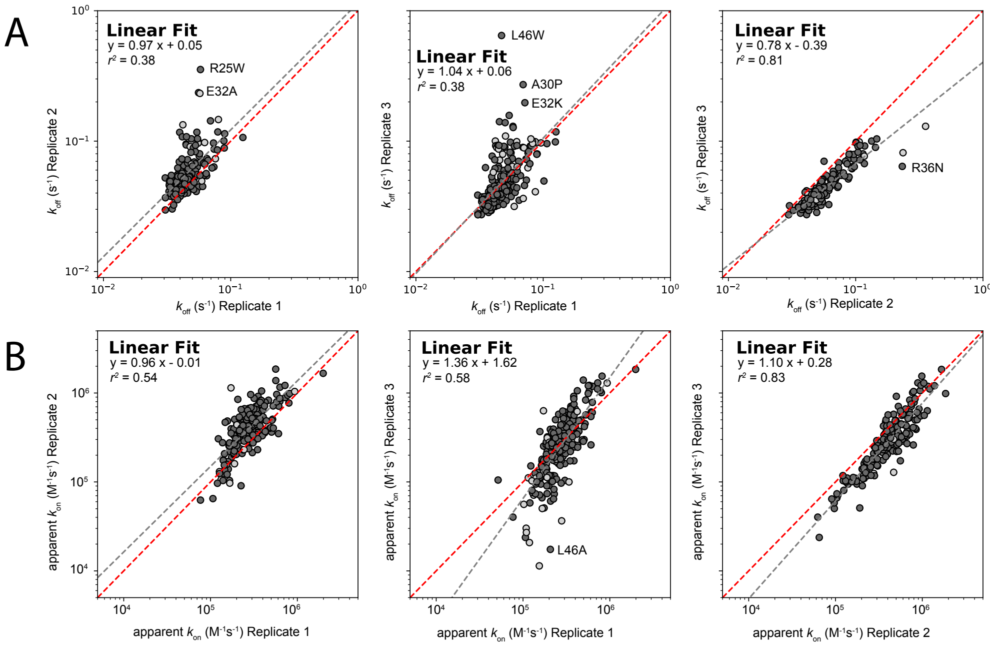

Figure S47: Pairwise comparisons of per-mutant **A** $k_{off}$s and **B** apparent $k_{on}$s for MAX mutants across 3 experiments. Markers indicate rates (median ± SEM) for each mutant. Linear fits are indicated by grey dashed lines; identity lines are indicated by red dashed lines. Light grey points indicate mutant $K_d$ measurements that are not statistically significantly different from background binding in all replicates, listed in Table S3. Identities of select outlier mutations are annotated.

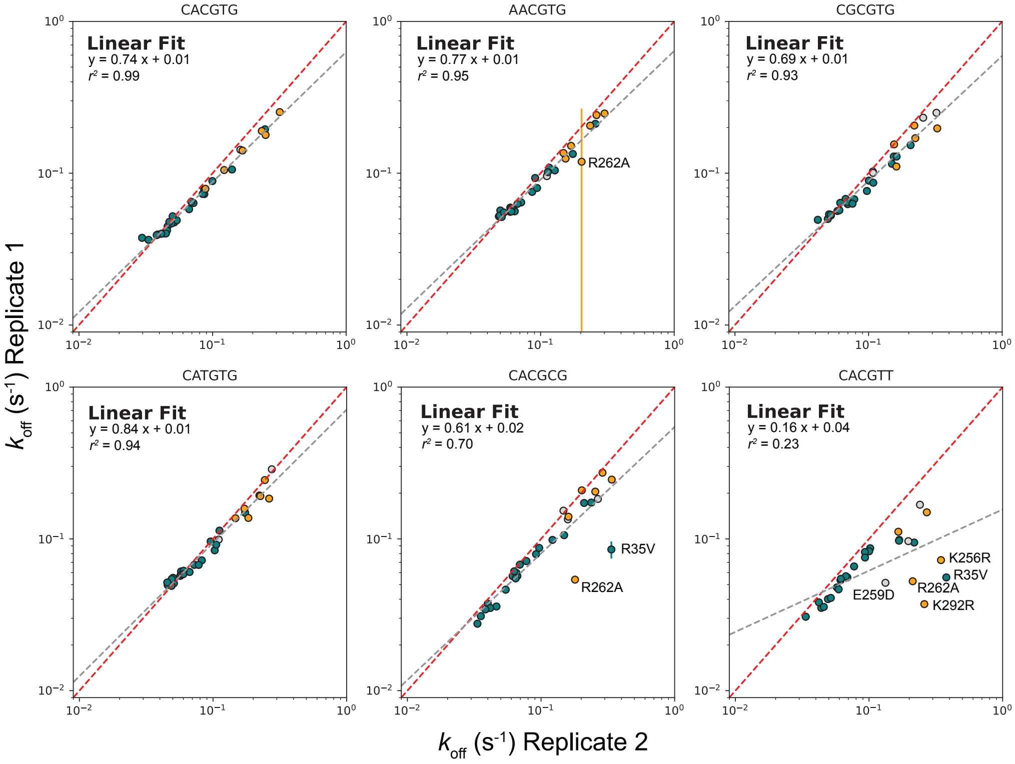

Figure S48: Pairwise comparisons of per-mutant $k_{off}$s for a small library of MAX (teal) and Pho4 (orange) mutants across 2 experiments. Markers indicate off-rates (median ± SEM) for each TF mutant. Linear fits are indicated by grey dashed lines; identity lines are indicated by red dashed lines. Light grey points indicate mutant $K_d$ measurements that are not statistically significantly different from background binding in all replicates, listed in Table S3.

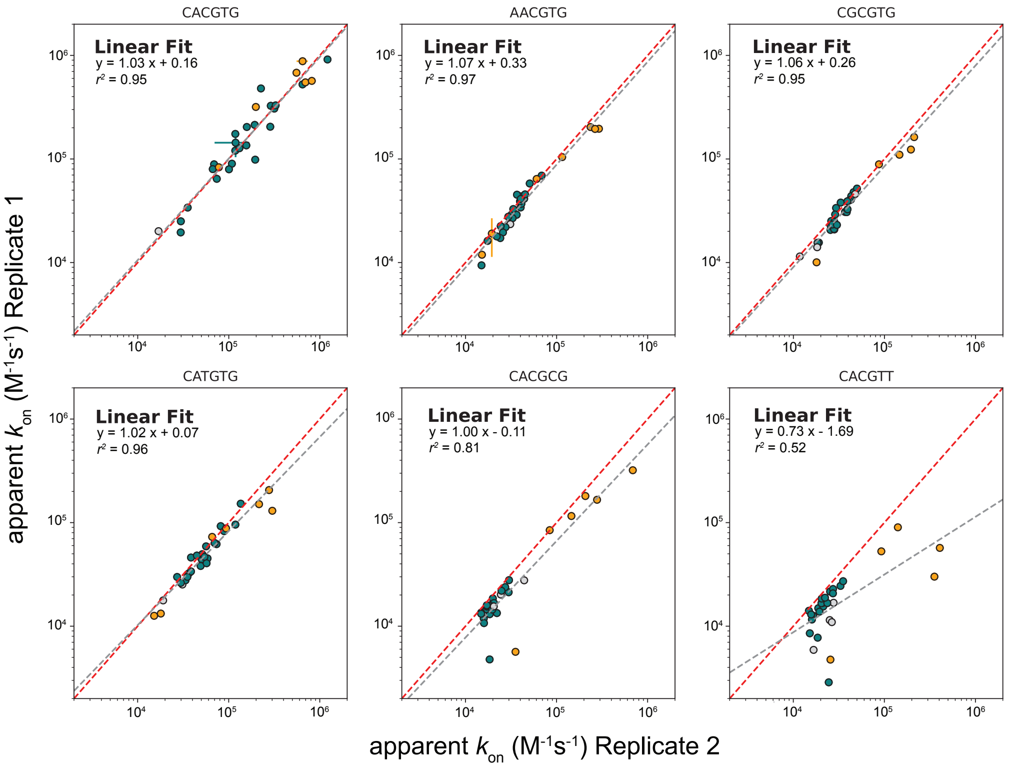

Figure S49: Pairwise comparisons of per-mutant apparent $k_{on}$s for a small library of MAX (teal) and Pho4 (orange) mutants across 2 experiments. Markers indicate on-rates (median $\pm$ SEM) for each TF mutant. Linear fits are indicated by grey dashed lines; identity lines are indicated by red dashed lines. Light grey points indicate mutant $K_d$ measurements that are not statistically significantly different from background binding in all replicates, listed in Table S3.

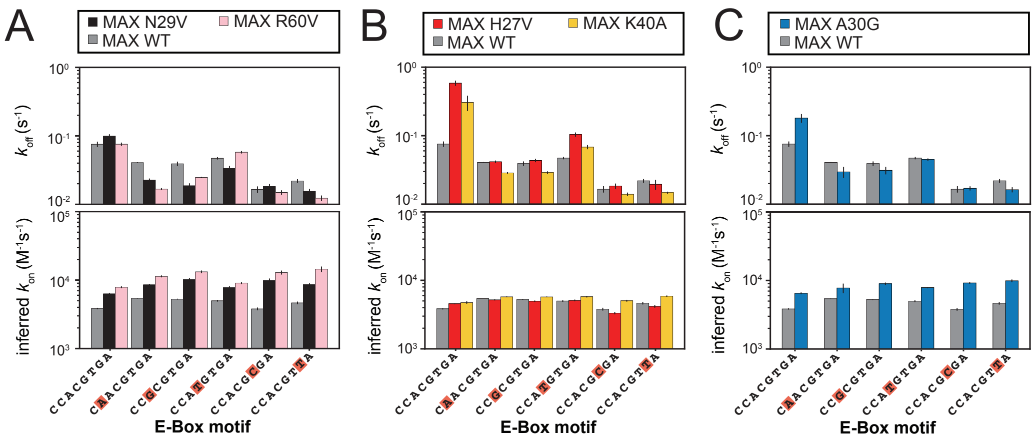

Figure S50: Values (median ± SEM) of measured off- (top) and on-rate (bottom) constants for selective MAX mutants interacting with many E-box variants. Mutations shown in **(A)** speed up off-rate for all DNA sequences, mutations in **(B)** speed up on-rate mostly for the cognate motif, and the MAX mutation in **(C)** exhibits both behaviors.

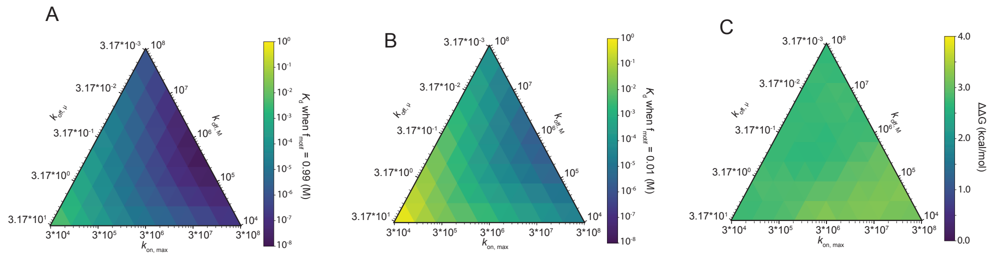

Figure S51: Ternary heat maps demonstrating that the 3-state model with one binding conformation shown in **6D** cannot explain differences in intrinsic selectivity for the same set of sequences. Simulated binding affinities for the most preferred sequence (defined as $f_{motif} = 0.99$) **(A)** and least preferred sequence (defined as $f_{motif} = 0.01$) **(B)** define binding selectivity **(C)**), the free energy difference between binding. All simulated affinities and free energies are shown as a function of microscopic rate constants $k_{off,\mu}$, $k_{off,M}$, and $k_{on,max}$. Each value represents the median across 3 simulation trajectories.

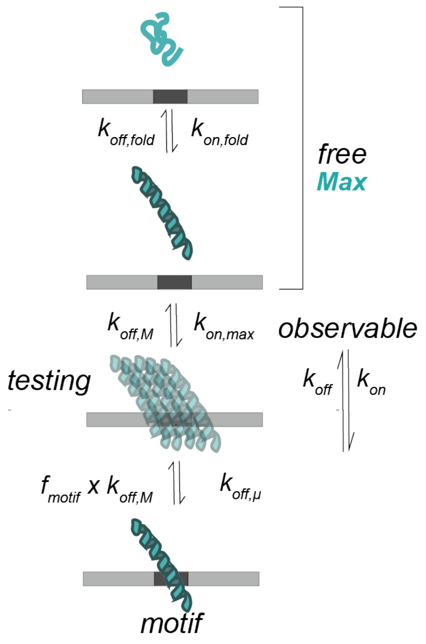

Figure S52: Example schematic showing 4-state model and associated rate constants describing folding-and-binding with a single binding-competent folded conformation.

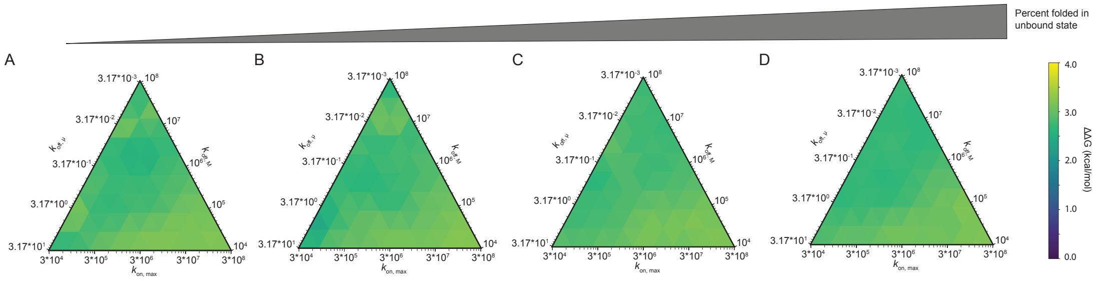

Figure S53: Ternary heat maps demonstrating that the 4-state model with one binding conformation shown in **S52** cannot explain differences in intrinsic selectivity for the same set of sequences. Each ternary heat map shows selectivity (the energetic difference between the most ($f_{motif} = 0.99$) and least ($f_{motif} = 0.01$) preferred sequences) as a function of microscopic rate constants $k_{off,\mu}$, $k_{off,M}$, and $k_{on,max}$, varying the degree of pre-folded structure in the unbound state (shown in order of least to most residual structure, **(A)** $\approx$ 25, **(B)** $\approx$ 50, **(C)** $\approx$ 75, and **(D)** $\approx$ 99 percent folded) between each plot. Each value represents the median across 3 simulation trajectories.

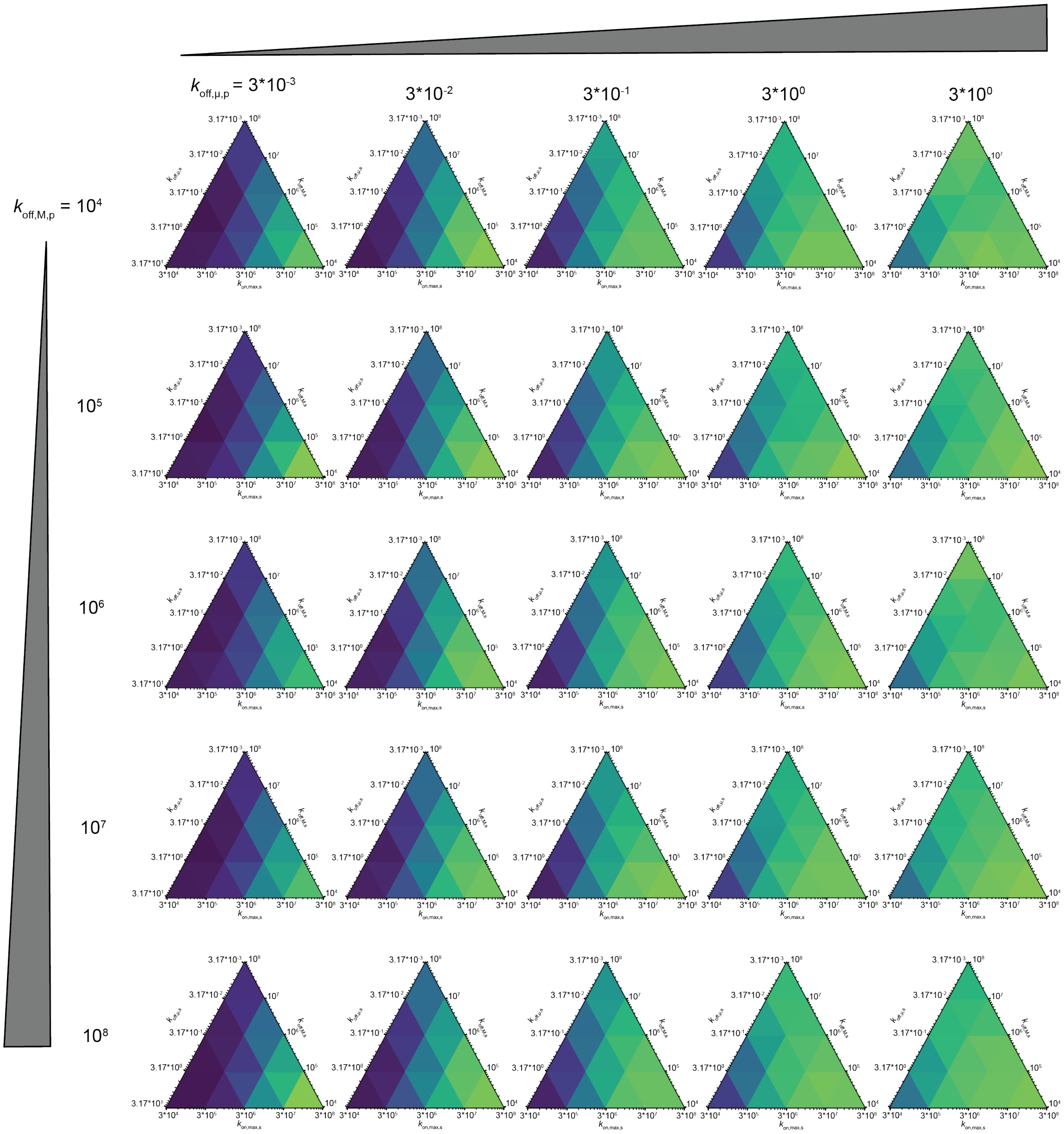

Figure S54: Ternary heat maps establishing that the 5-state model with distinct conformations model shown in **6F** can explain differences in intrinsic selectivity for the same set of sequences. Each ternary heat map shows selectivity (the energetic difference between the most ($f_{motif} = 0.99$) and least ($f_{motif} = 0.01$) preferred sequences) as a function of microscopic rate constants $k_{off,\mu,s}$, $k_{off,M,s}$, $k_{on,max,s}$, $k_{off,\mu,p}$, and $k_{off,M,p}$. Values on all ternary heat maps represent medians across 3 simulation trajectories when $k_{on,max,p} = 3 * 10^3$.

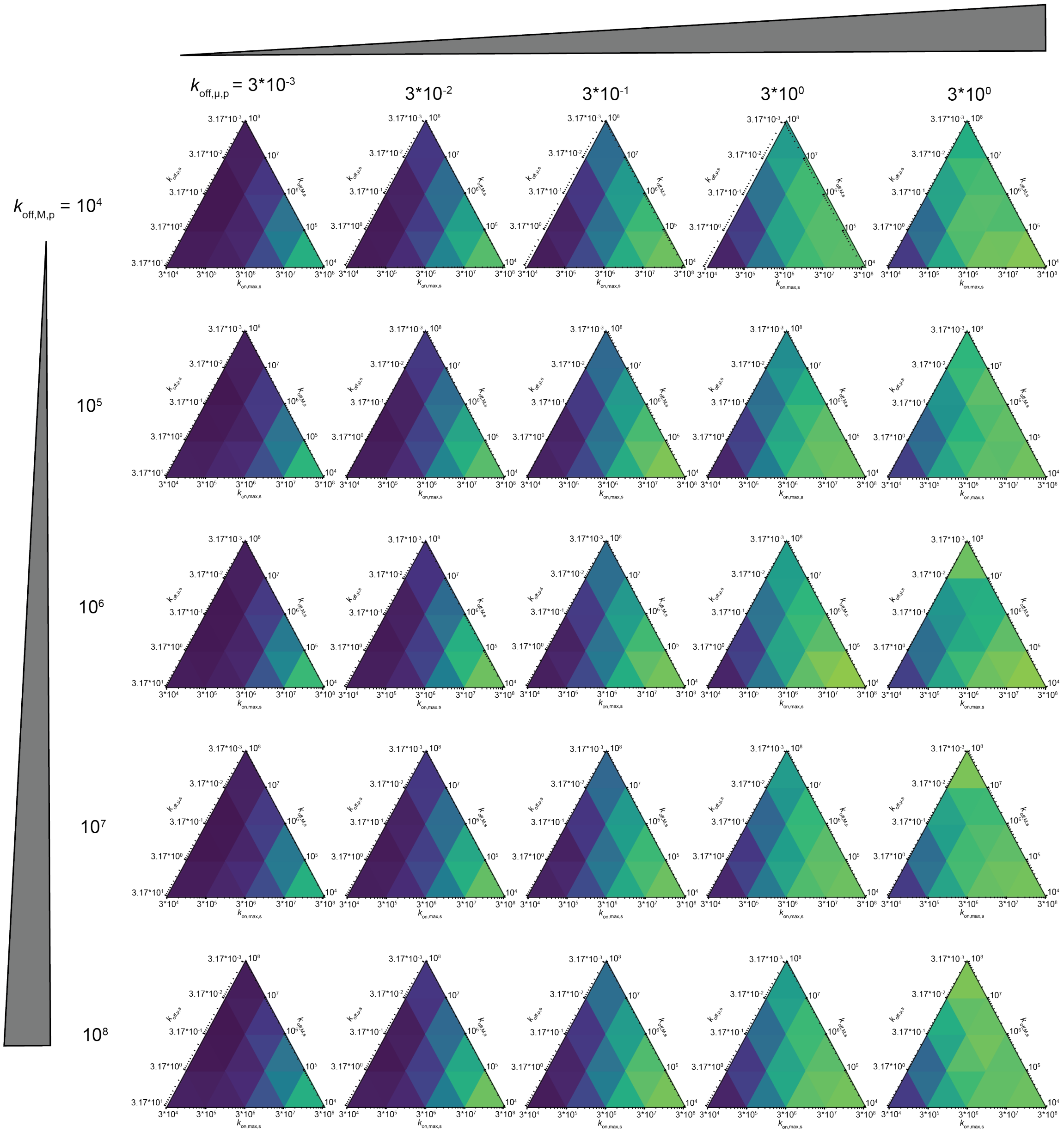

Figure S55: Ternary heat maps establishing that the 5-state model with distinct conformations model shown in **6F** can explain differences in intrinsic selectivity for the same set of sequences. Each ternary heat map shows selectivity (the energetic difference between the most ($f_{motif} = 0.99$) and least ($f_{motif} = 0.01$) preferred sequences) as a function of microscopic rate constants $k_{off,\mu,s}$, $k_{off,M,s}$, $k_{on,max,s}$, $k_{off,\mu,p}$, and $k_{off,M,p}$. Values on all ternary heat maps represent medians across 3 simulation trajectories when $k_{on,max,p} = 3 * 10^4$.

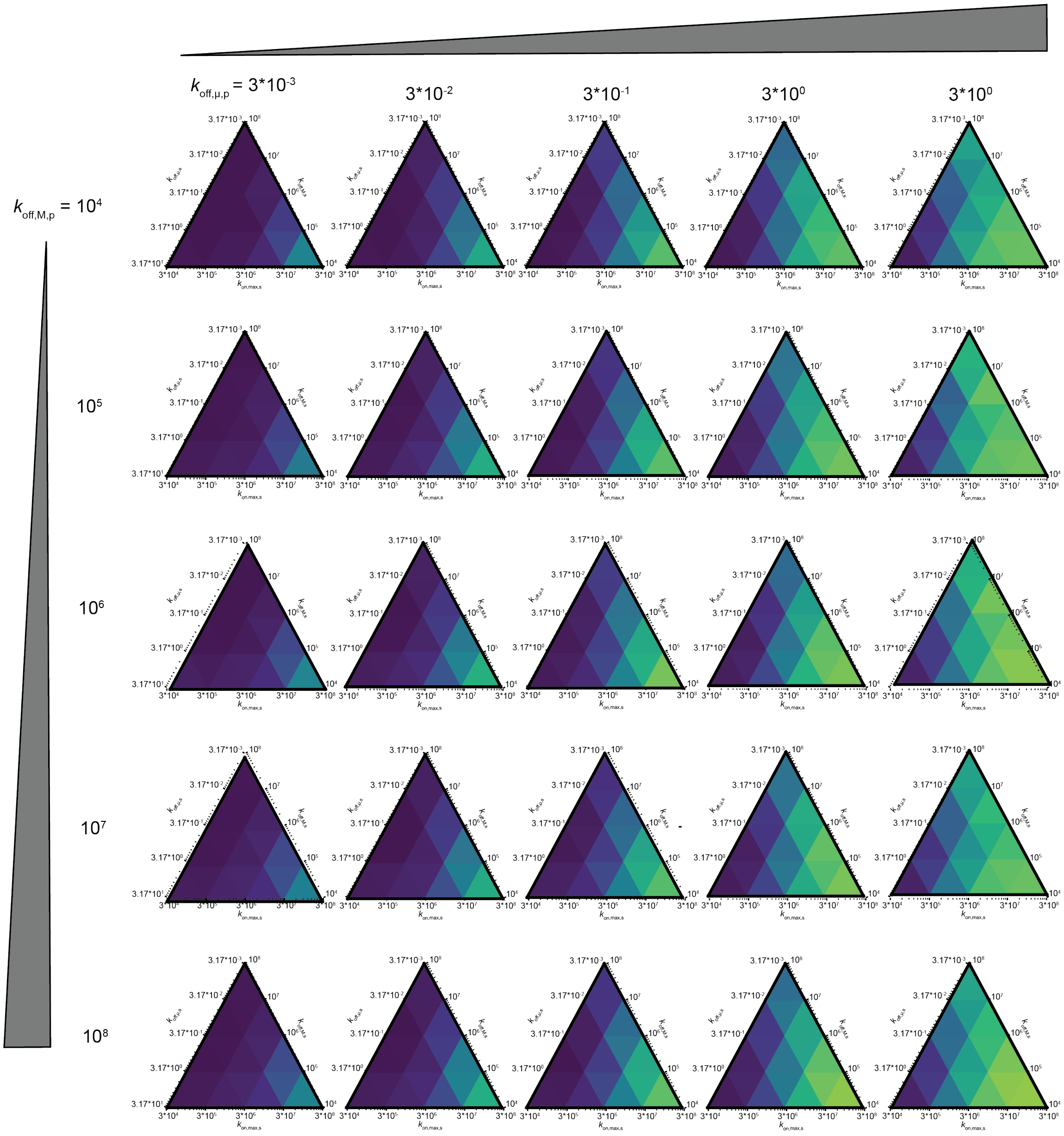

Figure S56: Ternary heat maps establishing that the 5-state model with distinct conformations model shown in **6F** can explain differences in intrinsic selectivity for the same set of sequences. Each ternary heat map shows selectivity (the energetic difference between the most ($f_{motif} = 0.99$) and least ($f_{motif} = 0.01$) preferred sequences) as a function of microscopic rate constants $k_{off,\mu,s}$, $k_{off,M,s}$, $k_{on,max,s}$, $k_{off,\mu,p}$, and $k_{off,M,p}$. Values on all ternary heat maps represent medians across 3 simulation trajectories when $k_{on,max,p} = 3 * 10^5$.

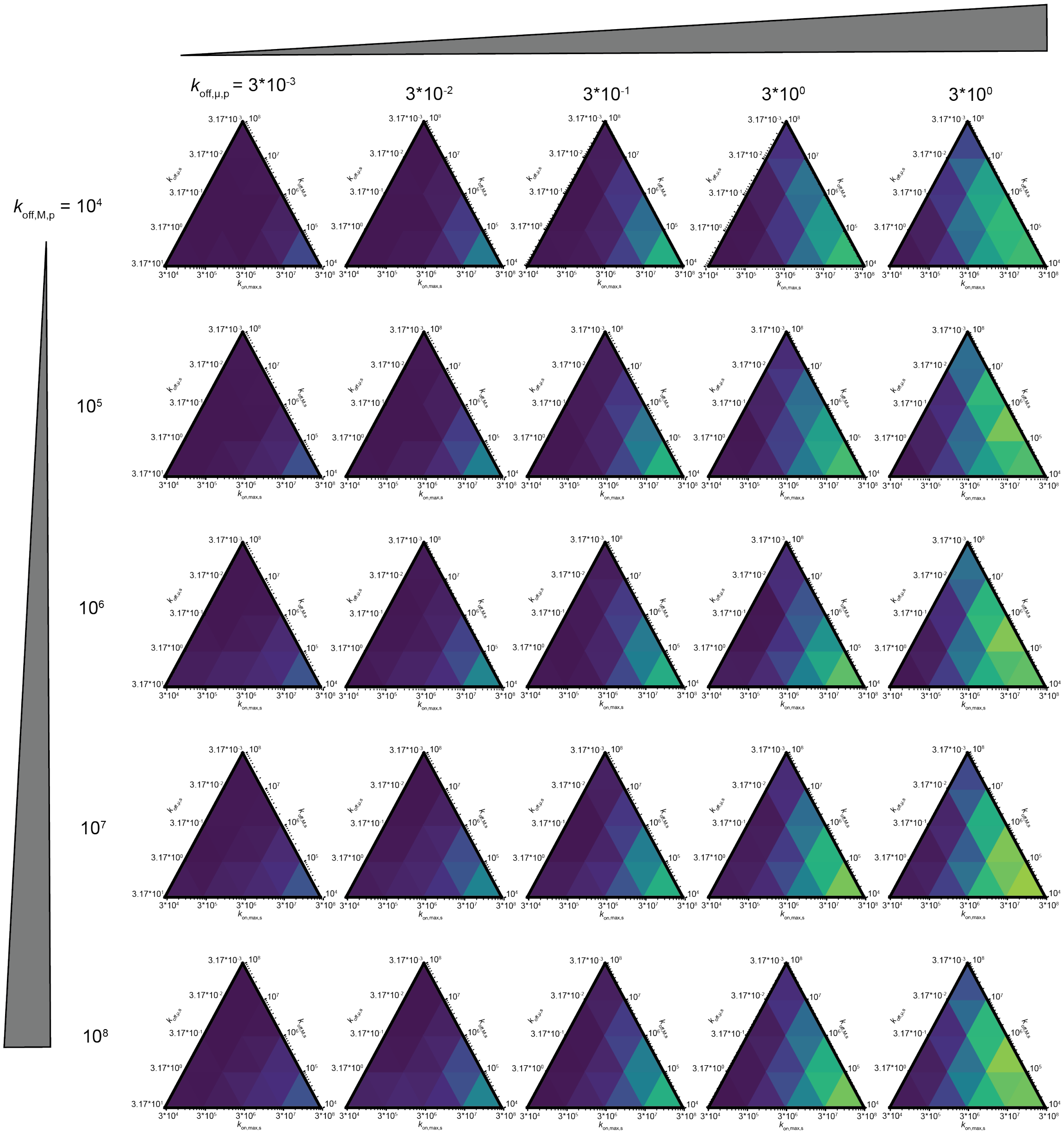

Figure S57: Ternary heat maps establishing that the 5-state model with distinct conformations model shown in **6F** can explain differences in intrinsic selectivity for the same set of sequences. Each ternary heat map shows selectivity (the energetic difference between the most ($f_{motif} = 0.99$) and least ($f_{motif} = 0.01$) preferred sequences) as a function of microscopic rate constants $k_{off,\mu,s}$, $k_{off,M,s}$, $k_{on,max,s}$, $k_{off,\mu,p}$, and $k_{off,M,p}$. Values on all ternary heat maps represent medians across 3 simulation trajectories when $k_{on,max,p} = 3 * 10^6$.

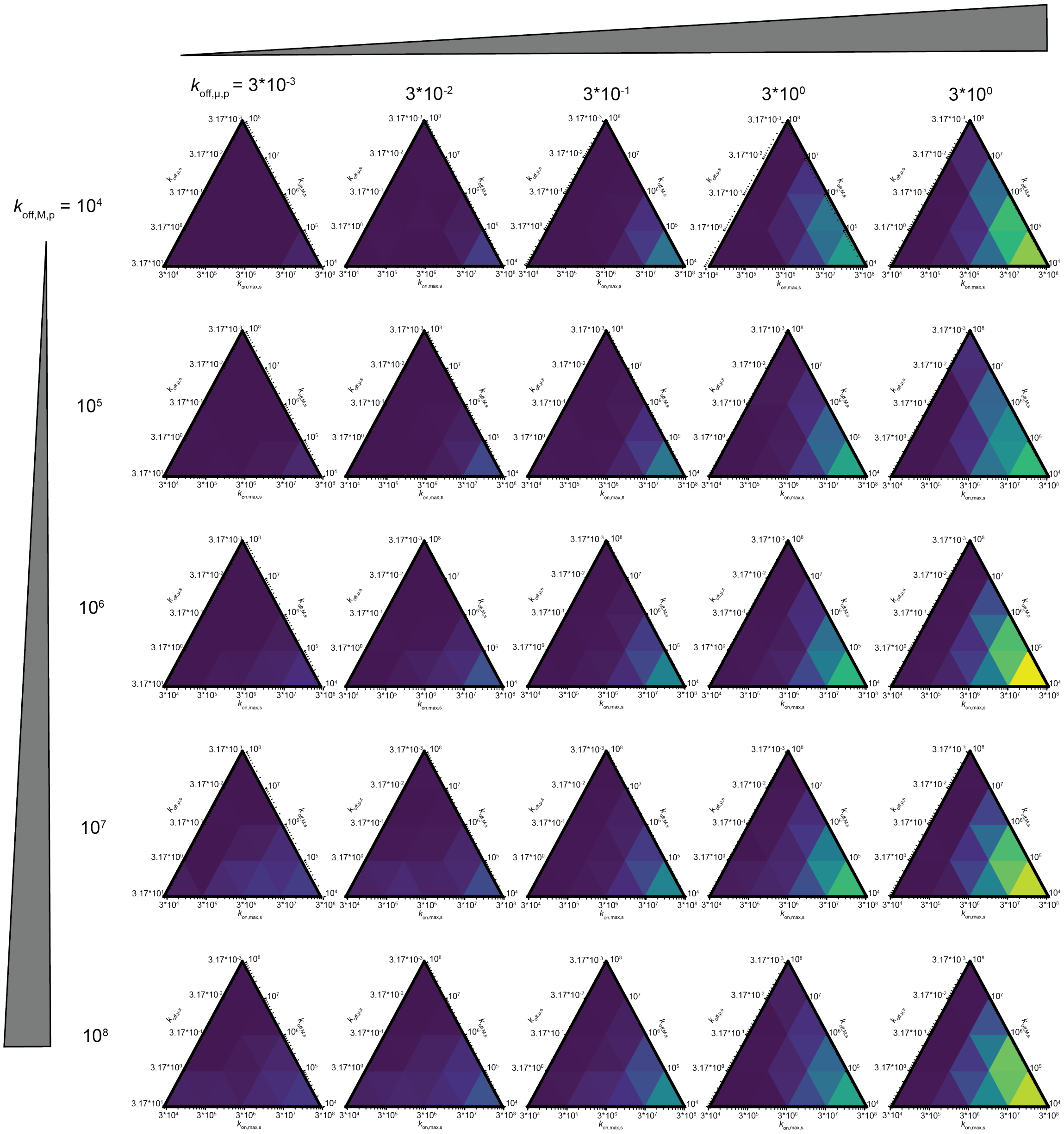

Figure S58: Ternary heat maps establishing that the 5-state model with distinct conformations model shown in **6F** can explain differences in intrinsic selectivity for the same set of sequences. Each ternary heat map shows selectivity (the energetic difference between the most ($f_{motif} = 0.99$) and least ($f_{motif} = 0.01$) preferred sequences) as a function of microscopic rate constants $k_{off,\mu,s}$, $k_{off,M,s}$, $k_{on,max,s}$, $k_{off,\mu,p}$, and $k_{off,M,p}$. Values on all ternary heat maps represent medians across 3 simulation trajectories when $k_{on,max,p} = 3 * 10^7$.

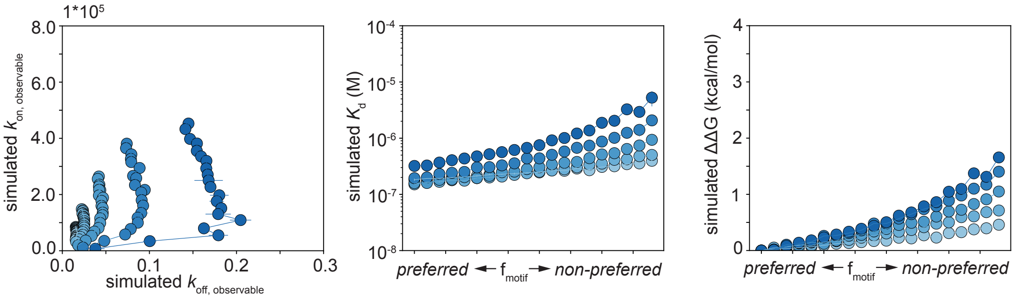

Figure S59: Co-varying $k_{on,max,specific}$ and $k_{off,\mu,specific}$ can recapitulate measured MAX A30G data in both kinetic (**A**), affinity (**B**), and free energy (**C**) space.

### 2 Supplemental tables

#### List of Tables

S1 Classification of non-alanine and valine scanning variants measured in this study. . . 69
S2 MAX mutants below eGFP expression threshold in all per-oligonucleotide replicates. 69
S3 Mutants at lower limit of assay detection (for which reported $K_d$ is an underestimate) determined via repeated-measurement ANOVA. . . . . . . . . . . . . . . . . . . . . . 70
S4 DNA sequences for all duplexing reactions and binding affinity measurements. The E-box motif within each sequence is marked in bold. The universal 3' sequence used for annealing a 5' Alexa-647-conjugated sequence is italicized. "Core" mutation denotes mutations within E-box motif. "Consensus" sequence was used as the wildtype DNA sequence for testing specificity altering MAX mutations via double-mutant cycles. . 71
S5 Effects of MAX allelic variants on DNA-binding. . . . . . . . . . . . . . . . . . . . . . 72
S6 Non-additive mutations within measurements of the full MAX mutant library, as assesed by comparison of expected versus measured $K_d$s when bound to the cognate CACGTG motif versus to mutated E-box motifs. . . . . . . . . . . . . . . . . . . . . . 73

| Mutation Type | Mutants |
|---|---|
| Predicted pathogenic allelic variants | [D6Y, I7M, E8K, E10K, E10D, E13K, Q15R, D23H, H28R, A30P, A30T, E32K, E32N, R35C, R35L, R35H, R36W, L46S, L46W, L46F, P51S, P51L, P51Q, R60Q, R60W, I71S, M74I, R75Q, R36K, D37G, R90W, V99I] |
| Variants of unknown significance (VUS) | [D135N, R100H, M74T, Y73C, I71N, A67D, K66R, R60W, A58V, A58T, R47W, L46F, S45G, I39V, K24R, S20P, A22S, D23N, K24R] |
| Biophysical | [D5P, E10P, Q15P, D23N, R25W, A26G, H27K, A30G, E32Q, R36L, R36M, R36N, R36Q, R36Y, D41G, S42I, S42N, D48G, V50F, S52G, R60E, R60K, K66Q, Y70F, Y70H, Y73H, M74L, R75K, N78K, A21D, A21K, A22D, A22E, A22G, A22K, R25W, R35F, R35K, R75E] |
| bHLH ortholog | [D23E, K24S, R25H, R25Q, H28N, L31R, E32D, R33K, R33Q, K34Q, R35Y, R25K, H28K, L31I, L31M, L31S] |

Table S1: Classification of non-alanine and valine scanning variants measured in this study.

| E-box motif | Mutants |
|---|---|
| CACGTG | [K24R, R36Q, T124I] |
| **A**ACGTG | [K24R, R36Q, T124I] |
| C**G**CGTG | [K24R, R36Q, T124I] |
| CA**T**GTG | [T124I] |
| CACG**C**G | [K24R, R36Q, T124I] |
| CACGT**T** | [K24R, R36Q, T124I] |

Table S2: MAX mutants below eGFP expression threshold in all per-oligonucleotide replicates.

| **E-box motif** | **MAX mutants un-resolvable from background binding** |
|---|---|
| CACGTG, large MAX mutant library | [R36L, R35A, R25W, R36W, R60E, R25V, R36V, K24R, R33V, T124I, R35H, R35K, R36N, R36K, R35L, E32V, R36M, R36Q] |
| CACGTG, small mutant library | [Max˙A26G, Pho4˙E259D, Max˙A30T] |
| **A**ACGTG, large MAX mutant library | [R35F, R35H, T124I, R60E, R25H, R36K, R35A, L46A, L46F, K34V, L64V, R35V, K24R, E32K, R33V, L64A, R35K, F43V, R60W, R35C, R36Q, N29A] |
| **A**ACGTG, small mutant library | [Max˙E32N, Max˙A30T] |
| C**G**CGTG, large MAX mutant library | [R60E, R36Q, L46A, K24R, R35H, R35A, T124I] |
| C**G**CGTG, small mutant library | [Pho4˙E259D, Max˙R35V, Pho4˙WT, Max˙H27VA30G, Max˙A26G, Max˙A30T] |
| CA**T**GTG, large MAX mutant library | [R36K, T124I, L46W] |
| CA**T**GTG, small mutant library | [Max˙E32N, Max˙A30T, Max˙R35V] |
| CACG**C**G, large MAX mutant library | [L64V, L64A, R25H, T124I, R36A, R35H, R33V, R60E, R36K, R60V, V50A, R36Q, L46F, R60A, R60W, R35A, R35L, K24R] |
| CACG**C**G, small mutant library | [Pho4˙E259D, Max˙R60W, Max˙H27VA30G, Max˙A26G, Max˙A30T] |
| CACGT**T**, large MAX mutant library | [L64V, R60W, R36A, R36L, R36Q, R25H, F43A, R35A, L64A, R35V, K24R, R60A, R60V, V50A, L46A, R35F, R33V, R36K, R35H, E32K, A258G, T124I, I71S, D48G, R35L, A67V, R60E, L46F] |
| CACGT**T**, small mutant library | [Pho4˙E259D, Max˙A30GK40A, Max˙H27VA30G, Max˙R60W, Max˙A30T] |

Table S3: Mutants at lower limit of assay detection (for which reported $K_d$ is an underestimate) determined via repeated-measurement ANOVA.

| **Name** | **Sequence** |
|---|---|
| CCACGTGA cognate motif | CAATACACTGTTATC AGACC **CACGTG** ACGAG CTACTCGTTCGGTTA *TCCGGCGGTATGAC* |
| CAACGTGA core mutation | CAATACACTGTTATC AGACC **AACGTG** ACGAG CTACTCGTTCGGTTA *TCCGGCGGTATGAC* |
| CCGCGTGA core mutation | CAATACACTGTTATC AGACC **CGCGTG** ACGAG CTACTCGTTCGGTTA *TCCGGCGGTATGAC* |
| CCATGTGA core mutation | CAATACACTGTTATC AGACC **CATGTG** ACGAG CTACTCGTTCGGTTA *TCCGGCGGTATGAC* |
| CCACGCGA core mutation | CAATACACTGTTATC AGACC **CACGCG** ACGAG CTACTCGTTCGGTTA *TCCGGCGGTATGAC* |
| CCACGTTA core mutation | CAATACACTGTTATC AGACC **CACGTT** ACGAG CTACTCGTTCGGTTA *TCCGGCGGTATGAC* |
| Universal primer | AlexaFluor-647-5′-GTCATACCGCCGGA-3′ |

Table S4: DNA sequences for all duplexing reactions and binding affinity measurements. The E-box motif within each sequence is marked in bold. The universal 3' sequence used for annealing a 5' Alexa-647-conjugated sequence is italicized. "Core" mutation denotes mutations within E-box motif. "Consensus" sequence was used as the wildtype DNA sequence for testing specificity altering MAX mutations via double-mutant cycles.

| **Mutation** | **Classification** | **Effect** |
|---|---|---|
| D6Y | Pathogenic | WT-like |
| I7M | VUS | WT-like |
| E8K | Pathogenic | WT-like |
| E10D | Pathogenic | WT-like |
| E13K | Pathogenic | WT-like |
| Q15R | Pathogenic | WT-like |
| S20P | VUS | Reduces affinity for CGCGTG and CACGTT |
| A22S | VUS | Reduces affinity for CGCGTG |
| D23N | VUS | Reduces affinity for CACGTT |
| D23H | Pathogenic | Affinity-altering; decreases affinity for all E-box motifs |
| H28R | Pathogenic | Alters sequence specificity; reduces binding to all E-box motifs. |
| A30P | Pathogenic | Reduces affinity for all E-box motifs |
| A30T | Pathogenic | Selectivity-increasing |
| E32K | Pathogenic | Reduces affinity for all E-box motifs. |
| E32N | Pathogenic | Alters sequence specificity; reduces binding to CACGTG, CGCGTG, and CACGTT |
| R35C | Pathogenic | Reduces affinity for all E-box motifs |
| R35L | Pathogenic | Reduces affinity for all E-box motifs |
| R35H | Pathogenic | Reduces affinity for all E-box motifs |
| R36W | VUS | Alters sequence specificity; reduces binding to all E-box motifs |
| R36K | Pathogenic | Reduces affinity for all E-box motifs |
| D37G | Pathogenic | Reduces affinity for CACGTG |
| I39V | VUS | WT-like |
| S45G | VUS | WT-like |
| L46S | Pathogenic | Reduces affinity for all E-box motifs |
| L46W | Pathogenic | Reduces affinity for all E-box motifs |
| L46F | VUS | Reduces affinity for all E-box motifs |
| R47W | VUS | Affinity-altering; decreases affinity for all E-box motifs |
| P51S | Pathogenic | Reduces affinity for all E-box motifs |
| P51L | Pathogenic | Reduces affinity for all E-box motifs |
| P51Q | Pathogenic | Reduces affinity for all E-box motifs |
| A58V | VUS | WT-like |
| R60Q | Pathogenic | Reduces affinity for CGCGTG and CACGTT |
| R60W | VUS | Selectivity-increasing |
| K66R | VUS | WT-like |
| A67D | VUS | Reduces affinity for AACGTG, CGCGTG, CACGCG, and CACGTT |
| I71N | VUS | Affinity-altering; decreases affinity for all E-box motifs |
| I71S | Pathogenic | Reduces affinity for all E-box motifs |
| M74I | Pathogenic | WT-like |
| R75Q | Pathogenic | WT-like |
| R90W | Pathogenic | WT-like |
| V99I | Pathogenic | WT-like |
| R100H | Pathogenic | Reduces affinity for CACGTT |
| M74I | Pathogenic | WT-like |

Table S5: Effects of MAX allelic variants on DNA-binding.

| **E-box motif** | **Non-additive MAX mutants** |
|---|---|
| **A**ACGTG | [E32V, R33Q, A67D, E32Q, I71A, H28N, R36V, P51A, M74A, P51L, D37G, D41A, E32A, S42V, A30G, E32D, I71S, Q15R, D65A, D37V, H28A, I63A, H28R, I7M, D37A, R36N, L53A, L46W, R36L, R36W, H28K, Q15V, L46V, I71V, V50A, Y73H, H28V, E14V, R17V, Y70A, F43A, S42I, R36Y, R36M, P51Q, R35Y, Y70V, P51V, P51S, N78K, P16V, D41V, H27A, R33K, A30P, E13K, A67V, L46S, S59V, R75K, Y70H, R17A, R35L, R25W, I71N, E32N, H27V] |
| C**G**CGTG | [L31S, E32V, R33Q, A67D, E32Q, R35F, I71A, L31V, G55A, N78A, H28N, S20V, R36V, P51A, P16A, R25K, M74A, P51L, R35C, D37G, D41A, E32A, S42V, R60W, K24V, E32K, A30G, D5V, E32D, R25Q, I71S, Q15R, A30T, L64V, A26G, D37V, H28A, A22V, I63A, A61V, S20A, D41G, H28R, T80A, R25V, I7M, R47W, D37A, N29V, D23E, L53A, R35K, L64A, L46W, R36L, R36W, K40A, H28K, Q15V, L46V, L46F, I71V, R76A, V50A, D23V, D65V, S20P, R60Q, Y73H, H28V, E14V, R17V, Y70A, F43A, Q15P, S42I, N29A, R60V, M74V, R36Y, I63V, R36M, A21K, K24S, S59A, R47V, A26V, R35V, P51Q, A22D, R35Y, Y70V, K34V, H38V, S42N, P51V, Y73A, L31M, P51S, N78K, P16V, Q62A, D41V, H27A, K57V, H79V, H79A, V50F, R33K, R33A, R60A, A67V, L46S, S59V, R75K, Y70H, L31A, R17A, R25W, I71N, E32N, H27V] |
| CA**T**GTG | [E32V, R33Q, A67D, E32Q, R35F, I71A, G55A, D48A, R36V, P51A, P16A, M74A, P51L, R35C, D37G, D41A, E32A, S52V, T68A, E32K, A30G, H27K, E32D, D48G, Q15R, D65A, A26G, D37V, I63A, D41G, H28R, I7M, R60E, R47W, D37A, R36N, G55V, D23E, L53A, R35K, L64A, D135N, R36W, A22K, H28K, Q15V, L46V, D23A, L46F, S45G, I71V, V50A, D65V, Y73H, H28V, R17V, R90W, Y70A, F43A, S42I, R60V, M74V, S42A, R75A, R36Y, I39V, R75E, S59A, R35V, E69V, P51Q, R35Y, Y70V, K34V, P51V, P51S, N78K, D23H, P16V, Q62A, D41V, V50F, Q72V, R33K, R35H, Y73V, R100H, E13K, R60A, A67V, L46S, S59V, R75K, Y70H, F18V, R17A, R35L, K66V, I71N, I39A, E32N] |
| CACG**C**G | [L31S, E32V, R33Q, A67D, E32Q, R35F, I71A, K57A, L31V, H28N, R36V, P51A, P16A, R25K, P51L, R35C, D37G, D41A, L46A, E32A, S42V, K24V, A30G, E32D, I71S, A26G, K77V, D37V, H28A, I63A, D41G, H44V, H28R, K24A, I7M, L31I, R47W, D37A, N29V, D23E, L53A, R35K, R47A, L46W, A30V, K40A, H28K, L46V, I71V, D23V, Y73H, H28V, Y70A, F43A, S42I, N29A, R36Y, D23N, R36M, K24S, S59A, A26V, R35V, P51Q, R35Y, Y70V, K34V, P51V, P51S, N78K, P16V, Q62A, D41V, H27A, V50F, R33K, A30P, R33A, E13K, L46S, S59V, R25W, I71N, E32N, H27V, A22S] |
| CACGT**T** | [L31S, E32V, R33Q, K66Q, E13A, A67D, E32Q, I71A, K57A, L31V, G55A, N78A, H28N, R36V, D12V, P51A, P16A, R25K, D37G, M74A, P51L, R35C, D41A, E32A, S52V, S42V, A30G, H27K, S45V, D5V, E32D, D5A, R36M, Q15R, A30T, D65A, A26G, F18A, D5P, K77V, D37V, H28A, I63A, A61V, S20A, D41G, H44V, E14A, H28R, T80A, I7M, L31I, R47W, P51V, D37A, R36N, N29V, D23E, L53A, R35K, Y70F, R47A, L46W, D135N, R36W, K40A, D12A, A22K, H28K, Q15V, L46V, D23A, K40V, I71V, R76A, D23V, D65V, Y73H, H28V, E14V, R17V, R90W, D6A, Y70A, Q54A, Q15P, H38A, S42I, M74V, S42A, R75A, R36Y, I63V, D23N, R75E, A21K, S11V, K24S, S59A, A26V, K34Q, E69V, R76V, P51Q, R35Y, Y70V, K34V, Y73A, L31M, P51S, Q19A, N78K, Q54V, P16V, Q62A, D41V, H27A, K57V, H79A, V50F, Q72V, R33K, A30P, R33A, E13K, L46S, S59V, R75K, Y70H, F18V, A58V, Q72A, L31A, R17A, Q19V, R25W, I71N, A22E, I39A, E32N, H27V] |

Table S6: Non-additive mutations within measurements of the full MAX mutant library, as assesed by comparison of expected versus measured $K_d$s when bound to the cognate CACGTG motif versus to mutated E-box motifs.
